# Extracting diamonds: Identifiability of 4-node cycles in level-1 phylogenetic networks under a pseudolikelihood coalescent model

**DOI:** 10.1101/2023.10.25.564087

**Authors:** George Tiley, Nan Liu, Claudia Solís-Lemus

## Abstract

Phylogenetic networks encode a broader picture of evolution by the inclusion of reticulate processes such as hybridization, introgression or horizontal gene transfer. Each reticulation event is represented by a “hybridization cycle”. Here, we investigate the statistical identifiability of the position of the hybrid node in a 4-node hybridization cycle in a semi-directed level-1 phylogenetic network. That is, we investigate if our model is able to detect the correct placement of the hybrid node in the hybridization cycle from concordance factors as data. While generic identifiability is easily attained under non-restrictive assumptions such as *t* ∈ (0, ∞) for all branches and *γ* ∈ (0, 1) for the inheritance probability of the hybrid edges, simulations show that accurate detection of these cycles can be complicated by inadequate sampling, small sample size or gene tree estimation error. We identify practical advice for evolutionary biologists on best sampling strategies to improve the detection of this type of hybridization cycle.

## 1 Introduction

The increasing evidence of reticulate processes in the Tree of Life has inspired the development of novel mathematical and statistical methods to infer phylogenetic networks from genomic data [26, 12, 28, 29]. A phylogenetic network is an extension of the tree graph by the inclusion of hybrid nodes that allow for two incoming parental edges which represent genetic material being transferred from two different sources. Methods to infer phylogenetic networks include classes from combinatorial [19, 20, 51, 36, 47], distance-based [49, 35, 50], or model-based under Markov models that use genetic sequences or markers as input [18, 40, 33] or under the multispecies coalescent model [41] that uses gene trees as input [54, 44, 56].

Model-based approaches provide accurate representations of the reticulate evolutionary process among species represented by the estimated phylogenetic network. However, the accurate estimation of this parameter requires proof of identifiability. Indeed, multiple studies have tackled the question of whether phylogenetic networks are identifiable under different models [37, 44, 17, 32, 7, 3, 46, 18, 50, 2], and here, we extend the identifiability proofs in [44, 46] to further validate the inference of level-1 phylogenetic networks under a pseudolikelihood model built on the multispecies network coalescent model. Since identifiability can be unattainable in many cases, in practice, one aims to prove *generic identifiability* which means that the network parameter is almost surely identifiable.

Let *N* be a semi-directed level-1 phylogenetic network with *h* hybridization events and *n* taxa. We denote this the *network parameter* of the model. Let *G* = {*G*_1_*, G*_2_*, …, G_g_*} be *g* estimated gene trees as input for the model. We can define a likelihood as the product of the probability under the multispecies network coalescent model [53] of each gene tree given the network parameter (with its numerical parameters as branch lengths and inheritance probabilities). For the case of a species tree as the model parameter, it has been proven that the distribution of gene trees identifies (or generically identifies) the tree topology, branch lengths and even root position under the multispecies tree coalescent model [11, 6, 5, 42]. The same is not known for the full likelihood model on species networks, but generic identifiability proofs have been derived for the case of a quartet-based pseudolikelihood model [44, 7, 46] that assumes probabilities of each observed quartet gene tree under its corresponding species quarnet are independent. These results show that the presence of hybridization events in *n*-taxon level-1 semi-directed phylogenetic networks is generically identifiable from the observed distribution of gene trees under a pseudolikelihood model.

Here, we begin to explore the question of network identifiability by focusing on level-1 phylogenetic networks with hybridization cycles of 4 nodes (4-node cycles). These types of cycles (denoted *diamonds* [39]) are relevant because in practice they seem to lead to flat pseudolikelihood scores [44]. That is, 4-node cycles that only differ on the placement of the hybrid node in the cycle can have similar pseudolikelihood scores. Here, we show that a semi-directed network (correct direction of hybrid edges) is generically identifiable from input unrooted gene trees, not only the unrooted network (placement of the cycle). This implies that the direction of the gene flow event (hybrid edge) can be distinguished with the pseudolikelihood model, providing an accurate evolutionary interpretation of the origin and target of reticulations. Our proof follows the same algebraic geometry techniques as in [44, 46] such as the definition of the set of polynomial equations under the coalescent model and the search for unique (or finitely many) solutions.

The organization of the manuscript is as follows. In Section 2, we establish the models more precisely and the approach used to prove generic identifiability. In Section 3, we define the theorems and proofs for the generic identifiability of the placement of the hybrid node in the 4-node hybridization cycle. In Sections 4 and 5, we illustrate the mathematical findings (and the practical considerations when dealing with limited data) with a simulation study. Finally, we present some discussion in Section 6.

## 2 Model Preliminaries

### 2.1 Semi-directed level-1 explicit phylogenetic networks

Our main parameter of interest is the topology *N* of a phylogenetic network (Definition 1) along with the numerical parameters of the vector of branch lengths (***t***) and a vector of inheritance probabilities (***γ***). The inheritance probabilities describe the proportion of genes inherited by a hybrid node from one of its hybrid parent (see Figure 1).

**Figure 1:**
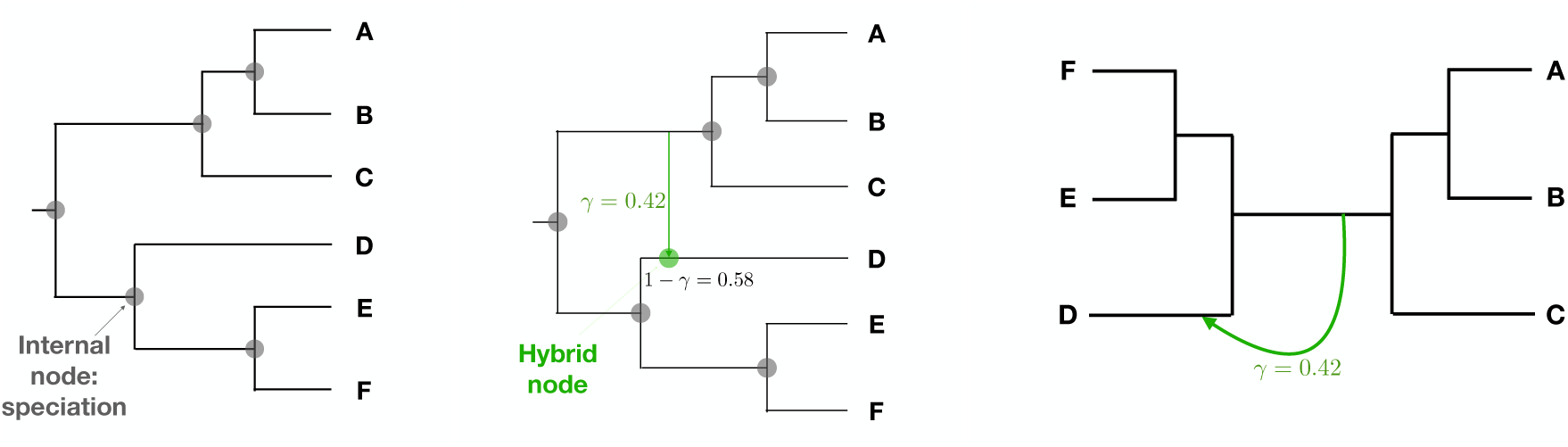
Left: Rooted phylogenetic tree of 6 taxa. Internal nodes (gray circles) represent speciation events and are usually omitted on phylogenetic trees. Center: Rooted phylogenetic network on 6 taxa with *h* = 1 hybridization event represented by the green edge. The hybrid node (in green) represents a reticulation event. The hybrid node has two parent edges: minor hybrid edge in green labeled *γ* = 0.42 and major hybrid edge in black labeled 1 − *γ* = 0.58. The reticulation event can represent different biological processes: hybridization, horizontal gene transfer or introgression. Right: Semi-directed network for the same biological scenario in center. Although the root location is unknown, its position is constrained by the direction of the hybrid edges. For example, the taxon D cannot be an outgroup.

**Definition 1.** A rooted explicit phylogenetic network [27] N *on taxon set X is a connected directed acyclic graph with vertices V* = {*r*} ∪ *V_L_* ∪ *V_H_*∪ *V_T_, edges E* = *E_H_* ∪ *E_T_ and a bijective leaf-labeling function f* : *V_L_* → *X with the following characteristics: i) The root r has indegree 0 and outdegree 2, ii) Any leaf v* ∈ *V_L_ has indegree 1 and outdegree 0, iii) Any tree node v* ∈ *V_T_ has indegree 1 and outdegree 2, iv) Any hybrid node v* ∈ *V_H_ has indegree 2 and outdegree 1, v) A tree edge e* ∈ *E_T_ is an edge whose child is a tree node, vi) A hybrid edge e* ∈ *E_H_ is an edge whose child is a hybrid node, vii) A hybrid edge e* ∈ *E_H_ has an inheritance probability parameter γ_e_ <* 1 *which represents the proportion of the genetic material that the child hybrid node received from this parent edge. For a tree edge e, γ_e_* = 1.

In a rooted explicit network, every internal node represents a biological process: speciation for tree nodes and hybridization for hybrid nodes. However, other types of phylogenetic networks, such as unrooted networks [27] and semi-directed networks [44, 18] are also useful representation of evolutionary relationships. Unrooted phylogenetic networks are typically obtained by suppressing the root node and the direction of all edges. In semi-directed networks, on the other hand, the root node is suppressed and we ignore the direction of all tree edges, but we maintain the direction of hybrid edges, thus keeping information on which nodes are hybrids. The placement of the root is then constrained, because the direction of the two hybrid edges to a given hybrid node inform the direction of time at this node: the third edge must be a tree edge directed away from the hybrid node and leading to all the hybrid’s descendants. Therefore the root cannot be placed on any descendant of any hybrid node, although it might be placed on some hybrid edges. See Figure 1 for the example of a rooted explicit phylogenetic network (center), and its semi-directed version (right).

Each hybridization event creates a blob (or cycle) which represents a subgraph with at least two nodes and no cut-edges. A cut-edge is any edge in the network whose removal disconnects the network. Traditionally, the term *cycle* is used for directed (rooted) networks, and the term *blob* is used for undirected (unrooted) networks. We make no distinction here as we will focus on *semi-directed* networks, and thus, some edges in the cycle (blob) will be directed, and some will not. We also note that for every hybridization event, there are two parent hybrid edges connected to the hybrid node: 1) major hybrid edge with inheritance probability *γ >* 0.5, and 2) a minor hybrid edge with inheritance probability *γ <* 0.5. Both edges are parametrized with the same *γ*.

Here, we focus on the case of semi-directed networks. In addition, we assume the following characteristics of the semi-directed network under study.

**Assumption 1.** Let N *have n species and h hybridization events (that is,* |*V_L_*| = *n and* |*V_h_*| = *h)*.

**Assumption 2.** We further assume that N *is of level-1* [27]*. That is, we assume that any given edge can be part of at most one cycle which means that there is no overlap between any two cycles (*[27, 43].

Thus, our parameters of interest are (*N,* ***t***, ***γ***) where *N* is an explicit semi-directed level-1 phylogenetic network that links the *n* species under study, and has *h* hybridization events. This network has two vectors of numerical parameters: 1) branch lengths ***t*** ∈ [0, ∞)*^ne^* for *n_e_* branches in the network, and 2) inheritance probabilities ***γ*** ∈ [0, 1]*^nh^* for *n_h_* minor hybrid edges.

### 2.2 Generic identifiability of the position of the hybrid node

To show that the position of the hybrid node in the hybridization cycle is generically identifiable, we represent every semi-directed level-1 network *N* as a set of polynomial concordance factor (CF) equations as in [44, 46], and we find if both systems of polynomial equations share solutions in the parameter space: *CF* (*N, **t**, **γ***) = *CF* (*N*^′^*, **t***^′^*, **γ***^′^).

First, each network can be decomposed into 4-taxon subnetworks (*quarnets* in [23]). That is, for a given network *N* with *n* ≥ 4 taxa, we consider all 4-taxon subsets S = {*s* = {*a, b, c, d*} : *a, b, c, d* ∈ *X*} to define the theoretical CFs expected under the coalescent model for each 4-taxon subset. These theoretical CFs are already derived for a species tree in [4], and for a species network in [44]. In both cases, the CFs do not depend on the position of the root. For the tree, the major CF is defined for the quartet that agrees with the species tree. That is, if the species tree has the split *ab*|*cd* with internal edge *t*, then the major CF would be *CF_ab_*_|_*_cd_* = 1 − 2/3 exp(−*t*). The CF for the minor resolutions (in disagreement with the species tree *ab*|*cd*) would then be *CF_ac_*_|_*_bd_* = *CF_ad_*_|_*_bc_* = 1/3 exp(−*t*) [24].

For the case of a 4-taxon network, the theoretical CFs are weighted averages of CFs on trees. For example, Figure 2 (center) shows a semi-directed 4-taxon network and the two displayed quartet trees depending on which hybrid edge is used (major hybrid edge used corresponds to the quartet on top, and minor hybrid edge used corresponds to the quartet on the bottom). The CFs on this 4-taxon network are then given by the weighted averages of CF of the two displayed quartet trees with weights: 1 − *γ* and *γ*. Namely,

- *CF_ab_*_|_*_cd_* = (1 − *γ*)(1 − 2/3 exp(−*t*_13_ − *t*_23_ − *t*_2_)) + *γ*(1/3 exp(−*t*_2_)) for the major resolution, and
- *CF_ac_*_|_*_bd_* = *CF_ad_*_|_*_bc_* = (1 − *γ*)(1/3 exp(−*t*_13_ − *t*_23_ − *t*_2_)) + *γ*(1/3 exp(−*t*_2_)) for the minor resolutions.

**Figure 2:**
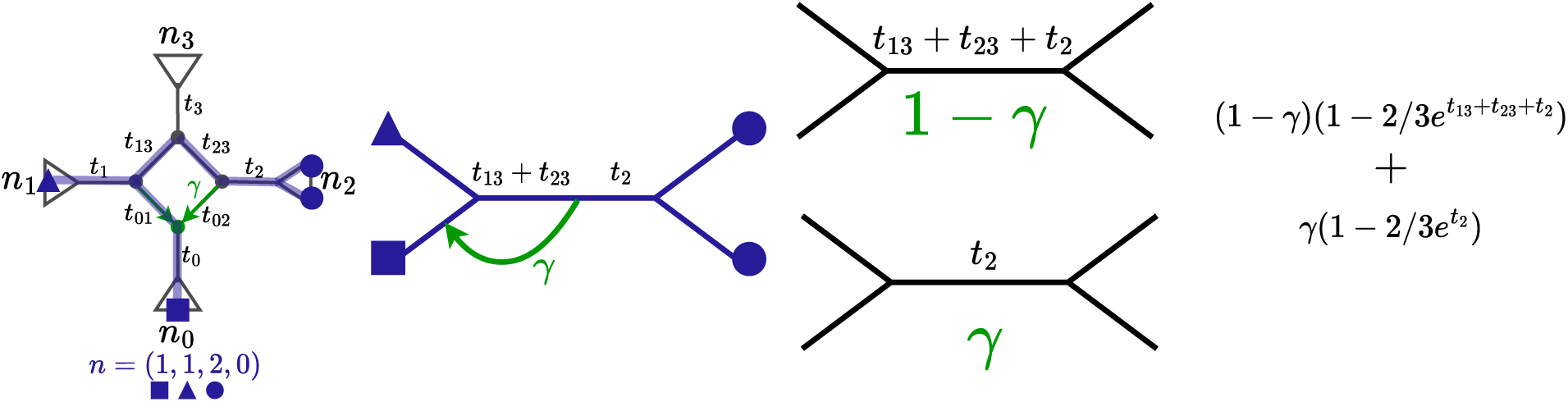
Left: Semi-directed network with a given 4-taxon subset (1, 1, 2, 0) highlighted in blue which corresponds to taking one individual in *n*_0_ (red square), one individual in *n*_1_ (red triangle) and two individuals in *n*_2_ (red circles). To obtain the CF equations for the resulting quarnet (center in blue), we split it as two quartet trees weighted by 1 − *γ* and *γ* (center in black). Right: Major CF equation for the quarnet as weighted average of CF equations on quartet trees. Because the two quartet trees have the same topology, the two minor CF equations of the quarnet are equal (not shown).

We show in the Appendix the system of polynomial equations for the four phylogenetic networks considered in this study: *N_down_*, *N_left_*, *N_right_*, and *N_up_* (Figure 3). Since each network is obtained by the rotation of the hybrid node in the hybridization cycle, there is a simple mapping of 4-taxon subsets that allows us to define the CF equations for all networks from the original *N_down_*. This process is also described in the Appendix.

**Figure 3:**
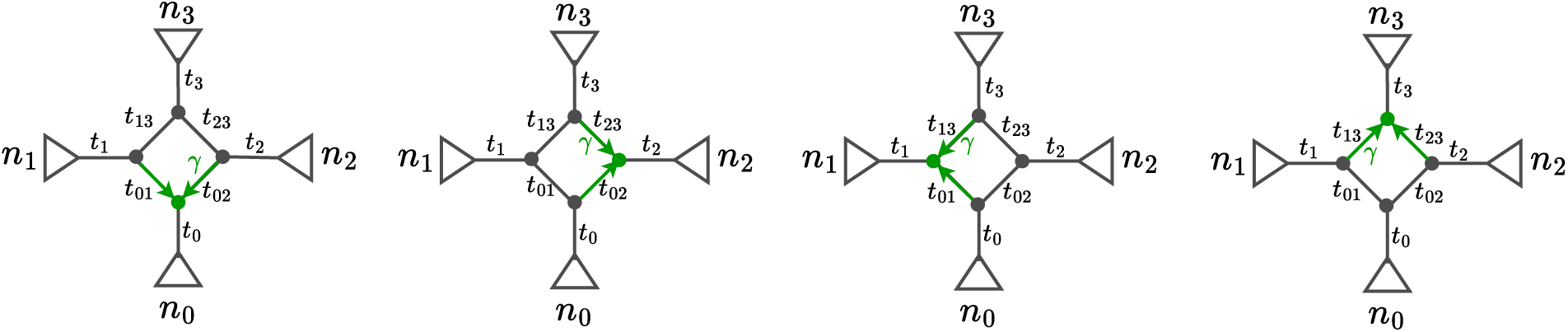
Semi-directed networks with one hybridization cycle with 4 nodes (diamond), but different position of the hybrid node in the cycle. We denote these networks *N_down_*, *N_right_*, *N_left_*, and *N_up_* respectively from left to right.

**Definition 2.** Let N *be n-taxon semi-directed level-1 explicit phylogenetic network with h hybridizations. This network defines a set of* 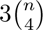 *CF equations under the coalescent model with parameters* ***t*** *and* ***γ****. Denote this system of equations as CF* (*N,* ***t***, ***γ***)*. If we change the variables to z_i_*= exp (−*t_i_*) *for all internal branch lengths, then CF* (*N,* ***z***, ***γ***) *is a system of polynomial equations*.

**Definition 3** (Generic identifiability of hybrid node position). Let N *be an n-taxon semi-directed level-1 explicit phylogenetic network with h hybridizations. We focus on one hybridization that has 4 nodes in the hybridization cycle. Let N*^′^ *be a network with the hybrid node rotated inside the 4-node hybridization cycle. Let CF* (*N,* ***t***, ***γ***) *be the system of polynomial CF equations defined by N, and let CF* (*N*^′^, ***t***^′^, ***γ***^′^) *be the system of polynomial CF equations defined by N*^′^*. We say the position of the hybrid node in the 4-node hybridization cycle in N is **identifiable** if the system of CF* (*N,* ***t***, ***γ***) = *CF* (*N*^′^, ***t***^′^, ***γ***^′^) *does not have solutions in any set of numerical parameters* (***t***, ***γ***, ***t^′^***, ***γ^′^***)*. We say the position of the hybrid node in the 4-node hybridization cycle in N is **generically identifiable** if the solution set of the system CF* (*N,* ***t***, ***γ***) = *CF* (*N*^′^, ***t***^′^, ***γ***^′^) *has measure zero*.

## 3 Identifiability of 4-node cycles in semi-directed level-1 phylogenetic networks

In this paper, we prove the following theorem regarding the generic identifiability of the placement of the hybrid node in the 4-node hybridization cycle in a level-1 network:

**Theorem 1.** Let N *be a semi-directed level-1 n-taxon phylogenetic network with one hybridization event that creates a 4-node hybridization cycle. Then, the placement of the hybrid node in the cycle is generically identifiable if i) n >* 5*, ii) t* ∈ (0, ∞) *for all branch lengths, and iii) γ* ∈ (0, 1) *for the inheritance probability corresponding to the hybridization event*.

To prove this theorem, we prove three separate theorems: *N_down_* versus *N_right_* (Theorem 2), *N_down_* versus *N_left_* (Theorem 3), and *N_down_* versus *N_up_* (Theorem 4). In this section, we show Theorem 2 and its proof, and the other two theorems and proofs are placed in the Appendix. For these three theorems, we assume the networks only have six taxa and one hybridization event. We generalize to *n* taxa and *h* hybridization events in Remark 2.

**Theorem 2.** Let N_down_ be a semi-directed level-1 6-taxon phylogenetic network with one hybridization event producing a hybridization cycle with 4 nodes. Without loss of generality, let the taxa be partitioned among clades as n_0_ = 1*, n*_1_ = 2*, n*_2_ = 1*, n*_3_ = 2 *(Figure 3). Let the hybrid node be ancestral to the clade n*_0_*. Let N_right_ be a semi-directed level-1 6-taxon phylogenetic network with one hybridization event producing a hybridization cycle with 4 nodes such that the unrooted version of N_right_ agrees with the unrooted version of N_down_. Let the hybrid node in the hybridization cycle in N_right_ be ancestral to the clade n*_2_*. Then, N_down_ and N_right_ are identifiable if t*_1_ *<* ∞*, t*_13_ *>* 0*, t*_3_ *<* ∞*, and γ* ∈ (0, 1).

*Proof.* Let *CF* (*N_down_,* ***z***, ***γ***) be the system of CF polynomial equations defined by *N_down_* and let *CF* (*N_right_,* ***z***^′^, ***γ***^′^) be the system of CF polynomial equations defined by *N_right_*. Both systems of equations can be found in the Appendix.

Let P = {*p*(***z****, **γ***) − *q*(***z***^′^*, **γ***^′^) : *p*(***z****, **γ***) ∈ *CF* (*N_down_, **z**, **γ***)*, q*(***z***^′^*, **γ***^′^) ∈ *CF* (*N_right_, **z***^′^*, **γ***^′^)} be the set of polynomial equations resulting from matching *CF* (*N_down_, **z**, **γ***) to *CF* (*N_right_, **z***^′^*, **γ***^′^) for every 4-taxon subset.

Using Macaulay2 [15], we compute the Gröbner basis of P on the (***z***, ***γ***) variables by any elimination order. All Macaulay2 scripts are available at https://github.com/gtiley/diamoNd-ideNtifiability.

The resulting ideal is given by

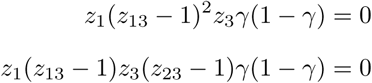

which represent the conditions that the (***z****, **γ***) variables need to satisfy for the polynomial set P to vanish to zero.

Thus, *N_down_*and *N_right_*are not identifiable in the subset of parameter space corresponding to {*z*_1_ = 0} ∪ {*z*_13_ = 1} ∪ {*z*_3_ = 0} ∪ {*γ* = 0} ∪ {*γ* = 1}.

**Remark 1.** We note that by assuming that t ∈ (0, ∞) *for all branch lengths, and γ* ∈ (0, 1) *for the inheritance probabilities, we can guarantee generic identifiability of the placement of the hybrid node in the 4-node hybridization cycle*.

**Remark 2.** The identifiability of the position of the hybrid node in 4-node cycles in n-taxon level-1 phylogenetic networks is obtained by noticing that we need at most 2 taxa per clade to define all the CF polynomial equations (Lemma 1 in [46]), and thus, if the hybridization cycles are identifiable with only one taxon in some clades, the addition of a second taxon will only reduce the set in the parameter space where the two networks are not (generically) identifiable.

## 4 Simulations

### 4.1 Simulations without gene tree estimation error

We simulated gene trees under the multispecies network coalescent (MNSC) using BPP v4.1.4 [14]. The level-1 network under investigation contains eight taxa and one 4-node hybridization cycle (diamond) (Figure 4a). To simulate gene trees under the MNSC, we require a rooted evolutionary history, and thus, we investigate three potential root placements: 1) a balanced root included in the cycle (*r*_1_ in Figure 4b), 2) a ladderized root with the root in the cycle (*r*_2_ in Figure 4c), and 3) a ladderized root such that the cycle occurs in the ingroup (*r*_3_ in Figure 4d). The root height is not the same between the balanced root and ladderized rootings, but the age of episodic gene flow is 1 ^1^ units of *θ*, population size measured in expected nucleotide diversity. The inheritance probability (*γ*) was 0.5 for all simulations. Simulations were carried out for nine diamonds for the three rooting scenarios, such that we changed if lineages were represented by a clade of two taxa or a single taxon (Figure 4e).

**Figure 4:**
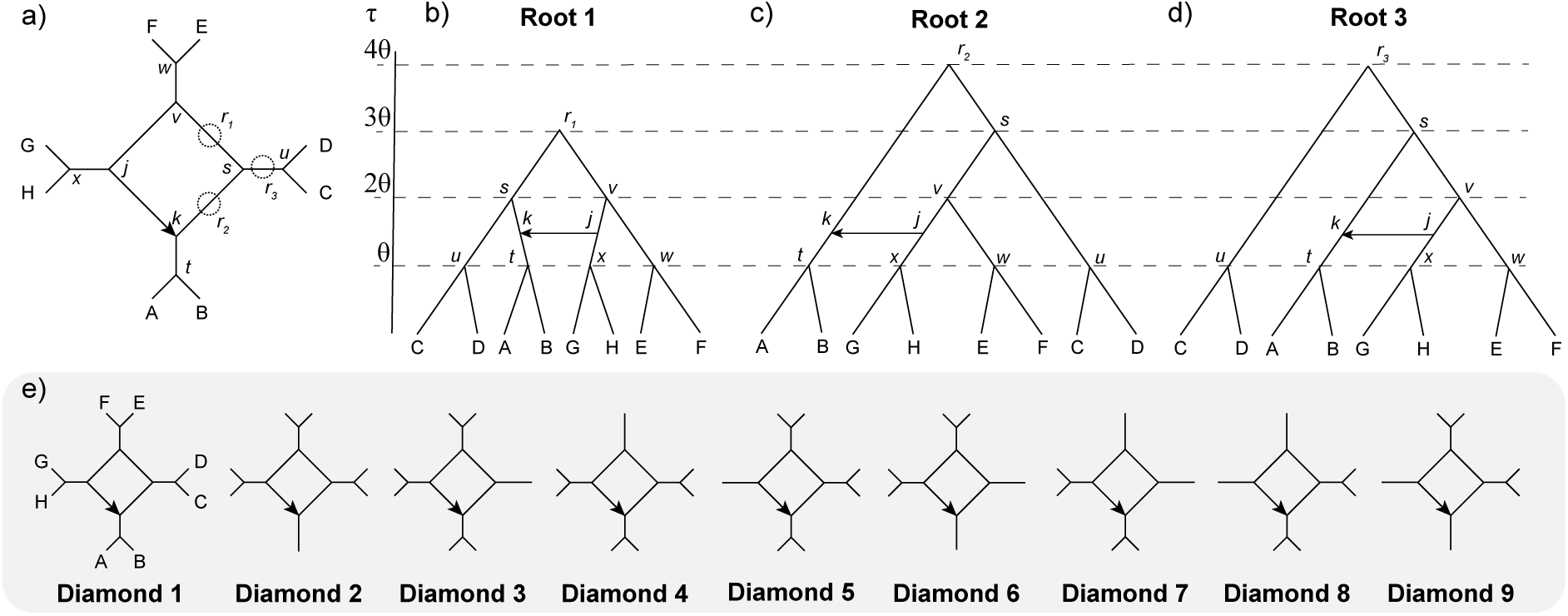
Semi-directed level-1 network topology with a diamond hybridization cycle. a) The semi-directed diamond has tip labels in capital letters and node labels in lower case italicized letters. The hybrid node is labeled as *k*. The location of the three possible roots are indicated by *r*_1_, *r*_2_, *r*_3_. b-d) Node heights for the three rooted networks in units of *θ* population size. The direction of episodic gene flow is shown with an arrow. e) For each root, nine networks were investigated, the first including all tips, the next four reducing one cherry to a single branch in an anticlockwise fashion, the next four reducing two cherries to two lineages with a single taxon.

Parameters for sequence and gene tree simulation under the MNSC was a *θ* of 0.01 constant over all nodes. The HKY model of nucleotide substitution [21] with a transition-transversion rate-ratio of 3 and equilibrium frequencies at *π_A_* = *π_T_* = 0.3 and *π_C_* = *π_G_* = 0.2 was selected. A low degree of among site rate variation was incorporated using a gamma distribution [52] with a shape and rate of 0.6. A strict clock was used among branches and there is no among-locus rate variation. Simulated sequences have 1000 base pairs. One-hundred simulations were performed for each of the 27 rooted networks, using 100, 500, 1000, and 5000 loci.

We then selected the ladderized root (root 3 in Figure 4) to assess the changes in performance due to the value of the inheritance parameter. We varied the number of gene trees and the inheritance rate across nine diamonds, as illustrated in Figure 4e. Specifically, we examined nine parameter combinations using the following values: 250, 1000, and 4000 gene trees, paired with inheritance rate of 0.05, 0.25, and 0.5. For each diamond and parameter combination, we performed 100 replicate simulations. Scripts for reproducing simulation experiments are available at https://github.com/gtiley/diamoNd-ideNtifiability.

### 4.2 Incorporating gene tree error into simulation experiments

Biological data are often messy with many potential sources of gene tree estimation error. For example, deep paralogy can lead to gene trees that conflict with species trees [34] and lead to rooting errors when using an outgroup [22, 25]. Such gene tree errors can be prevalent in groups with histories of large-scale gene duplication and loss such as plants [30] and insects [31]. These rooting errors may negatively effect the performance of inference methods, especially those based on rooted triples. Therefore, we studied two types of gene tree estimation error: 1) for each simulation condition, we randomly re-rooted 10% and 30% of the gene trees. Because rooting errors may also be accompanied by other topological errors, we estimated networks from the simulated data incorporating a random re-rooting and a random nearest neighbor interchange (NNI) move to 10% and 30% of gene trees. 2) We estimated gene trees under different levels of noise as described next.

In the second stage, based on the initial simulation results in Section 4.1, we identified five diamonds (*d*_1_*, d*_2_*, d*_3_*, d*_8_*, d*_9_) that exhibited representative performance for all diamonds, and these were further investigated under gene tree estimation error. For this simulation, we fixed the number of gene trees at 1000 and the inheritance rate at 0.25, while varying the sequence length across 250, 1000, and 4000 for each of the five selected diamonds. We utilized IQ-TREE v2.3.6 [9] to estimate the gene trees. Every scenario was replicated 20 times. Scripts for introducing random roots and NNI moves to trees are available at https://github.com/gtiley/diamoNd-ideNtifiability.

### 4.3 Network estimation

Networks were estimated from the simulated or estimated gene trees using pseudolikelihood methods that utilize the gene trees topology only. We used two methods: 1) the SNaQ function [44] implemented through the PhyloNetworks v0.15.0 Julia package [45] in Julia v1.6.5 and 2) the I*N*fer*N*etwork_MPL function [55] implemented through PhyloNet v3.8.3 [48]. Notably, SNaQ uses unrooted quartets as data while I*N*fer*N*etwork_MPL uses rooted triples (more on rooting error later). Both analyses used ten independent runs with the maximum number of allowed reticulation events set to zero, one, or two.

Three summary statistics were used to evaluate the performance of the estimations. First, we calculated the proportion of simulations where only one reticulation was detected based on a two-point pseudolikehood score difference. Although the pseudolikelihood does not have straight-forward implementation of model selection such as the Akaike Information Criterion [1], we used differences in pseudolikelihood scores as some operational criteria to evaluate detection of reticulation edges among the replicates in lieu of slope heuristics [8] or goodness-of-fit tests [10] that might be more appropriate for an empirical investigation. Second, we verified if the estimated network had the same topology as the true network using the hardwiredClusterDista*N*ce function [27] in PhyloNetworks. While the absolute distance between networks is difficult to interpret, a distance of 0 means the network topologies are identical. Third, we checked if the true minor hybrid edge is present in the estimated network with only one estimated hybridization event. This is done by drawing bootstrap support from the estimated network onto the true network with the hybridBootstrapSupport function from PhyloNetworks. A bootstrap support of 100% shows that the true hybrid edge is present in the estimated network, even if some other aspects of the network are incorrect. Scripts for calculating these summary statistics are available at https://github.com/gtiley/diamoNd-ideNtifiability.

## 5 Results

### 5.1 Simulation results: Error-free gene trees

The pseudolikelihood methods implemented in PhyloNetworks and PhyloNet generally perform well across all simulation conditions. SNaQ is capable of correctly identifying the presence of one reticulation event, recovering the correct network, and recovering the reticulation node in all simulations as the number of gene trees increased (Supplementary Figures 11, 12, 13). For diamonds *d*_6_ and *d*_9_ when the rooting is ladderized and in the cycle (*r*_2_) or ladderized and out of the cycle (*r*_3_), SNaQ sometimes missed the correct network or reticulation node when the number of gene trees was low, but converged to the correct network as gene trees increased from 100 to 5000. I*N*fer*N*etwork_MPL was more efficient with respect to the number of gene trees for *d*_6_ or *d*_9_ and performed well for *r*_1_ (Supplementary Figure 14) and *r*_2_ (Supplementary Figure 15), but sometimes struggled with *r*_3_ (Supplementary Figure 16). Across all diamonds for *r*_3_, I*N*fer*N*etwork_MPL did not always detect one reticulation edge, such that two reticulation edges were preferred over none (Supplementary Data). Even when considering the estimated networks that only allowed one reticulation event, the network was not always correct; although, the correct reticulation node is almost always recovered (Supplementary Figure 16). If the recovery of the reticulation node is the most favorable criteria of an estimator, both SNaQ and I*N*fer*N*etwork_MPL perform well with error free gene trees, with SNaQ requiring more gene trees in cases where the the hybrid node had a single taxon for a descendant instead of a cherry (Figure 5).

**Figure 5:**
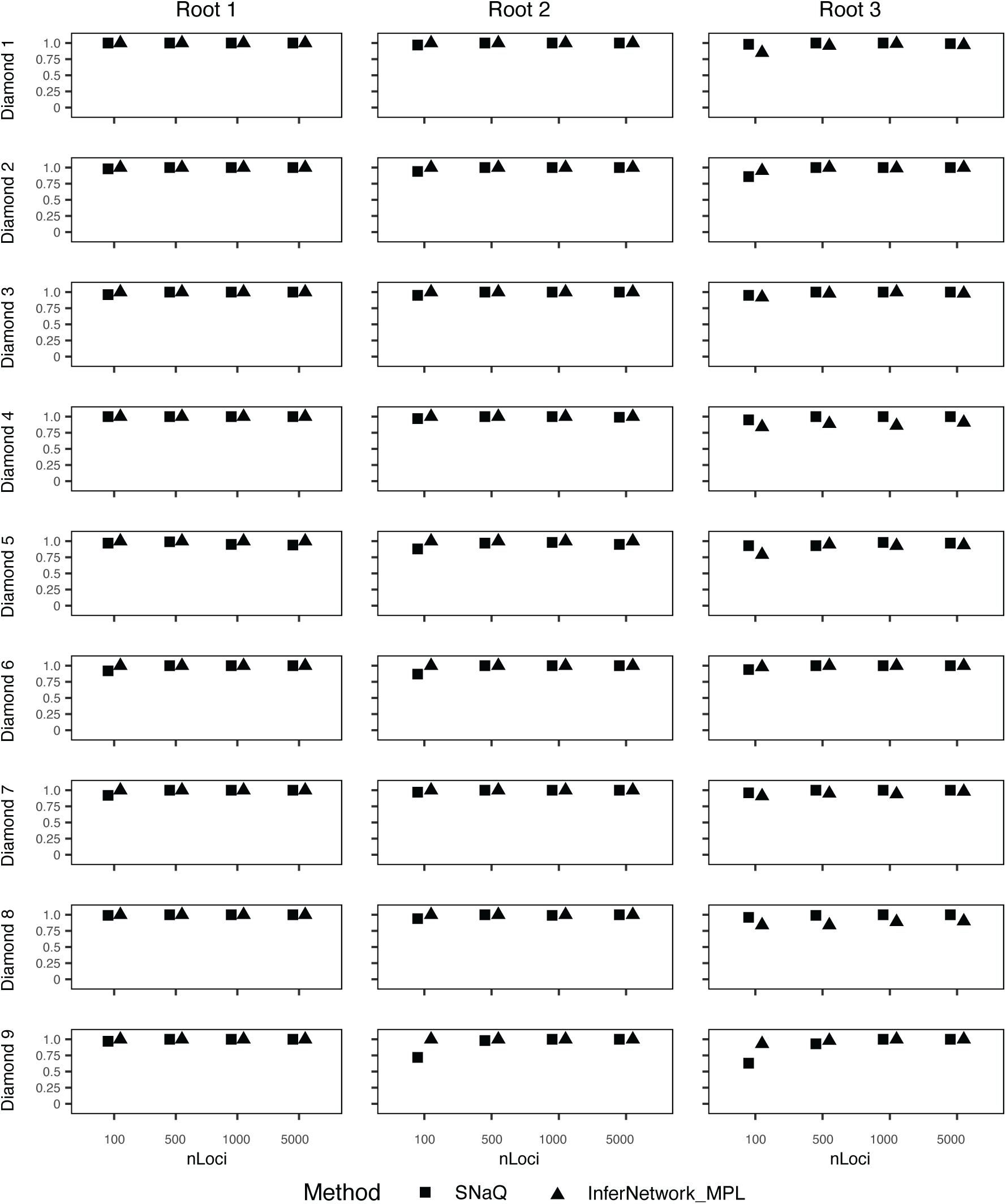
Proportion of 100 simulation replicates where the true reticulation edge is recovered by SNaQ (squares) or I*N*fer*N*etwork_MPL (triangles). X-axis corresponds to the number of gene trees in the input sample. The diamond and root numbers correspond to Figure 4.

When the inheritance rate is 0.25 and 0.5, SNaQ is capable of identifying the presence of one reticulation event, recovering the reticulation node, and recovering the correct network when the number of gene trees increased (Figure 6). For diamond *d*_5_, *d*_6_ and *d*_9_, the correct network or reticulation node was sometimes missed when the number of gene trees was low (250), but converged to the correct network as the gene trees increased to 1000 and 4000. When the inheritance rate is 0.05, recovering the correct network and recovering the reticulation node had an increasing trend across all diamonds while the number of gene trees increased. Diamond *d*_9_ could only be recovered about 50 percent even with 4000 gene trees. Furthermore, SNaQ could identify the presence of the one reticulation event for all diamonds excluding *d*_6_ and *d*_9_, whose presence rate is decreasing while the number of gene trees increased. Since *d*_6_ and *d*_9_ are symmetric, their results both reflected false negative when including more gene trees.

**Figure 6:**
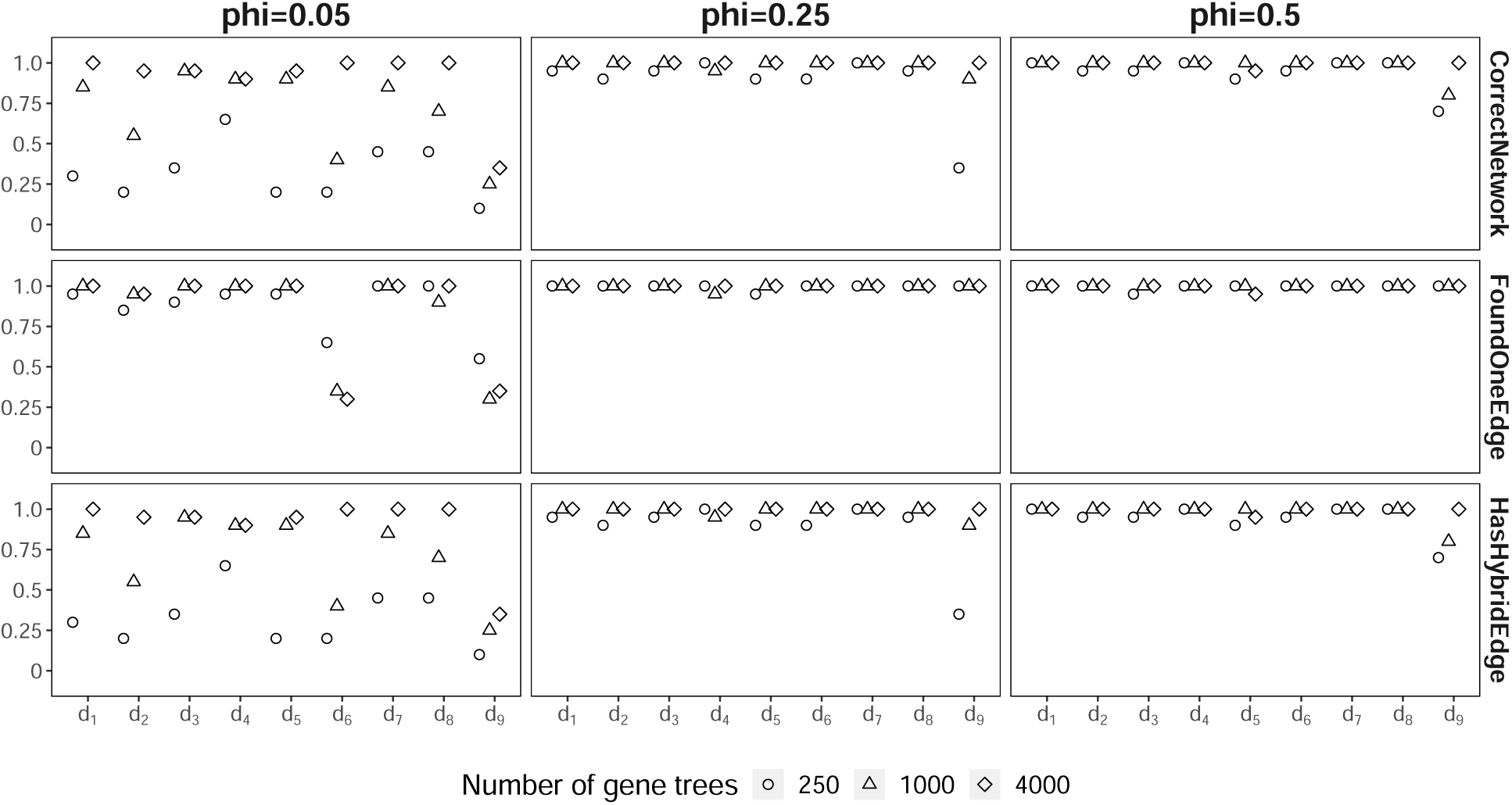
SNaQ results for root 3 with different value of the number of gene trees paired with the inheritance rate. *One Edge Detected* is the proportion out of 20 replicates that 1 reticulate edge was correctly inferred by the pseudolikehood scores. *Network is Correct* is the proportion of 20 replicates where the estimated topology when allowing only one reticulation event is identical to the true topology. *Hybrid Edge Recovered* is the proportion of 20 replicates where the correct reticulation edge was inferred, regardless if other parts of the estimated network were incorrect, when allowing only one reticulation event. Phi represents the inheritance rate. Diamond *d*_1_ through *d*_9_ correspond to Figure 4e.

### 5.2 Simulation results: Gene trees with error

SNaQ tolerated rooting errors well when they were present in both 10% (Supplementary Figures 17, 18, 19) and 30% (Supplementary Figures 20, 21, 22) of gene trees. The patterns for *d*_6_ and *d*_9_ with *r*_2_ and *r*_3_ from error-free gene trees were observed again, which was expected since SNaQ used unrooted quartets as data. I*N*fer*N*etwork_MPL was largely robust to root error in 10% of the gene trees for *r*_1_ and *r*_2_, with the exception of sometimes detecting more than one reticulation edge (Supplementary Figures 23 and 24). However, it was difficult to recover the correct network for *r*_3_ and the proportion of incorrect networks sometimes increased with the number of gene trees (Supplementary Figure 25). The lowest proportions of correct network estimations are observed for diamonds *d*_4_, *d*_7_, and *d*_9_, which are not all cases where the hybrid node has a single taxon as a descendant. The overall network being incorrect does not always mean that the estimated reticulation node is wrong, but the decreased performance compared to simulations without gene tree error was drastic (Supplementary Material Table 15). As the proportion of gene tree errors increased from 10% to 30%, similar patterns were observed for I*N*fer*N*etwork_MPL, but with a higher proportion of network estimation errors (Supplementary Figure 26, 27, 28). For example, the reticulation node was never recovered for *d*_9_ with *r*_3_ and 5000 gene trees (Supplementary Figure 28).

Introducing an NNI move to the gene trees with random roots had little effect on SNaQ’s ability to correctly detect one reticulation node, estimate the correct network, and recover the reticulation node. SNaQ nearly converges to the correct answer in all simulation conditions as the number of gene trees increased, regardless of whether errors were present in 10% (Supplementary Figures 29, 30, 31) or 30% (Supplementary Figures 32, 33, 34) of gene trees.

The NNI error did not greatly affect I*N*fer*N*etwork_MPL compared to the rooting error alone. I*N*fer*N*etwork_MPL is robust to gene tree error with wrong roots and the NNI move in 10% of gene trees for *r*_1_ and *r*_2_ (Supplementary Figures 35, 36). There are many estimation errors for *r*_3_, with the reticulation node detected about half of the time with 5000 gene trees for *d*_4_ and *d*_9_; however, the reticulation node is almost always recovered despite the estimated network being wrong for *d*_6_ (Supplementary Figure 37). When 30% of the gene trees had rooting and NNI errors, I*N*fer*N*etwork_MPL still performed well for *r*_1_ and *r*_2_ while the proportion of incorrect estimates increaseed for *r*_3_ (Supplementary Figures 38, 39, 40). Even with the more conservative assumption of 10% gene tree error, SNaQ was more robust than I*N*fer*N*etwork_MPL, at least when errors cause rooting problems (Figure 7). The estimation performance was overall sensitive to the root placement of the simulated network and sampling of a single taxon versus cherries as hybrid node descendants despite the diamond remaining constant across simulations.

**Figure 7:**
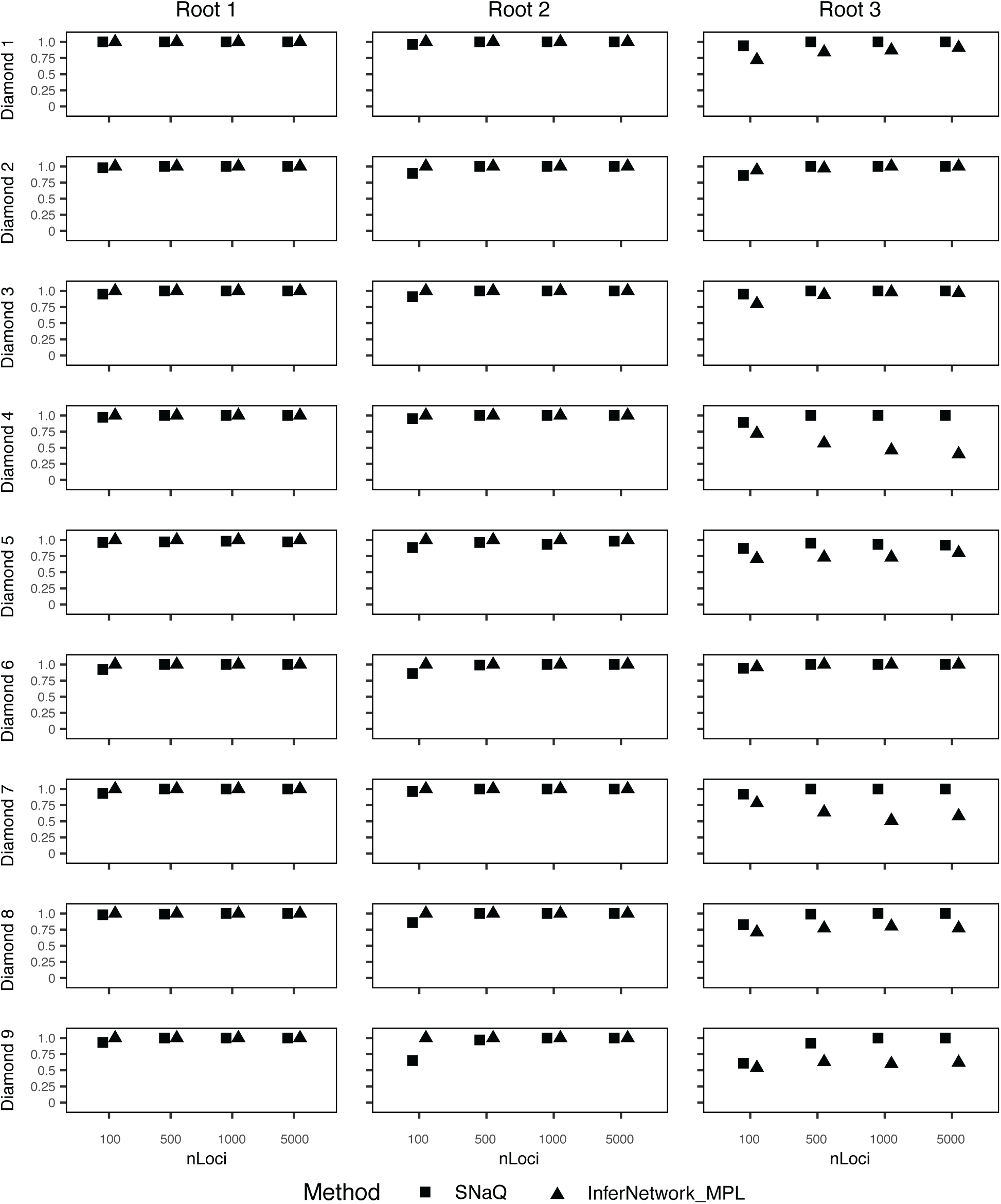
Proportion of 100 simulation replicates where the reticulation node was recovered by SNaQ (squares) or I*N*fer*N*etwork_MPL (triangles). X-axis corresponds to the number of gene trees in the input sample. The diamond and root numbers correspond to Figure 4. Error was introduced into 10% of gene trees for each round of simulation by randomly rerooting the gene tree and performing a random NNI move.

When utilizing estimated gene trees, accurate estimation improves as sequence length increases as is expected (Figure 8). Only d9 performed a different trend as the fraction dropped for large sequence length.

**Figure 8:**
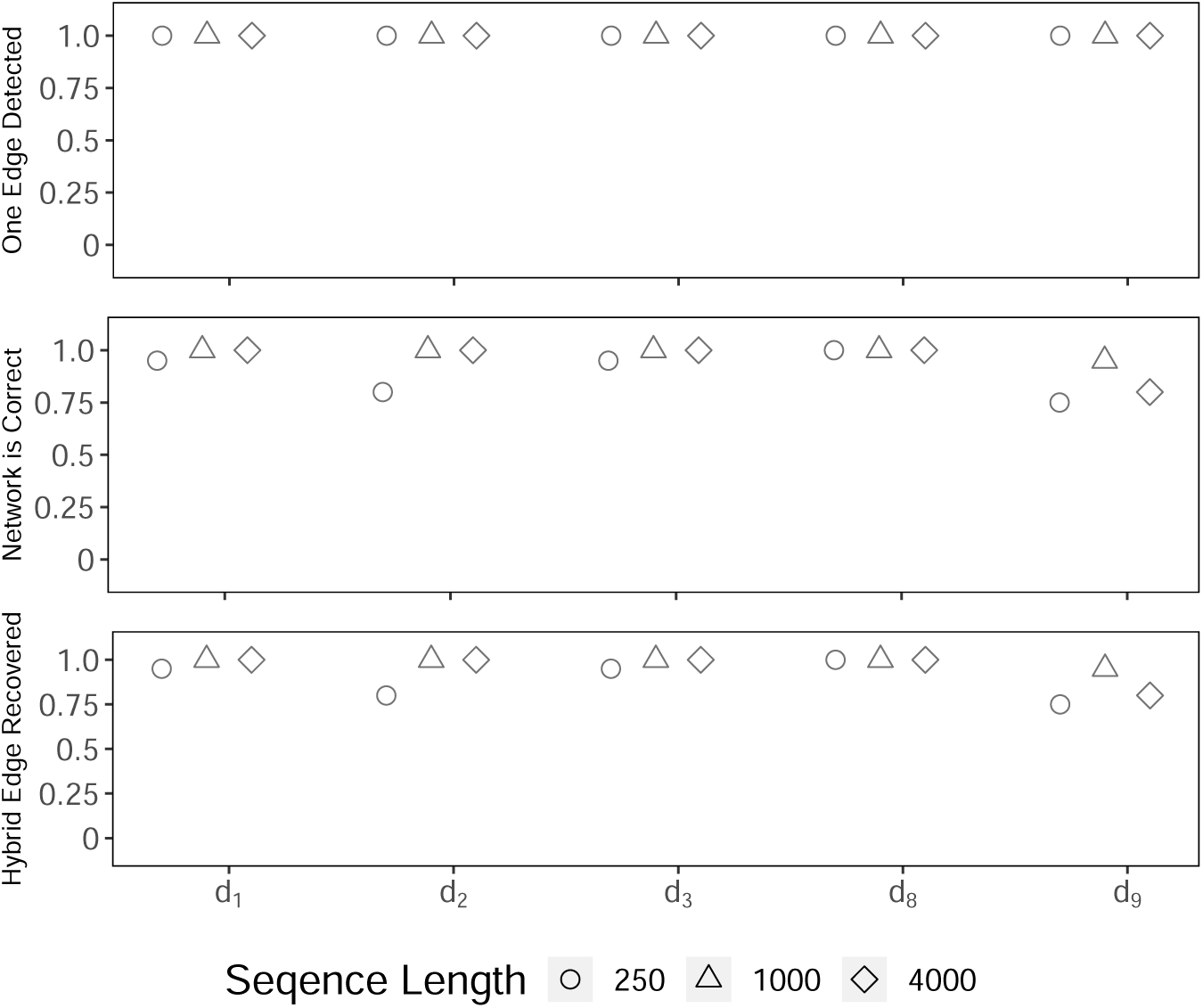
SNaQ results for root 3 with errors in gene trees. *One Edge Detected* is the proportion out of 20 replicates that 1 reticulate edge was correctly inferred by the pseudolikehood scores. *Network is Correct* is the proportion of 20 replicates where the estimated topology when allowing only one reticulation event is identical to the true topology. *Hybrid Edge Recovered* is the proportion of 20 replicates where the correct reticulation edge was inferred, regardless if other parts of the estimated network were incorrect, when allowing only one reticulation event. Diamonds correspond to Figure 4e.

## 6 Discussion

### 6.1 Mathematical insights

We have shown that the placement of the hybrid node in a 4-node hybridization cycle is generically identifiable in semi-directed level-1 phylogenetic networks. This result has important biological implications such as correct rooting of semi-directed network which provide more information about the actual speciation process from the origin of the clade. More work is needed to understand the identifiability of larger (and smaller) cycles. We decided to focus on the case of 4-node hybridization cycle because of empirical evidence suggesting flat pseudolikelihood on these hybridization events. However, smaller cycles are also biologically interesting as they describe hybridization events between more closely related populations. It has already been proven that 2-node cycles are not identifiable, but 3-node cycles are whenever we have sufficient sampling from the hybrid and sister clades [44, 46]. Thus, we can explore the identifiability of the placement of the hybrid node in the hybridization 3-node cycle as future work. In particular, we want to answer the following question:

**Question 1.** Let N_1_*, N*_2_*, N*_3_ *be the three rotations of the n-taxon semi-directed level-1 phylogenetic networks with one hybridization event producing a hybridization cycle of 3 nodes. Are N*_1_*, N*_2_*, N*_3_ *(generically) identifiable?*

Furthermore, all existing identifiability studies on semi-directed phylogenetic networks are restricted to a level-1 network [44, 46, 7, 17, 18, 2]. Questions on the identifiability of higher level networks remain open.

### 6.2 Biological insights from simulations and best practices

Pseudolikelihood methods can perform well for estimating a level-1 network from gene tree distributions in the absence of gene tree error, consistent with previous simulations and empirical analyses [55, 44]. PhyloNet’s I*N*fer*N*etwork_MPL was more efficient than PhyloNetwork’s SNaQ for diamonds *d*_2_, *d*_6_, and *d*_9_, capable of recovering the correct network with as few as 100 gene trees where SNaQ would require at least 500 (Figure 5). However, I*N*fer*N*etwork_MPL was not always perfect even with error-free gene trees such as *d*_8_ for *r*_3_. In addition, SNaQ was far more robust to the types of gene tree errors explored here than I*N*fer*N*etwork_MPL, capable of detecting one reticulation event and recovering the correct network in most conditions when provided a sufficient number of gene trees (Figure 7). I*N*fer*N*etwork_MPL could still detect the presence of a reticulation edge more often than not depending on the diamond analysed with *r*_3_, but the estimated network was frequently incorrect and if one were using unknown empirical data, there is a risk of estimating additional false positive reticulation nodes in the presence of gene tree rooting errors (Supplementary Data).

The diamonds and rootings where network estimation errors were most prevalent have implications for empirical analyses. SNaQ errors primarily occurred for *d*_2_, *d*_6_, and *d*_9_, all diamonds where the hybrid node descendant was a single taxon (Figure 4e). This implies that sampling two or more individuals for the lineage of interest could provide an increase in statistical power over a single taxon, at least for SNaQ and methods that use unrooted gene trees. Interestingly, network estimation errors were most prevalent when the cycle was restricted to an ingroup (*r*_3_). I*N*fer*N*etwork_MPL especially struggled with *d*_4_, *d*_7_, and *d*_9_ in the presence of errors, as it appears more data would not be helpful. Precisely why these diamonds performed poorly is unclear, but it could be an artifact of how errors were introduced to simulations. For example, *d*_4_ and *d*_7_ both have cherries for the descendants of the hybrid node. Since the outgroup errors were done randomly, this created more chances for a descendant of the hybrid node to be the outgroup. Outgroup lineages are typically sampled at an appropriate evoltionary distance to polarize phylogenetic relationships or site patterns in the context of *D*-statistics [16] [13] [38], but our results show that rooting errors and potentially other sources of topological errors (e.g. deep paralogy, assembly or genotyping, biases in starting material such as museum tissue vs fresh) could be misleading for estimating level-1 networks with rooted triples. An unrooted quartet-based method such as SNaQ should be more robust to gene tree error, but rooted triple-based methods should perform well and can be beneficial for a low number of gene trees (e.g. 100) if gene tree quality can be reliably evaluated.

## Acknowledgements

This project has received funding from the European Union’s Horizon 2020 research and innovation programme under the Marie Sklodowska-Curie grant agreement No. 101026923, awarded to GPT. This work was also funded by the National Science Foundation [DEB-2144367 to CSL].

## A Phylogenetic networks as a set of polynomial equations

### A.1 CF equations for *N_down_*: *CF* (*N_down_, z, γ*)

**Table 1:**
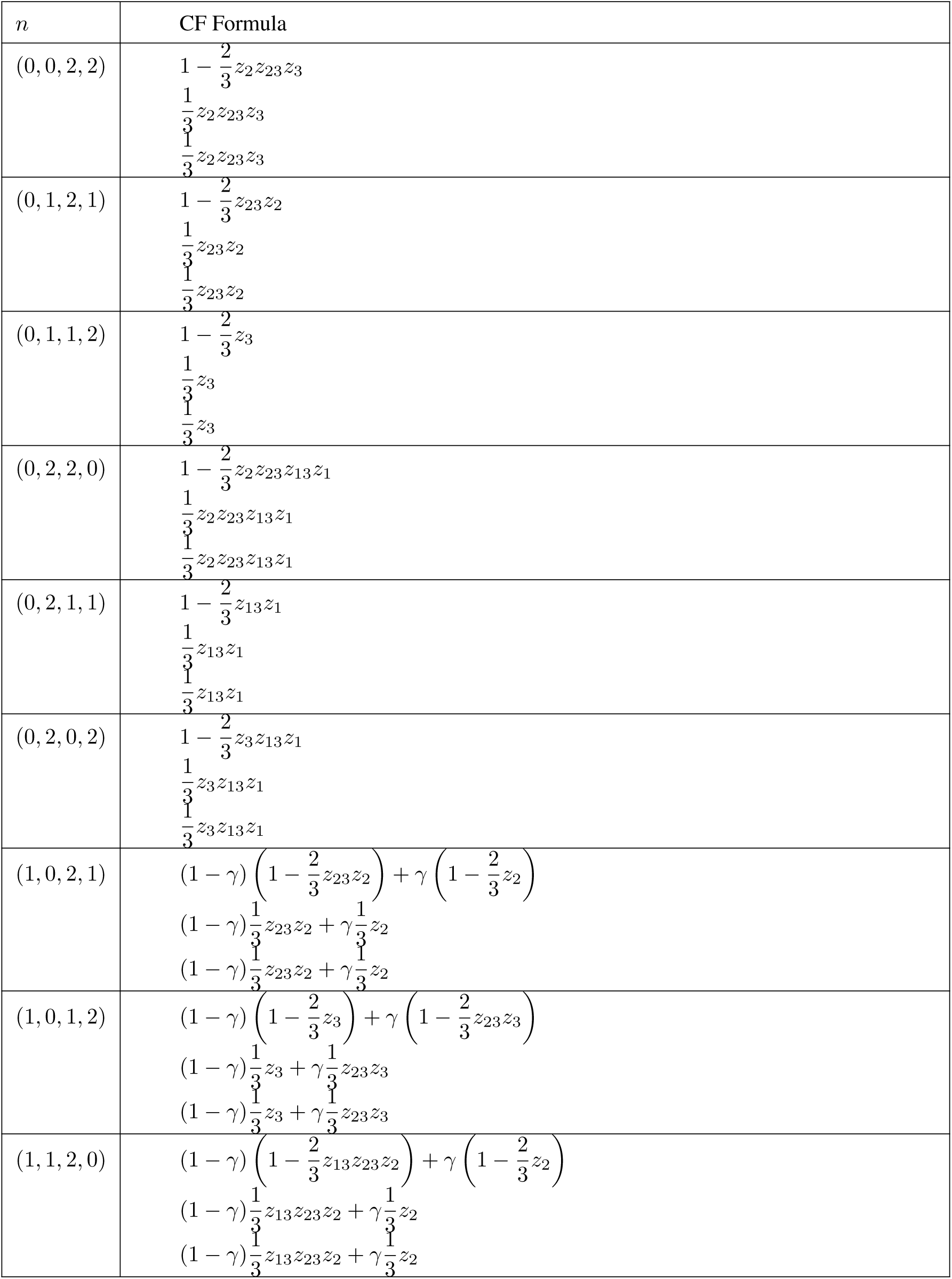
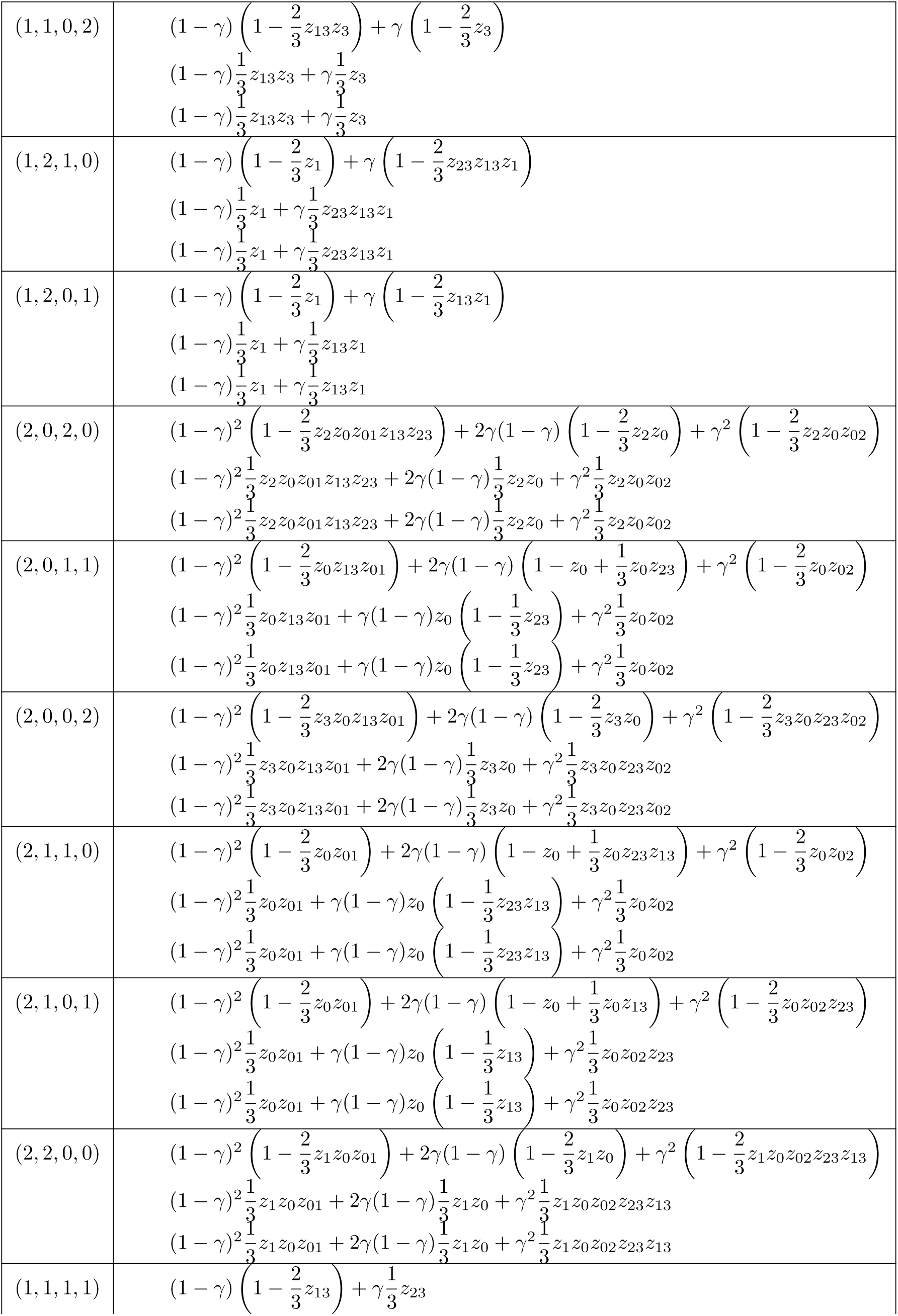
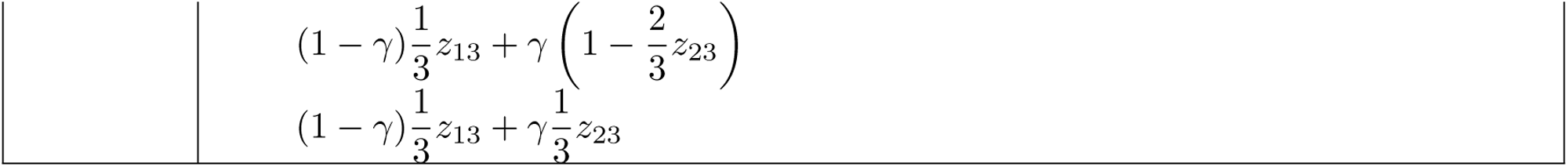
CF equations for *N_down_*: *CF* (*N_down_, z, γ*)

### A.**2** Definition of CF equations for network obtained by the rotation of hybrid in hybridization cycle

We start by noticing that there is a correspondence between a quartet in the network *N_down_* and a quartet in any other network. For example, in Figure 9, we want to match the equations corresponding to *n* = (0, 1, 2, 1) in *N_down_* and *N_right_*. We can see that the equations for the quartet *n* = (0, 1, 2, 1) in *N_right_* can be obtained from the equations in quartet *n* = (2, 0, 1, 1) in *N_down_*.

**Figure 9:**
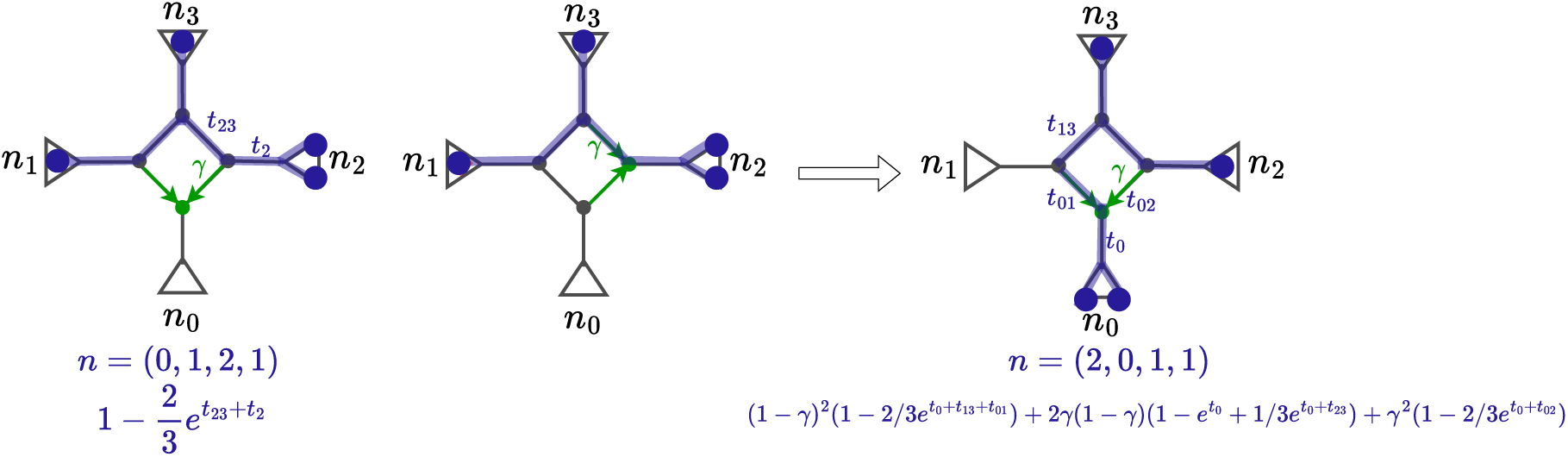
Correspondence between the quartet *n* = (0, 1, 2, 1) in *N_down_* (left), the same quartet in *N_right_* (center) and the quartet *n* = (2, 0, 1, 1) in *N_down_* (right).

Therefore, in order to get the equations for *N_right_*, we need to identify to which quartets they correspond in the *N_down_*.

Note that only the last quartet (1, 1, 1, 1) is not a mere rotation. We have to compute the CFs for the specific network (Figure 10).

**Figure 10:**
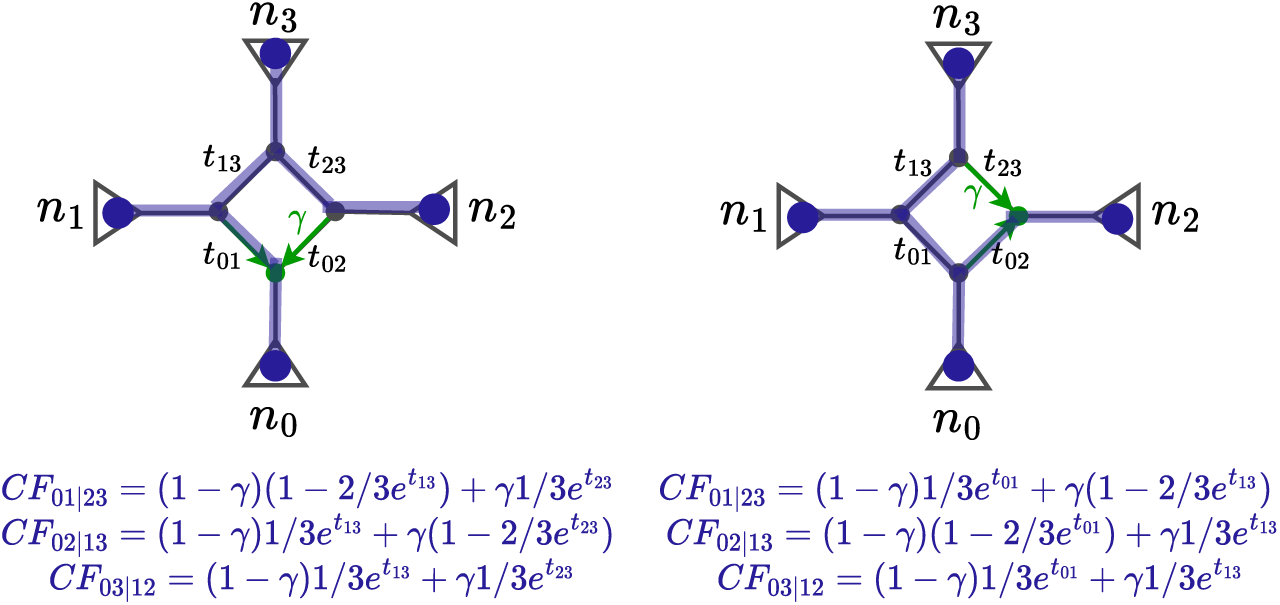
CF equations for quartet (1, 1, 1, 1) comparing the *N_down_* and *N_right_* networks.

### A.3 CF equations for *N_right_*: *CF* (*N_right_, z, γ*)

**Table 2:** CF equations for *N_right_*: *CF* (*N_right_, z, γ*)

| $n \in \mathcal{N}_{right}$ | $n \in \mathcal{N}_{down}$ | CF Formula |
| --- | --- | --- |
| (0, 0, 2, 2) | (2, 0, 2, 0) | $(1 - \gamma)^2 \left(1 - \frac{2}{3} z_2 z_0 z_{01} z_{13} z_{23}\right) + 2\gamma(1 - \gamma) \left(1 - \frac{2}{3} z_2 z_0\right) + \gamma^2 \left(1 - \frac{2}{3} z_2 z_0 z_{02}\right)$<br>$(1 - \gamma)^2 \frac{1}{3} z_2 z_0 z_{01} z_{13} z_{23} + 2\gamma(1 - \gamma) \frac{1}{3} z_2 z_0 + \gamma^2 \frac{1}{3} z_2 z_0 z_{02}$<br>$(1 - \gamma)^2 \frac{1}{3} z_2 z_0 z_{01} z_{13} z_{23} + 2\gamma(1 - \gamma) \frac{1}{3} z_2 z_0 + \gamma^2 \frac{1}{3} z_2 z_0 z_{02}$ |
| (0, 1, 2, 1) | (2, 0, 1, 1) | $(1 - \gamma)^2 \left(1 - \frac{2}{3} z_0 z_{13} z_{01}\right) + 2\gamma(1 - \gamma) \left(1 - z_0 + \frac{1}{3} z_0 z_{23}\right) + \gamma^2 \left(1 - \frac{2}{3} z_0 z_{02}\right)$<br>$(1 - \gamma)^2 \frac{1}{3} z_0 z_{13} z_{01} + \gamma(1 - \gamma) z_0 \left(1 - \frac{1}{3} z_{23}\right) + \gamma^2 \frac{1}{3} z_0 z_{02}$<br>$(1 - \gamma)^2 \frac{1}{3} z_0 z_{13} z_{01} + \gamma(1 - \gamma) z_0 \left(1 - \frac{1}{3} z_{23}\right) + \gamma^2 \frac{1}{3} z_0 z_{02}$ |
| (0, 1, 1, 2) | (1, 0, 2, 1) | $(1 - \gamma) \left(1 - \frac{2}{3} z_{23} z_2\right) + \gamma \left(1 - \frac{2}{3} z_2\right)$<br>$(1 - \gamma) \frac{1}{3} z_{23} z_2 + \gamma \frac{1}{3} z_2$<br>$(1 - \gamma) \frac{1}{3} z_{23} z_2 + \gamma \frac{1}{3} z_2$ |
| (0, 2, 2, 0) | (2, 0, 0, 2) | $(1 - \gamma)^2 \left(1 - \frac{2}{3} z_3 z_0 z_{13} z_{01}\right) + 2\gamma(1 - \gamma) \left(1 - \frac{2}{3} z_3 z_0\right) + \gamma^2 \left(1 - \frac{2}{3} z_3 z_0 z_{23} z_{02}\right)$<br>$(1 - \gamma)^2 \frac{1}{3} z_3 z_0 z_{13} z_{01} + 2\gamma(1 - \gamma) \frac{1}{3} z_3 z_0 + \gamma^2 \frac{1}{3} z_3 z_0 z_{23} z_{02}$<br>$(1 - \gamma)^2 \frac{1}{3} z_3 z_0 z_{13} z_{01} + 2\gamma(1 - \gamma) \frac{1}{3} z_3 z_0 + \gamma^2 \frac{1}{3} z_3 z_0 z_{23} z_{02}$ |
| (0, 2, 1, 1) | (1, 0, 1, 2) | $(1 - \gamma) \left(1 - \frac{2}{3} z_3\right) + \gamma \left(1 - \frac{2}{3} z_{23} z_3\right)$<br>$(1 - \gamma) \frac{1}{3} z_3 + \gamma \frac{1}{3} z_{23} z_3$<br>$(1 - \gamma) \frac{1}{3} z_3 + \gamma \frac{1}{3} z_{23} z_3$ |
| (0, 2, 0, 2) | (0, 0, 2, 2) | $1 - \frac{2}{3} z_2 z_{23} z_3$<br>$\frac{1}{3} z_2 z_{23} z_3$<br>$\frac{1}{3} z_2 z_{23} z_3$ |
| (1, 0, 2, 1) | (2, 1, 1, 0) | $(1 - \gamma)^2 \left(1 - \frac{2}{3} z_0 z_{01}\right) + 2\gamma(1 - \gamma) \left(1 - z_0 + \frac{1}{3} z_0 z_{23} z_{13}\right) + \gamma^2 \left(1 - \frac{2}{3} z_0 z_{02}\right)$<br>$(1 - \gamma)^2 \frac{1}{3} z_0 z_{01} + \gamma(1 - \gamma) z_0 \left(1 - \frac{1}{3} z_{23} z_{13}\right) + \gamma^2 \frac{1}{3} z_0 z_{02}$<br>$(1 - \gamma)^2 \frac{1}{3} z_0 z_{01} + \gamma(1 - \gamma) z_0 \left(1 - \frac{1}{3} z_{23} z_{13}\right) + \gamma^2 \frac{1}{3} z_0 z_{02}$ |
| (1, 0, 1, 2) | (1, 1, 2, 0) | $(1 - \gamma) \left(1 - \frac{2}{3} z_{13} z_{23} z_2\right) + \gamma \left(1 - \frac{2}{3} z_2\right)$<br>$(1 - \gamma) \frac{1}{3} z_{13} z_{23} z_2 + \gamma \frac{1}{3} z_2$<br>$(1 - \gamma) \frac{1}{3} z_{13} z_{23} z_2 + \gamma \frac{1}{3} z_2$ |
| (1, 1, 2, 0) | (2, 1, 0, 1) | $(1 - \gamma)^2 \left(1 - \frac{2}{3} z_0 z_{01}\right) + 2\gamma(1 - \gamma) \left(1 - z_0 + \frac{1}{3} z_0 z_{13}\right) + \gamma^2 \left(1 - \frac{2}{3} z_0 z_{02} z_{23}\right)$<br>$(1 - \gamma)^2 \frac{1}{3} z_0 z_{01} + \gamma(1 - \gamma) z_0 \left(1 - \frac{1}{3} z_{13}\right) + \gamma^2 \frac{1}{3} z_0 z_{02} z_{23}$<br>$(1 - \gamma)^2 \frac{1}{3} z_0 z_{01} + \gamma(1 - \gamma) z_0 \left(1 - \frac{1}{3} z_{13}\right) + \gamma^2 \frac{1}{3} z_0 z_{02} z_{23}$ |
| (1, 1, 0, 2) | (0, 1, 2, 1) | $1 - \frac{2}{3}z_{23}z_2$<br>$\frac{1}{3}z_{23}z_2$<br>$\frac{1}{3}z_{23}z_2$ |
| (1, 2, 1, 0) | (1, 1, 0, 2) | $(1 - \gamma) \left( 1 - \frac{2}{3}z_{13}z_3 \right) + \gamma \left( 1 - \frac{2}{3}z_3 \right)$<br>$(1 - \gamma) \frac{1}{3}z_{13}z_3 + \gamma \frac{1}{3}z_3$<br>$(1 - \gamma) \frac{1}{3}z_{13}z_3 + \gamma \frac{1}{3}z_3$ |
| (1, 2, 0, 1) | (0, 1, 1, 2) | $1 - \frac{2}{3}z_3$<br>$\frac{1}{3}z_3$<br>$\frac{1}{3}z_3$ |
| (2, 0, 2, 0) | (2, 2, 0, 0) | $(1 - \gamma)^2 \left( 1 - \frac{2}{3}z_1z_0z_{01} \right) + 2\gamma(1 - \gamma) \left( 1 - \frac{2}{3}z_1z_0 \right) + \gamma^2 \left( 1 - \frac{2}{3}z_1z_0z_{02}z_{23}z_{13} \right)$<br>$(1 - \gamma)^2 \frac{1}{3}z_1z_0z_{01} + 2\gamma(1 - \gamma) \frac{1}{3}z_1z_0 + \gamma^2 \frac{1}{3}z_1z_0z_{02}z_{23}z_{13}$<br>$(1 - \gamma)^2 \frac{1}{3}z_1z_0z_{01} + 2\gamma(1 - \gamma) \frac{1}{3}z_1z_0 + \gamma^2 \frac{1}{3}z_1z_0z_{02}z_{23}z_{13}$ |
| (2, 0, 1, 1) | (1, 2, 1, 0) | $(1 - \gamma) \left( 1 - \frac{2}{3}z_1 \right) + \gamma \left( 1 - \frac{2}{3}z_{23}z_{13}z_1 \right)$<br>$(1 - \gamma) \frac{1}{3}z_1 + \gamma \frac{1}{3}z_{23}z_{13}z_1$<br>$(1 - \gamma) \frac{1}{3}z_1 + \gamma \frac{1}{3}z_{23}z_{13}z_1$ |
| (2, 0, 0, 2) | (0, 2, 2, 0) | $1 - \frac{2}{3}z_2z_{23}z_{13}z_1$<br>$\frac{1}{3}z_2z_{23}z_{13}z_1$<br>$\frac{1}{3}z_2z_{23}z_{13}z_1$ |
| (2, 1, 1, 0) | (1, 2, 0, 1) | $(1 - \gamma) \left( 1 - \frac{2}{3}z_1 \right) + \gamma \left( 1 - \frac{2}{3}z_{13}z_1 \right)$<br>$(1 - \gamma) \frac{1}{3}z_1 + \gamma \frac{1}{3}z_{13}z_1$<br>$(1 - \gamma) \frac{1}{3}z_1 + \gamma \frac{1}{3}z_{13}z_1$ |
| (2, 1, 0, 1) | (0, 2, 1, 1) | $1 - \frac{2}{3}z_{13}z_1$<br>$\frac{1}{3}z_{13}z_1$<br>$\frac{1}{3}z_{13}z_1$ |
| (2, 2, 0, 0) | (0, 2, 0, 2) | $1 - \frac{2}{3}z_3z_{13}z_1$<br>$\frac{1}{3}z_3z_{13}z_1$<br>$\frac{1}{3}z_3z_{13}z_1$ |
| (1, 1, 1, 1) | | $(1 - \gamma) \frac{1}{3}z_{01} + \gamma \left( 1 - \frac{2}{3}z_{13} \right)$<br>$(1 - \gamma) \left( 1 - \frac{2}{3}z_{01} \right) + \gamma \frac{1}{3}z_{13}$<br>$(1 - \gamma) \frac{1}{3}z_{01} + \gamma \frac{1}{3}z_{13}$ |

### A.4 CF equations for *N_left_*: *CF* (*N_left_, z, γ*)

**Table 3:**
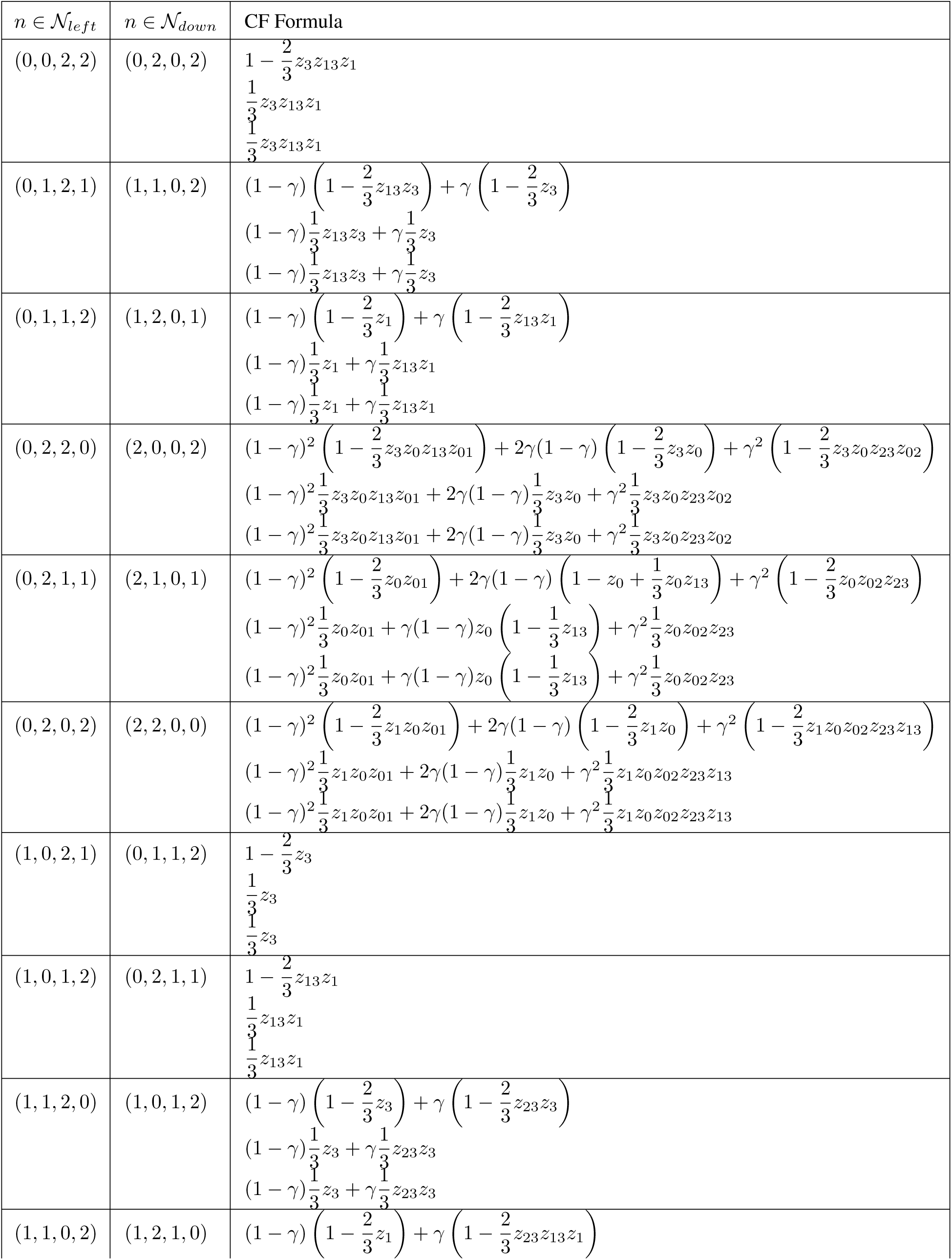
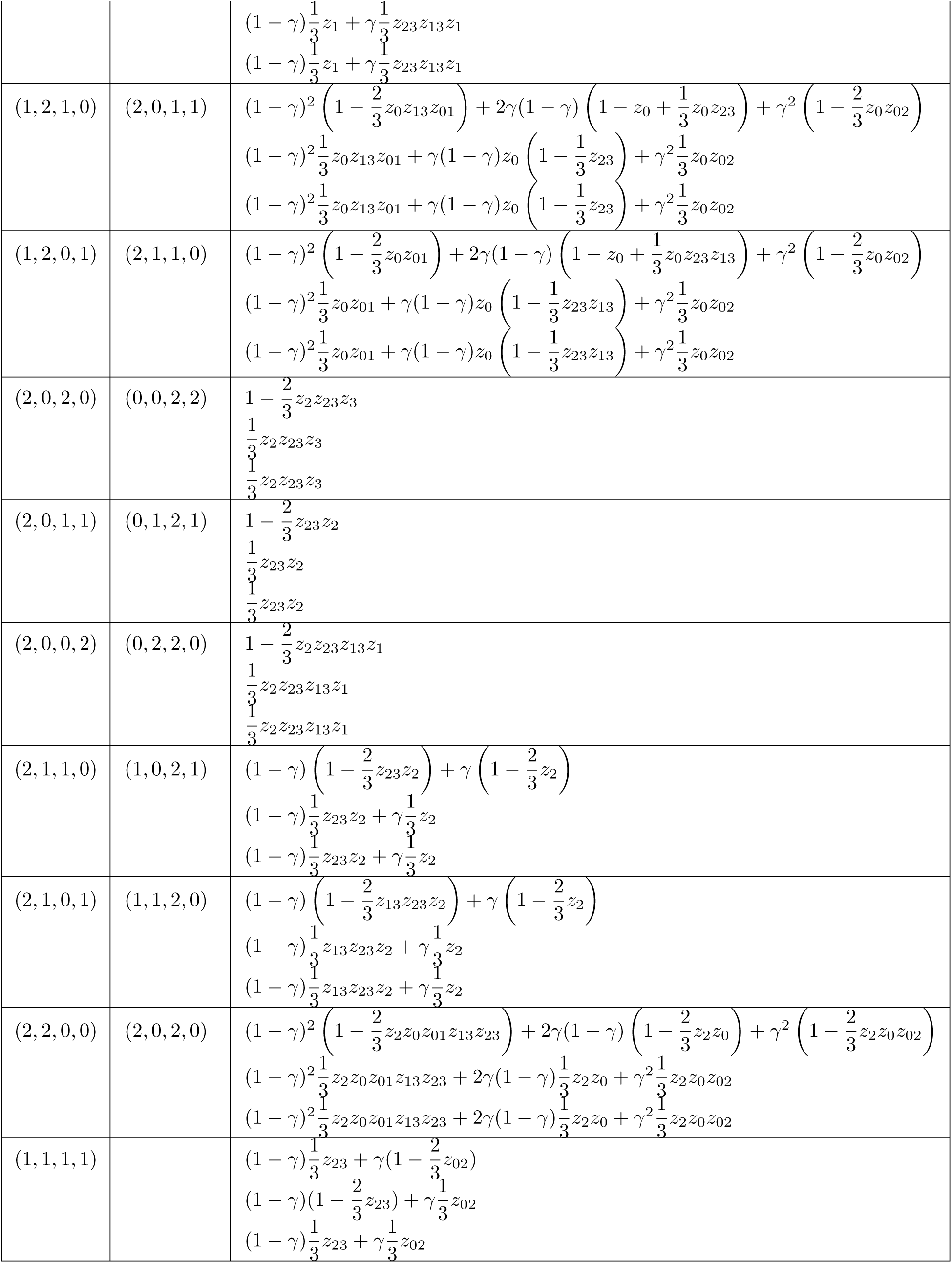
CF equations for *N_left_*: *CF* (*N_left_, z, γ*)

### A.5 CF equations for *N_up_*: *CF* (*N_up_, z, γ*)

**Table 4:** CF equations for *N_up_*: *CF* (*N_up_, z, γ*)

| $n \in \mathcal{N}_{up}$ | $n \in \mathcal{N}_{down}$ | CF Formula |
| --- | --- | --- |
| (0, 0, 2, 2) | (2, 2, 0, 0) | $(1 - \gamma)^2 \left(1 - \frac{2}{3} z_1 z_0 z_{01}\right) + 2\gamma(1 - \gamma) \left(1 - \frac{2}{3} z_1 z_0\right) + \gamma^2 \left(1 - \frac{2}{3} z_1 z_0 z_{02} z_{23} z_{13}\right)$ $(1 - \gamma)^2 \frac{1}{3} z_1 z_0 z_{01} + 2\gamma(1 - \gamma) \frac{1}{3} z_1 z_0 + \gamma^2 \frac{1}{3} z_1 z_0 z_{02} z_{23} z_{13}$ $(1 - \gamma)^2 \frac{1}{3} z_1 z_0 z_{01} + 2\gamma(1 - \gamma) \frac{1}{3} z_1 z_0 + \gamma^2 \frac{1}{3} z_1 z_0 z_{02} z_{23} z_{13}$ |
| (0, 1, 2, 1) | (1, 2, 1, 0) | $(1 - \gamma) \left(1 - \frac{2}{3} z_1\right) + \gamma \left(1 - \frac{2}{3} z_{23} z_{13} z_1\right)$ $(1 - \gamma) \frac{1}{3} z_1 + \gamma \frac{1}{3} z_{23} z_{13} z_1$ $(1 - \gamma) \frac{1}{3} z_1 + \gamma \frac{1}{3} z_{23} z_{13} z_1$ |
| (0, 1, 1, 2) | (2, 1, 1, 0) | $(1 - \gamma)^2 \left(1 - \frac{2}{3} z_0 z_{01}\right) + 2\gamma(1 - \gamma) \left(1 - z_0 + \frac{1}{3} z_0 z_{23} z_{13}\right) + \gamma^2 \left(1 - \frac{2}{3} z_0 z_{02}\right)$ $(1 - \gamma)^2 \frac{1}{3} z_0 z_{01} + \gamma(1 - \gamma) z_0 \left(1 - \frac{1}{3} z_{23} z_{13}\right) + \gamma^2 \frac{1}{3} z_0 z_{02}$ $(1 - \gamma)^2 \frac{1}{3} z_0 z_{01} + \gamma(1 - \gamma) z_0 \left(1 - \frac{1}{3} z_{23} z_{13}\right) + \gamma^2 \frac{1}{3} z_0 z_{02}$ |
| (0, 2, 2, 0) | (0, 2, 2, 0) | $1 - \frac{2}{3} z_2 z_{23} z_{13} z_1$ $\frac{1}{3} z_2 z_{23} z_{13} z_1$ $\frac{1}{3} z_2 z_{23} z_{13} z_1$ |
| (0, 2, 1, 1) | (1, 1, 2, 0) | $(1 - \gamma) \left(1 - \frac{2}{3} z_{13} z_{23} z_2\right) + \gamma \left(1 - \frac{2}{3} z_2\right)$ $(1 - \gamma) \frac{1}{3} z_{13} z_{23} z_2 + \gamma \frac{1}{3} z_2$ $(1 - \gamma) \frac{1}{3} z_{13} z_{23} z_2 + \gamma \frac{1}{3} z_2$ |
| (0, 2, 0, 2) | (2, 0, 2, 0) | $(1 - \gamma)^2 \left(1 - \frac{2}{3} z_2 z_0 z_{01} z_{13} z_{23}\right) + 2\gamma(1 - \gamma) \left(1 - \frac{2}{3} z_2 z_0\right) + \gamma^2 \left(1 - \frac{2}{3} z_2 z_0 z_{02}\right)$ $(1 - \gamma)^2 \frac{1}{3} z_2 z_0 z_{01} z_{13} z_{23} + 2\gamma(1 - \gamma) \frac{1}{3} z_2 z_0 + \gamma^2 \frac{1}{3} z_2 z_0 z_{02}$ $(1 - \gamma)^2 \frac{1}{3} z_2 z_0 z_{01} z_{13} z_{23} + 2\gamma(1 - \gamma) \frac{1}{3} z_2 z_0 + \gamma^2 \frac{1}{3} z_2 z_0 z_{02}$ |
| (1, 0, 2, 1) | (1, 2, 0, 1) | $(1 - \gamma) \left(1 - \frac{2}{3} z_1\right) + \gamma \left(1 - \frac{2}{3} z_{13} z_1\right)$ $(1 - \gamma) \frac{1}{3} z_1 + \gamma \frac{1}{3} z_{13} z_1$ $(1 - \gamma) \frac{1}{3} z_1 + \gamma \frac{1}{3} z_{13} z_1$ |
| (1, 0, 1, 2) | (2, 1, 0, 1) | $(1 - \gamma)^2 \left(1 - \frac{2}{3} z_0 z_{01}\right) + 2\gamma(1 - \gamma) \left(1 - z_0 + \frac{1}{3} z_0 z_{13}\right) + \gamma^2 \left(1 - \frac{2}{3} z_0 z_{02} z_{23}\right)$ $(1 - \gamma)^2 \frac{1}{3} z_0 z_{01} + \gamma(1 - \gamma) z_0 \left(1 - \frac{1}{3} z_{13}\right) + \gamma^2 \frac{1}{3} z_0 z_{02} z_{23}$ $(1 - \gamma)^2 \frac{1}{3} z_0 z_{01} + \gamma(1 - \gamma) z_0 \left(1 - \frac{1}{3} z_{13}\right) + \gamma^2 \frac{1}{3} z_0 z_{02} z_{23}$ |
| (1, 1, 2, 0) | (0, 2, 1, 1) | $1 - \frac{2}{3} z_{13} z_1$ $\frac{1}{3} z_{13} z_1$ $\frac{1}{3} z_{13} z_1$ |
| (1, 1, 0, 2) | (2, 0, 1, 1) | $(1 - \gamma)^2 \left(1 - \frac{2}{3} z_0 z_{13} z_{01}\right) + 2\gamma(1 - \gamma) \left(1 - z_0 + \frac{1}{3} z_0 z_{23}\right) + \gamma^2 \left(1 - \frac{2}{3} z_0 z_{02}\right)$<br>$(1 - \gamma)^2 \frac{1}{3} z_0 z_{13} z_{01} + \gamma(1 - \gamma) z_0 \left(1 - \frac{1}{3} z_{23}\right) + \gamma^2 \frac{1}{3} z_0 z_{02}$<br>$(1 - \gamma)^2 \frac{1}{3} z_0 z_{13} z_{01} + \gamma(1 - \gamma) z_0 \left(1 - \frac{1}{3} z_{23}\right) + \gamma^2 \frac{1}{3} z_0 z_{02}$ |
| (1, 2, 1, 0) | (0, 1, 2, 1) | $1 - \frac{2}{3} z_{23} z_2$<br>$\frac{1}{3} z_{23} z_2$<br>$\frac{1}{3} z_{23} z_2$ |
| (1, 2, 0, 1) | (1, 0, 2, 1) | $(1 - \gamma) \left(1 - \frac{2}{3} z_{23} z_2\right) + \gamma \left(1 - \frac{2}{3} z_2\right)$<br>$(1 - \gamma) \frac{1}{3} z_{23} z_2 + \gamma \frac{1}{3} z_2$<br>$(1 - \gamma) \frac{1}{3} z_{23} z_2 + \gamma \frac{1}{3} z_2$ |
| (2, 0, 2, 0) | (0, 2, 0, 2) | $1 - \frac{2}{3} z_3 z_{13} z_1$<br>$\frac{1}{3} z_3 z_{13} z_1$<br>$\frac{1}{3} z_3 z_{13} z_1$ |
| (2, 0, 1, 1) | (1, 1, 0, 2) | $(1 - \gamma) \left(1 - \frac{2}{3} z_{13} z_3\right) + \gamma \left(1 - \frac{2}{3} z_3\right)$<br>$(1 - \gamma) \frac{1}{3} z_{13} z_3 + \gamma \frac{1}{3} z_3$<br>$(1 - \gamma) \frac{1}{3} z_{13} z_3 + \gamma \frac{1}{3} z_3$ |
| (2, 0, 0, 2) | (2, 0, 0, 2) | $(1 - \gamma)^2 \left(1 - \frac{2}{3} z_3 z_0 z_{13} z_{01}\right) + 2\gamma(1 - \gamma) \left(1 - \frac{2}{3} z_3 z_0\right) + \gamma^2 \left(1 - \frac{2}{3} z_3 z_0 z_{23} z_{02}\right)$<br>$(1 - \gamma)^2 \frac{1}{3} z_3 z_0 z_{13} z_{01} + 2\gamma(1 - \gamma) \frac{1}{3} z_3 z_0 + \gamma^2 \frac{1}{3} z_3 z_0 z_{23} z_{02}$<br>$(1 - \gamma)^2 \frac{1}{3} z_3 z_0 z_{13} z_{01} + 2\gamma(1 - \gamma) \frac{1}{3} z_3 z_0 + \gamma^2 \frac{1}{3} z_3 z_0 z_{23} z_{02}$ |
| (2, 1, 1, 0) | (0, 1, 1, 2) | $1 - \frac{2}{3} z_3$<br>$\frac{1}{3} z_3$<br>$\frac{1}{3} z_3$ |
| (2, 1, 0, 1) | (1, 0, 1, 2) | $(1 - \gamma) \left(1 - \frac{2}{3} z_3\right) + \gamma \left(1 - \frac{2}{3} z_{23} z_3\right)$<br>$(1 - \gamma) \frac{1}{3} z_3 + \gamma \frac{1}{3} z_{23} z_3$<br>$(1 - \gamma) \frac{1}{3} z_3 + \gamma \frac{1}{3} z_{23} z_3$ |
| (2, 2, 0, 0) | (0, 0, 2, 2) | $1 - \frac{2}{3} z_2 z_{23} z_3$<br>$\frac{1}{3} z_2 z_{23} z_3$<br>$\frac{1}{3} z_2 z_{23} z_3$ |
| (1, 1, 1, 1) | | $(1 - \gamma) \left(1 - \frac{2}{3} z_{02}\right) + \gamma \frac{1}{3} z_{01}$<br>$(1 - \gamma) \frac{1}{3} z_{02} + \gamma \left(1 - \frac{2}{3} z_{01}\right)$<br>$(1 - \gamma) \frac{1}{3} z_{02} + \gamma \frac{1}{3} z_{01}$ |

## B Phylogenetic networks as a set of polynomial equations: ***n* = 6** taxa

### B.1 CF equations for 4-cycle network *N_down_* for *N* = 1212 (Diamond 6)

**Table 5:** CF equations for 4-cycle network *N_down_* for *N* = 1212 (Diamond 6)

| $n$ | CF Formula |
| --- | --- |
| $(0, 1, 1, 2)$ | $1 - \frac{2}{3}z_3$<br>$\frac{1}{3}z_3$<br>$\frac{1}{3}z_3$ |
| $(0, 2, 1, 1)$ | $1 - \frac{2}{3}z_{13}z_1$<br>$\frac{1}{3}z_{13}z_1$<br>$\frac{1}{3}z_{13}z_1$ |
| $(0, 2, 0, 2)$ | $1 - \frac{2}{3}z_3z_{13}z_1$<br>$\frac{1}{3}z_3z_{13}z_1$<br>$\frac{1}{3}z_3z_{13}z_1$ |
| $(1, 0, 1, 2)$ | $(1 - \gamma) \left( 1 - \frac{2}{3}z_3 \right) + \gamma \left( 1 - \frac{2}{3}z_{23}z_3 \right)$<br>$(1 - \gamma) \frac{1}{3}z_3 + \gamma \frac{1}{3}z_{23}z_3$<br>$(1 - \gamma) \frac{1}{3}z_3 + \gamma \frac{1}{3}z_{23}z_3$ |
| $(1, 1, 0, 2)$ | $(1 - \gamma) \left( 1 - \frac{2}{3}z_{13}z_3 \right) + \gamma \left( 1 - \frac{2}{3}z_3 \right)$<br>$(1 - \gamma) \frac{1}{3}z_{13}z_3 + \gamma \frac{1}{3}z_3$<br>$(1 - \gamma) \frac{1}{3}z_{13}z_3 + \gamma \frac{1}{3}z_3$ |
| $(1, 2, 1, 0)$ | $(1 - \gamma) \left( 1 - \frac{2}{3}z_1 \right) + \gamma \left( 1 - \frac{2}{3}z_{23}z_{13}z_1 \right)$<br>$(1 - \gamma) \frac{1}{3}z_1 + \gamma \frac{1}{3}z_{23}z_{13}z_1$<br>$(1 - \gamma) \frac{1}{3}z_1 + \gamma \frac{1}{3}z_{23}z_{13}z_1$ |
| $(1, 2, 0, 1)$ | $(1 - \gamma) \left( 1 - \frac{2}{3}z_1 \right) + \gamma \left( 1 - \frac{2}{3}z_{13}z_1 \right)$<br>$(1 - \gamma) \frac{1}{3}z_1 + \gamma \frac{1}{3}z_{13}z_1$<br>$(1 - \gamma) \frac{1}{3}z_1 + \gamma \frac{1}{3}z_{13}z_1$ |
| $(1, 1, 1, 1)$ | $(1 - \gamma) \left( 1 - \frac{2}{3}z_{13} \right) + \gamma \frac{1}{3}z_{23}$<br>$(1 - \gamma) \frac{1}{3}z_{13} + \gamma \left( 1 - \frac{2}{3}z_{23} \right)$<br>$(1 - \gamma) \frac{1}{3}z_{13} + \gamma \frac{1}{3}z_{23}$ |

### B.2 CF equations for 4-cycle network *N_right_* for *N* = 1122 (Diamond 9)

**Table 6:** CF equations for 4-cycle network *N_right_* for *N* = 1122 (Diamond 9)

| $n \in \mathcal{N}_{right}$ | $n \in \mathcal{N}_{down}$ | CF Formula |
| --- | --- | --- |
| $(0, 1, 1, 2)$ | $(1, 0, 2, 1)$ | $(1 - \gamma) \left( 1 - \frac{2}{3} z_{23} z_2 \right) + \gamma \left( 1 - \frac{2}{3} z_2 \right)$<br>$(1 - \gamma) \frac{1}{3} z_{23} z_2 + \gamma \frac{1}{3} z_2$<br>$(1 - \gamma) \frac{1}{3} z_{23} z_2 + \gamma \frac{1}{3} z_2$ |
| $(0, 2, 1, 1)$ | $(1, 0, 1, 2)$ | $(1 - \gamma) \left( 1 - \frac{2}{3} z_3 \right) + \gamma \left( 1 - \frac{2}{3} z_{23} z_3 \right)$<br>$(1 - \gamma) \frac{1}{3} z_3 + \gamma \frac{1}{3} z_{23} z_3$<br>$(1 - \gamma) \frac{1}{3} z_3 + \gamma \frac{1}{3} z_{23} z_3$ |
| $(0, 2, 0, 2)$ | $(0, 0, 2, 2)$ | $1 - \frac{2}{3} z_2 z_{23} z_3$<br>$\frac{1}{3} z_2 z_{23} z_3$<br>$\frac{1}{3} z_2 z_{23} z_3$ |
| $(1, 0, 1, 2)$ | $(1, 1, 2, 0)$ | $(1 - \gamma) \left( 1 - \frac{2}{3} z_{13} z_{23} z_2 \right) + \gamma \left( 1 - \frac{2}{3} z_2 \right)$<br>$(1 - \gamma) \frac{1}{3} z_{13} z_{23} z_2 + \gamma \frac{1}{3} z_2$<br>$(1 - \gamma) \frac{1}{3} z_{13} z_{23} z_2 + \gamma \frac{1}{3} z_2$ |
| $(1, 1, 0, 2)$ | $(0, 1, 2, 1)$ | $1 - \frac{2}{3} z_{23} z_2$<br>$\frac{1}{3} z_{23} z_2$<br>$\frac{1}{3} z_{23} z_2$ |
| $(1, 2, 1, 0)$ | $(1, 1, 0, 2)$ | $(1 - \gamma) \left( 1 - \frac{2}{3} z_{13} z_3 \right) + \gamma \left( 1 - \frac{2}{3} z_3 \right)$<br>$(1 - \gamma) \frac{1}{3} z_{13} z_3 + \gamma \frac{1}{3} z_3$<br>$(1 - \gamma) \frac{1}{3} z_{13} z_3 + \gamma \frac{1}{3} z_3$ |
| $(1, 2, 0, 1)$ | $(0, 1, 1, 2)$ | $1 - \frac{2}{3} z_3$<br>$\frac{1}{3} z_3$<br>$\frac{1}{3} z_3$ |
| $(1, 1, 1, 1)$ | | $(1 - \gamma) \frac{1}{3} z_{01} + \gamma \left( 1 - \frac{2}{3} z_{13} \right)$<br>$(1 - \gamma) \left( 1 - \frac{2}{3} z_{01} \right) + \gamma \frac{1}{3} z_{13}$<br>$(1 - \gamma) \frac{1}{3} z_{01} + \gamma \frac{1}{3} z_{13}$ |

### B.3 CF equations for 4-cycle network *N_left_* for *N* = 2211 (Diamond 7)

**Table 7:**
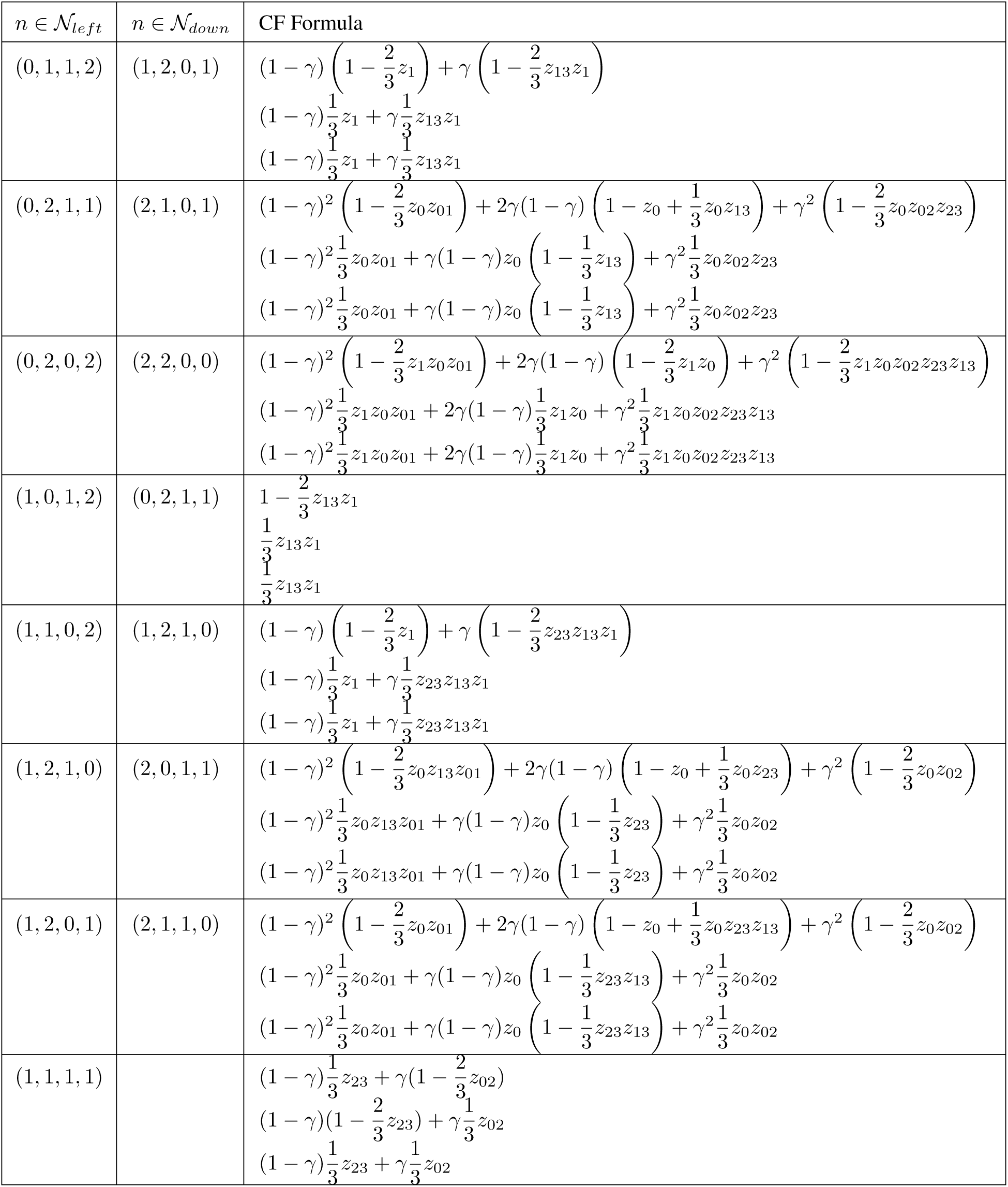
CF equations for 4-cycle network *N_left_* for *N* = 2211 (Diamond 7)

### B.4 CF equations for 4-cycle network *N_up_* for *N* = 2121 (Diamond 8)

**Table 8:** CF equations for 4-cycle network *N_up_* for *N* = 2121 (Diamond 8)

| $n \in \mathcal{N}_{up}$ | $n \in \mathcal{N}_{down}$ | CF Formula |
| --- | --- | --- |
| (0, 1, 1, 2) | (2, 1, 1, 0) | $(1 - \gamma)^2 \left(1 - \frac{2}{3} z_0 z_{01}\right) + 2\gamma(1 - \gamma) \left(1 - z_0 + \frac{1}{3} z_0 z_{23} z_{13}\right) + \gamma^2 \left(1 - \frac{2}{3} z_0 z_{02}\right)$ $(1 - \gamma)^2 \frac{1}{3} z_0 z_{01} + \gamma(1 - \gamma) z_0 \left(1 - \frac{1}{3} z_{23} z_{13}\right) + \gamma^2 \frac{1}{3} z_0 z_{02}$ $(1 - \gamma)^2 \frac{1}{3} z_0 z_{01} + \gamma(1 - \gamma) z_0 \left(1 - \frac{1}{3} z_{23} z_{13}\right) + \gamma^2 \frac{1}{3} z_0 z_{02}$ |
| (0, 2, 1, 1) | (1, 1, 2, 0) | $(1 - \gamma) \left(1 - \frac{2}{3} z_{13} z_{23} z_2\right) + \gamma \left(1 - \frac{2}{3} z_2\right)$ $(1 - \gamma) \frac{1}{3} z_{13} z_{23} z_2 + \gamma \frac{1}{3} z_2$ $(1 - \gamma) \frac{1}{3} z_{13} z_{23} z_2 + \gamma \frac{1}{3} z_2$ |
| (0, 2, 0, 2) | (2, 0, 2, 0) | $(1 - \gamma)^2 \left(1 - \frac{2}{3} z_2 z_0 z_{01} z_{13} z_{23}\right) + 2\gamma(1 - \gamma) \left(1 - \frac{2}{3} z_2 z_0\right) + \gamma^2 \left(1 - \frac{2}{3} z_2 z_0 z_{02}\right)$ $(1 - \gamma)^2 \frac{1}{3} z_2 z_0 z_{01} z_{13} z_{23} + 2\gamma(1 - \gamma) \frac{1}{3} z_2 z_0 + \gamma^2 \frac{1}{3} z_2 z_0 z_{02}$ $(1 - \gamma)^2 \frac{1}{3} z_2 z_0 z_{01} z_{13} z_{23} + 2\gamma(1 - \gamma) \frac{1}{3} z_2 z_0 + \gamma^2 \frac{1}{3} z_2 z_0 z_{02}$ |
| (1, 0, 1, 2) | (2, 1, 0, 1) | $(1 - \gamma)^2 \left(1 - \frac{2}{3} z_0 z_{01}\right) + 2\gamma(1 - \gamma) \left(1 - z_0 + \frac{1}{3} z_0 z_{13}\right) + \gamma^2 \left(1 - \frac{2}{3} z_0 z_{02} z_{23}\right)$ $(1 - \gamma)^2 \frac{1}{3} z_0 z_{01} + \gamma(1 - \gamma) z_0 \left(1 - \frac{1}{3} z_{13}\right) + \gamma^2 \frac{1}{3} z_0 z_{02} z_{23}$ $(1 - \gamma)^2 \frac{1}{3} z_0 z_{01} + \gamma(1 - \gamma) z_0 \left(1 - \frac{1}{3} z_{13}\right) + \gamma^2 \frac{1}{3} z_0 z_{02} z_{23}$ |
| (1, 1, 0, 2) | (2, 0, 1, 1) | $(1 - \gamma)^2 \left(1 - \frac{2}{3} z_0 z_{13} z_{01}\right) + 2\gamma(1 - \gamma) \left(1 - z_0 + \frac{1}{3} z_0 z_{23}\right) + \gamma^2 \left(1 - \frac{2}{3} z_0 z_{02}\right)$ $(1 - \gamma)^2 \frac{1}{3} z_0 z_{13} z_{01} + \gamma(1 - \gamma) z_0 \left(1 - \frac{1}{3} z_{23}\right) + \gamma^2 \frac{1}{3} z_0 z_{02}$ $(1 - \gamma)^2 \frac{1}{3} z_0 z_{13} z_{01} + \gamma(1 - \gamma) z_0 \left(1 - \frac{1}{3} z_{23}\right) + \gamma^2 \frac{1}{3} z_0 z_{02}$ |
| (1, 2, 1, 0) | (0, 1, 2, 1) | $1 - \frac{2}{3} z_{23} z_2$ $\frac{1}{3} z_{23} z_2$ $\frac{1}{3} z_{23} z_2$ |
| (1, 2, 0, 1) | (1, 0, 2, 1) | $(1 - \gamma) \left(1 - \frac{2}{3} z_{23} z_2\right) + \gamma \left(1 - \frac{2}{3} z_2\right)$ $(1 - \gamma) \frac{1}{3} z_{23} z_2 + \gamma \frac{1}{3} z_2$ $(1 - \gamma) \frac{1}{3} z_{23} z_2 + \gamma \frac{1}{3} z_2$ |
| (1, 1, 1, 1) | | $(1 - \gamma) \left(1 - \frac{2}{3} z_{02}\right) + \gamma \frac{1}{3} z_{01}$ $(1 - \gamma) \frac{1}{3} z_{02} + \gamma \left(1 - \frac{2}{3} z_{01}\right)$ $(1 - \gamma) \frac{1}{3} z_{02} + \gamma \frac{1}{3} z_{01}$ |

## C Phylogenetic networks as a set of polynomial equations: ***n* = 7** taxa

### C.1 CF equations for *N_down_* for *N* = 1222 (Diamond 2)

**Table 9:** CF equations for *N_down_* for *N* = 1222 (Diamond 2)

| $n$ | CF Formula |
| --- | --- |
| $(0, 0, 2, 2)$ | $1 - \frac{2}{3} z_2 z_{23} z_3$ $\frac{1}{3} z_2 z_{23} z_3$ $\frac{1}{3} z_2 z_{23} z_3$ |
| $(0, 1, 2, 1)$ | $1 - \frac{2}{3} z_{23} z_2$ $\frac{1}{3} z_{23} z_2$ $\frac{1}{3} z_{23} z_2$ |
| $(0, 1, 1, 2)$ | $1 - \frac{2}{3} z_3$ $\frac{1}{3} z_3$ $\frac{1}{3} z_3$ |
| $(0, 2, 2, 0)$ | $1 - \frac{2}{3} z_2 z_{23} z_{13} z_1$ $\frac{1}{3} z_2 z_{23} z_{13} z_1$ $\frac{1}{3} z_2 z_{23} z_{13} z_1$ |
| $(0, 2, 1, 1)$ | $1 - \frac{2}{3} z_{13} z_1$ $\frac{1}{3} z_{13} z_1$ $\frac{1}{3} z_{13} z_1$ |
| $(0, 2, 0, 2)$ | $1 - \frac{2}{3} z_3 z_{13} z_1$ $\frac{1}{3} z_3 z_{13} z_1$ $\frac{1}{3} z_3 z_{13} z_1$ |
| $(1, 0, 2, 1)$ | $(1 - \gamma) \left( 1 - \frac{2}{3} z_{23} z_2 \right) + \gamma \left( 1 - \frac{2}{3} z_2 \right)$ $(1 - \gamma) \frac{1}{3} z_{23} z_2 + \gamma \frac{1}{3} z_2$ $(1 - \gamma) \frac{1}{3} z_{23} z_2 + \gamma \frac{1}{3} z_2$ |
| $(1, 0, 1, 2)$ | $(1 - \gamma) \left( 1 - \frac{2}{3} z_3 \right) + \gamma \left( 1 - \frac{2}{3} z_{23} z_3 \right)$ $(1 - \gamma) \frac{1}{3} z_3 + \gamma \frac{1}{3} z_{23} z_3$ $(1 - \gamma) \frac{1}{3} z_3 + \gamma \frac{1}{3} z_{23} z_3$ |
| $(1, 1, 2, 0)$ | $(1 - \gamma) \left( 1 - \frac{2}{3} z_{13} z_{23} z_2 \right) + \gamma \left( 1 - \frac{2}{3} z_2 \right)$ $(1 - \gamma) \frac{1}{3} z_{13} z_{23} z_2 + \gamma \frac{1}{3} z_2$ $(1 - \gamma) \frac{1}{3} z_{13} z_{23} z_2 + \gamma \frac{1}{3} z_2$ |
| $(1, 1, 0, 2)$ | $(1 - \gamma) \left( 1 - \frac{2}{3} z_{13} z_3 \right) + \gamma \left( 1 - \frac{2}{3} z_3 \right)$ $(1 - \gamma) \frac{1}{3} z_{13} z_3 + \gamma \frac{1}{3} z_3$ $(1 - \gamma) \frac{1}{3} z_{13} z_3 + \gamma \frac{1}{3} z_3$ |
| $(1, 2, 1, 0)$ | $(1 - \gamma) \left( 1 - \frac{2}{3} z_1 \right) + \gamma \left( 1 - \frac{2}{3} z_{23} z_{13} z_1 \right)$ $(1 - \gamma) \frac{1}{3} z_1 + \gamma \frac{1}{3} z_{23} z_{13} z_1$ $(1 - \gamma) \frac{1}{3} z_1 + \gamma \frac{1}{3} z_{23} z_{13} z_1$ |
| $(1, 2, 0, 1)$ | $(1 - \gamma) \left( 1 - \frac{2}{3} z_1 \right) + \gamma \left( 1 - \frac{2}{3} z_{13} z_1 \right)$ $(1 - \gamma) \frac{1}{3} z_1 + \gamma \frac{1}{3} z_{13} z_1$ $(1 - \gamma) \frac{1}{3} z_1 + \gamma \frac{1}{3} z_{13} z_1$ |
| $(1, 1, 1, 1)$ | $(1 - \gamma) \left( 1 - \frac{2}{3} z_{13} \right) + \gamma \frac{1}{3} z_{23}$ $(1 - \gamma) \frac{1}{3} z_{13} + \gamma \left( 1 - \frac{2}{3} z_{23} \right)$ $(1 - \gamma) \frac{1}{3} z_{13} + \gamma \frac{1}{3} z_{23}$ |

### C.2 CF equations for *N_right_* for *N* = 2122 (Diamond 5)

**Table 10:**
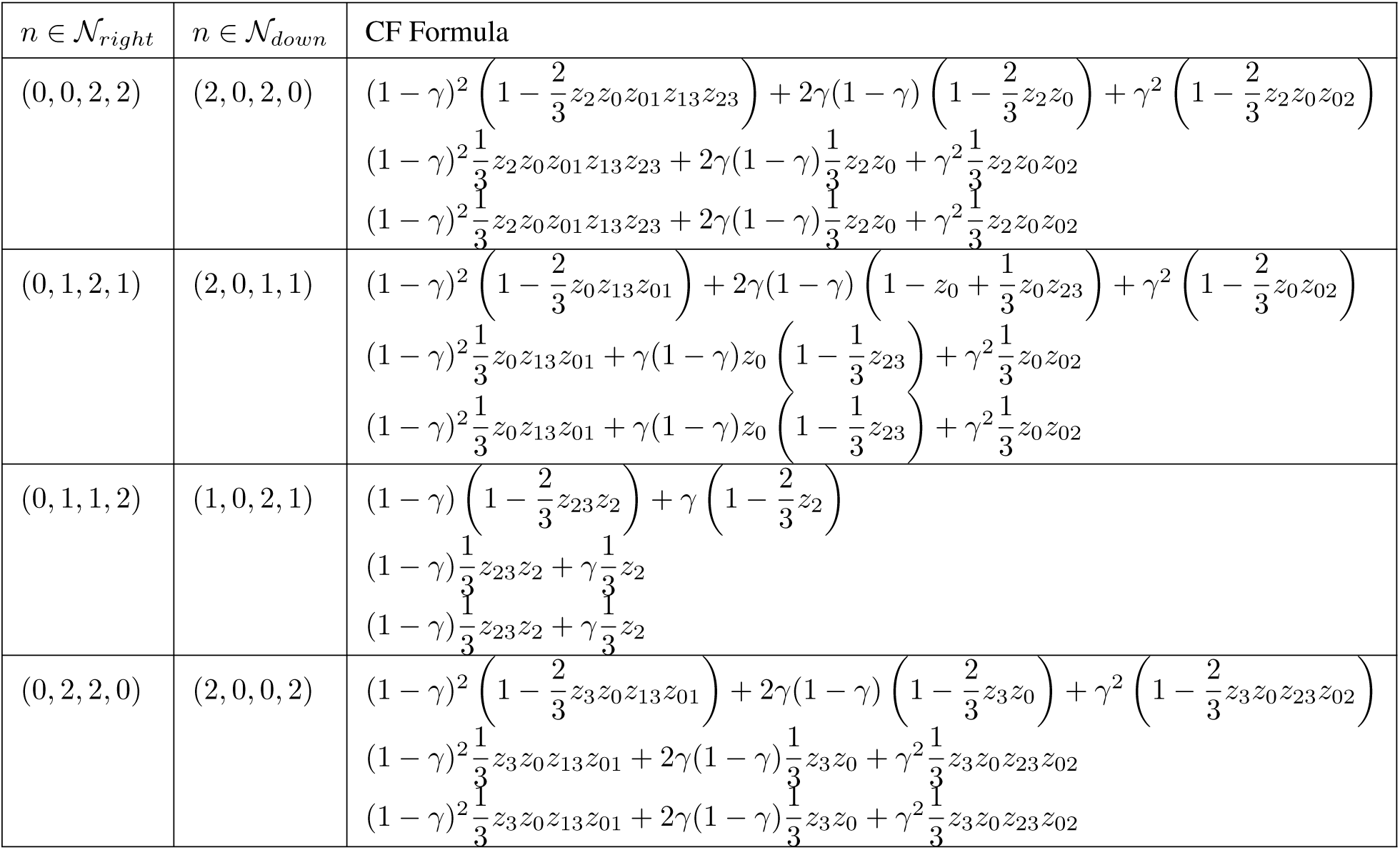
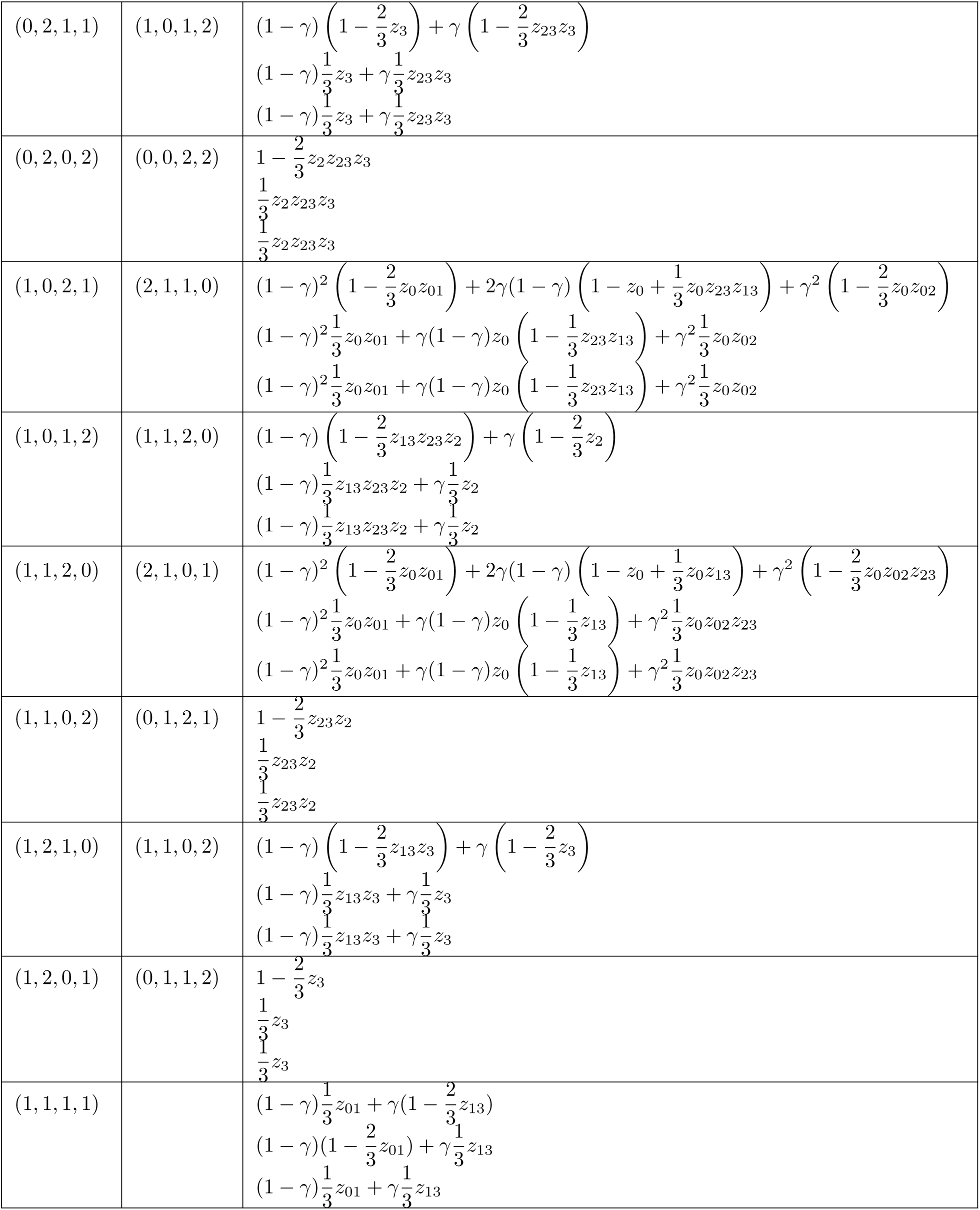
CF equations for *N_right_* for *N* = 2122 (Diamond 5)

| $n \in \mathcal{N}_{right}$ | $n \in \mathcal{N}_{down}$ | CF Formula |
| --- | --- | --- |
| $(0, 0, 2, 2)$ | $(2, 0, 2, 0)$ | $(1 - \gamma)^2 \left( 1 - \frac{2}{3} z_2 z_0 z_{01} z_{13} z_{23} \right) + 2\gamma(1 - \gamma) \left( 1 - \frac{2}{3} z_2 z_0 \right) + \gamma^2 \left( 1 - \frac{2}{3} z_2 z_0 z_{02} \right)$ $(1 - \gamma)^2 \frac{1}{3} z_2 z_0 z_{01} z_{13} z_{23} + 2\gamma(1 - \gamma) \frac{1}{3} z_2 z_0 + \gamma^2 \frac{1}{3} z_2 z_0 z_{02}$ $(1 - \gamma)^2 \frac{1}{3} z_2 z_0 z_{01} z_{13} z_{23} + 2\gamma(1 - \gamma) \frac{1}{3} z_2 z_0 + \gamma^2 \frac{1}{3} z_2 z_0 z_{02}$ |
| $(0, 1, 2, 1)$ | $(2, 0, 1, 1)$ | $(1 - \gamma)^2 \left( 1 - \frac{2}{3} z_0 z_{13} z_{01} \right) + 2\gamma(1 - \gamma) \left( 1 - z_0 + \frac{1}{3} z_0 z_{23} \right) + \gamma^2 \left( 1 - \frac{2}{3} z_0 z_{02} \right)$ $(1 - \gamma)^2 \frac{1}{3} z_0 z_{13} z_{01} + \gamma(1 - \gamma) z_0 \left( 1 - \frac{1}{3} z_{23} \right) + \gamma^2 \frac{1}{3} z_0 z_{02}$ $(1 - \gamma)^2 \frac{1}{3} z_0 z_{13} z_{01} + \gamma(1 - \gamma) z_0 \left( 1 - \frac{1}{3} z_{23} \right) + \gamma^2 \frac{1}{3} z_0 z_{02}$ |
| $(0, 1, 1, 2)$ | $(1, 0, 2, 1)$ | $(1 - \gamma) \left( 1 - \frac{2}{3} z_{23} z_2 \right) + \gamma \left( 1 - \frac{2}{3} z_2 \right)$ $(1 - \gamma) \frac{1}{3} z_{23} z_2 + \gamma \frac{1}{3} z_2$ $(1 - \gamma) \frac{1}{3} z_{23} z_2 + \gamma \frac{1}{3} z_2$ |
| $(0, 2, 2, 0)$ | $(2, 0, 0, 2)$ | $(1 - \gamma)^2 \left( 1 - \frac{2}{3} z_3 z_0 z_{13} z_{01} \right) + 2\gamma(1 - \gamma) \left( 1 - \frac{2}{3} z_3 z_0 \right) + \gamma^2 \left( 1 - \frac{2}{3} z_3 z_0 z_{23} z_{02} \right)$ $(1 - \gamma)^2 \frac{1}{3} z_3 z_0 z_{13} z_{01} + 2\gamma(1 - \gamma) \frac{1}{3} z_3 z_0 + \gamma^2 \frac{1}{3} z_3 z_0 z_{23} z_{02}$ $(1 - \gamma)^2 \frac{1}{3} z_3 z_0 z_{13} z_{01} + 2\gamma(1 - \gamma) \frac{1}{3} z_3 z_0 + \gamma^2 \frac{1}{3} z_3 z_0 z_{23} z_{02}$ |
| (0, 2, 1, 1) | (1, 0, 1, 2) | $(1 - \gamma) \left(1 - \frac{2}{3}z_3\right) + \gamma \left(1 - \frac{2}{3}z_{23}z_3\right)$<br>$(1 - \gamma)\frac{1}{3}z_3 + \gamma\frac{1}{3}z_{23}z_3$<br>$(1 - \gamma)\frac{1}{3}z_3 + \gamma\frac{1}{3}z_{23}z_3$ |
| (0, 2, 0, 2) | (0, 0, 2, 2) | $1 - \frac{2}{3}z_2z_{23}z_3$<br>$\frac{1}{3}z_2z_{23}z_3$<br>$\frac{1}{3}z_2z_{23}z_3$ |
| (1, 0, 2, 1) | (2, 1, 1, 0) | $(1 - \gamma)^2 \left(1 - \frac{2}{3}z_0z_{01}\right) + 2\gamma(1 - \gamma) \left(1 - z_0 + \frac{1}{3}z_0z_{23}z_{13}\right) + \gamma^2 \left(1 - \frac{2}{3}z_0z_{02}\right)$<br>$(1 - \gamma)^2 \frac{1}{3}z_0z_{01} + \gamma(1 - \gamma)z_0 \left(1 - \frac{1}{3}z_{23}z_{13}\right) + \gamma^2 \frac{1}{3}z_0z_{02}$<br>$(1 - \gamma)^2 \frac{1}{3}z_0z_{01} + \gamma(1 - \gamma)z_0 \left(1 - \frac{1}{3}z_{23}z_{13}\right) + \gamma^2 \frac{1}{3}z_0z_{02}$ |
| (1, 0, 1, 2) | (1, 1, 2, 0) | $(1 - \gamma) \left(1 - \frac{2}{3}z_{13}z_{23}z_2\right) + \gamma \left(1 - \frac{2}{3}z_2\right)$<br>$(1 - \gamma)\frac{1}{3}z_{13}z_{23}z_2 + \gamma\frac{1}{3}z_2$<br>$(1 - \gamma)\frac{1}{3}z_{13}z_{23}z_2 + \gamma\frac{1}{3}z_2$ |
| (1, 1, 2, 0) | (2, 1, 0, 1) | $(1 - \gamma)^2 \left(1 - \frac{2}{3}z_0z_{01}\right) + 2\gamma(1 - \gamma) \left(1 - z_0 + \frac{1}{3}z_0z_{13}\right) + \gamma^2 \left(1 - \frac{2}{3}z_0z_{02}z_{23}\right)$<br>$(1 - \gamma)^2 \frac{1}{3}z_0z_{01} + \gamma(1 - \gamma)z_0 \left(1 - \frac{1}{3}z_{13}\right) + \gamma^2 \frac{1}{3}z_0z_{02}z_{23}$<br>$(1 - \gamma)^2 \frac{1}{3}z_0z_{01} + \gamma(1 - \gamma)z_0 \left(1 - \frac{1}{3}z_{13}\right) + \gamma^2 \frac{1}{3}z_0z_{02}z_{23}$ |
| (1, 1, 0, 2) | (0, 1, 2, 1) | $1 - \frac{2}{3}z_{23}z_2$<br>$\frac{1}{3}z_{23}z_2$<br>$\frac{1}{3}z_{23}z_2$ |
| (1, 2, 1, 0) | (1, 1, 0, 2) | $(1 - \gamma) \left(1 - \frac{2}{3}z_{13}z_3\right) + \gamma \left(1 - \frac{2}{3}z_3\right)$<br>$(1 - \gamma)\frac{1}{3}z_{13}z_3 + \gamma\frac{1}{3}z_3$<br>$(1 - \gamma)\frac{1}{3}z_{13}z_3 + \gamma\frac{1}{3}z_3$ |
| (1, 2, 0, 1) | (0, 1, 1, 2) | $1 - \frac{2}{3}z_3$<br>$\frac{1}{3}z_3$<br>$\frac{1}{3}z_3$ |
| (1, 1, 1, 1) | | $(1 - \gamma)\frac{1}{3}z_{01} + \gamma\left(1 - \frac{2}{3}z_{13}\right)$<br>$(1 - \gamma)\left(1 - \frac{2}{3}z_{01}\right) + \gamma\frac{1}{3}z_{13}$<br>$(1 - \gamma)\frac{1}{3}z_{01} + \gamma\frac{1}{3}z_{13}$ |

### C.3 CF equations for *N_left_* for *N* = 2212 (Diamond 3)

**Table 11:**
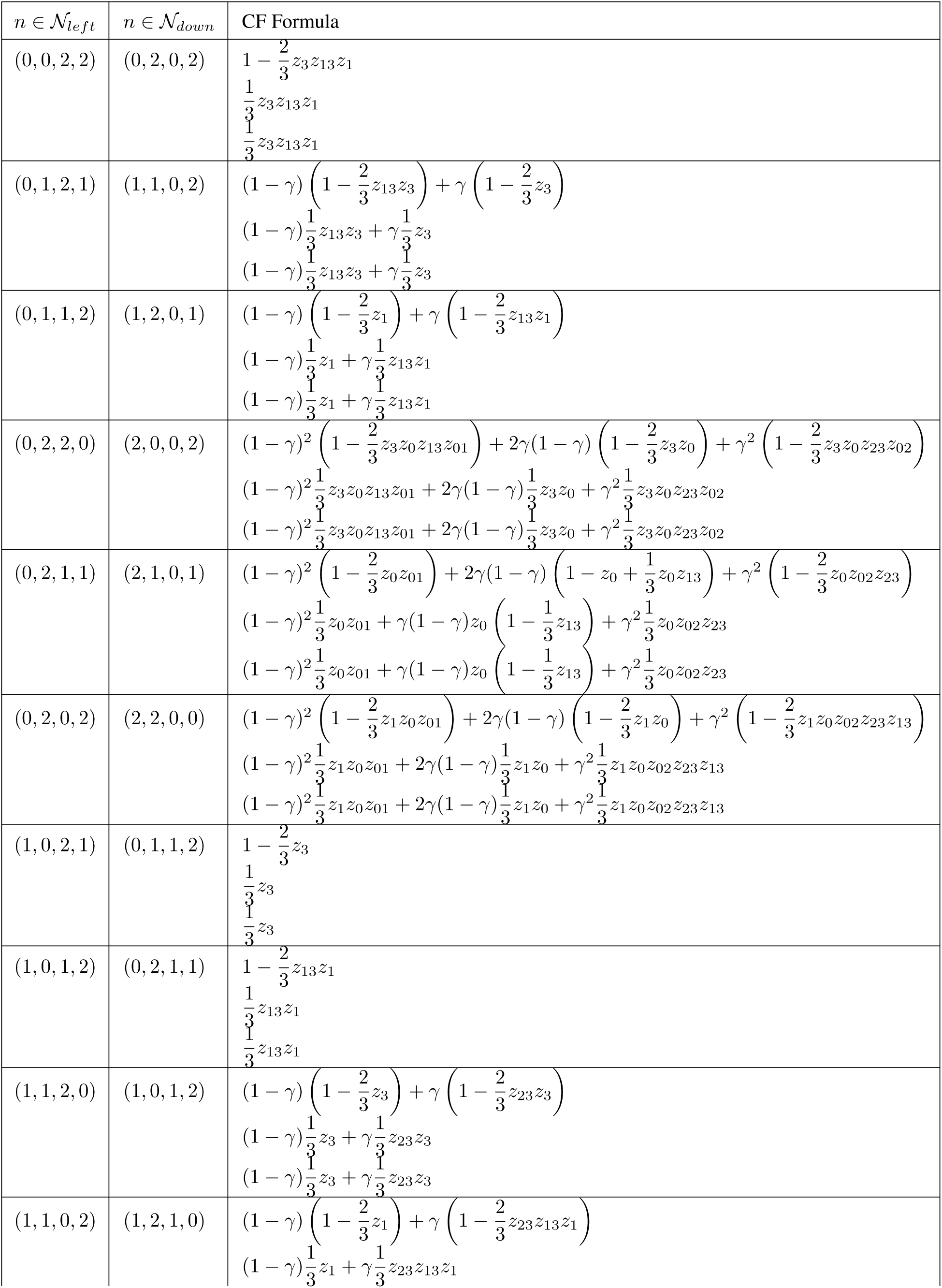
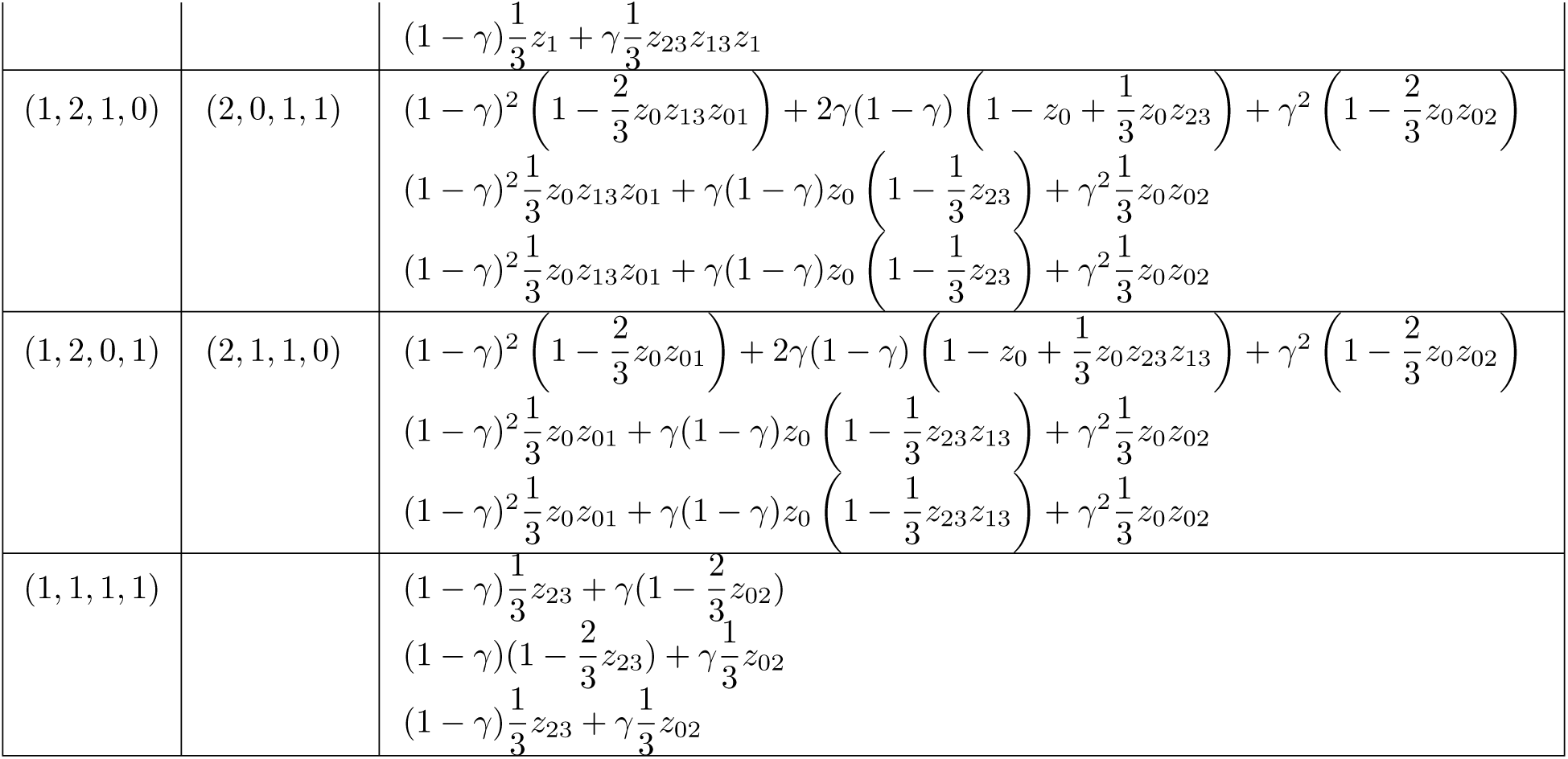
CF equations for *N_left_* for *N* = 2212 (Diamond 3)

### C.4 CF equations for *N_up_* for *N* = 2221 (Diamond 4)

**Table 12:** CF equations for *N_up_* for *N* = 2221 (Diamond 4)

| $n \in \mathcal{N}_{up}$ | $n \in \mathcal{N}_{down}$ | CF Formula |
| --- | --- | --- |
| (0, 0, 2, 2) | (2, 2, 0, 0) | $(1 - \gamma)^2 \left(1 - \frac{2}{3}z_1z_0z_{01}\right) + 2\gamma(1 - \gamma) \left(1 - \frac{2}{3}z_1z_0\right) + \gamma^2 \left(1 - \frac{2}{3}z_1z_0z_{02}z_{23}z_{13}\right)$<br>$(1 - \gamma)^2 \frac{1}{3}z_1z_0z_{01} + 2\gamma(1 - \gamma)\frac{1}{3}z_1z_0 + \gamma^2 \frac{1}{3}z_1z_0z_{02}z_{23}z_{13}$<br>$(1 - \gamma)^2 \frac{1}{3}z_1z_0z_{01} + 2\gamma(1 - \gamma)\frac{1}{3}z_1z_0 + \gamma^2 \frac{1}{3}z_1z_0z_{02}z_{23}z_{13}$ |
| (0, 1, 2, 1) | (1, 2, 1, 0) | $(1 - \gamma) \left(1 - \frac{2}{3}z_1\right) + \gamma \left(1 - \frac{2}{3}z_{23}z_{13}z_1\right)$<br>$(1 - \gamma)\frac{1}{3}z_1 + \gamma\frac{1}{3}z_{23}z_{13}z_1$<br>$(1 - \gamma)\frac{1}{3}z_1 + \gamma\frac{1}{3}z_{23}z_{13}z_1$ |
| (0, 1, 1, 2) | (2, 1, 1, 0) | $(1 - \gamma)^2 \left(1 - \frac{2}{3}z_0z_{01}\right) + 2\gamma(1 - \gamma) \left(1 - z_0 + \frac{1}{3}z_0z_{23}z_{13}\right) + \gamma^2 \left(1 - \frac{2}{3}z_0z_{02}\right)$<br>$(1 - \gamma)^2 \frac{1}{3}z_0z_{01} + \gamma(1 - \gamma)z_0 \left(1 - \frac{1}{3}z_{23}z_{13}\right) + \gamma^2 \frac{1}{3}z_0z_{02}$<br>$(1 - \gamma)^2 \frac{1}{3}z_0z_{01} + \gamma(1 - \gamma)z_0 \left(1 - \frac{1}{3}z_{23}z_{13}\right) + \gamma^2 \frac{1}{3}z_0z_{02}$ |
| (0, 2, 2, 0) | (0, 2, 2, 0) | $1 - \frac{2}{3}z_2z_{23}z_{13}z_1$<br>$\frac{1}{3}z_2z_{23}z_{13}z_1$<br>$\frac{1}{3}z_2z_{23}z_{13}z_1$ |
| (0, 2, 1, 1) | (1, 1, 2, 0) | $(1 - \gamma) \left(1 - \frac{2}{3}z_{13}z_{23}z_2\right) + \gamma \left(1 - \frac{2}{3}z_2\right)$<br>$(1 - \gamma)\frac{1}{3}z_{13}z_{23}z_2 + \gamma\frac{1}{3}z_2$<br>$(1 - \gamma)\frac{1}{3}z_{13}z_{23}z_2 + \gamma\frac{1}{3}z_2$ |
| (0, 2, 0, 2) | (2, 0, 2, 0) | $(1 - \gamma)^2 \left(1 - \frac{2}{3} z_2 z_0 z_{01} z_{13} z_{23}\right) + 2\gamma(1 - \gamma) \left(1 - \frac{2}{3} z_2 z_0\right) + \gamma^2 \left(1 - \frac{2}{3} z_2 z_0 z_{02}\right)$<br>$(1 - \gamma)^2 \frac{1}{3} z_2 z_0 z_{01} z_{13} z_{23} + 2\gamma(1 - \gamma) \frac{1}{3} z_2 z_0 + \gamma^2 \frac{1}{3} z_2 z_0 z_{02}$<br>$(1 - \gamma)^2 \frac{1}{3} z_2 z_0 z_{01} z_{13} z_{23} + 2\gamma(1 - \gamma) \frac{1}{3} z_2 z_0 + \gamma^2 \frac{1}{3} z_2 z_0 z_{02}$ |
| (1, 0, 2, 1) | (1, 2, 0, 1) | $(1 - \gamma) \left(1 - \frac{2}{3} z_1\right) + \gamma \left(1 - \frac{2}{3} z_{13} z_1\right)$<br>$(1 - \gamma) \frac{1}{3} z_1 + \gamma \frac{1}{3} z_{13} z_1$<br>$(1 - \gamma) \frac{1}{3} z_1 + \gamma \frac{1}{3} z_{13} z_1$ |
| (1, 0, 1, 2) | (2, 1, 0, 1) | $(1 - \gamma)^2 \left(1 - \frac{2}{3} z_0 z_{01}\right) + 2\gamma(1 - \gamma) \left(1 - z_0 + \frac{1}{3} z_0 z_{13}\right) + \gamma^2 \left(1 - \frac{2}{3} z_0 z_{02} z_{23}\right)$<br>$(1 - \gamma)^2 \frac{1}{3} z_0 z_{01} + \gamma(1 - \gamma) z_0 \left(1 - \frac{1}{3} z_{13}\right) + \gamma^2 \frac{1}{3} z_0 z_{02} z_{23}$<br>$(1 - \gamma)^2 \frac{1}{3} z_0 z_{01} + \gamma(1 - \gamma) z_0 \left(1 - \frac{1}{3} z_{13}\right) + \gamma^2 \frac{1}{3} z_0 z_{02} z_{23}$ |
| (1, 1, 2, 0) | (0, 2, 1, 1) | $1 - \frac{2}{3} z_{13} z_1$<br>$\frac{1}{3} z_{13} z_1$<br>$\frac{1}{3} z_{13} z_1$ |
| (1, 1, 0, 2) | (2, 0, 1, 1) | $(1 - \gamma)^2 \left(1 - \frac{2}{3} z_0 z_{13} z_{01}\right) + 2\gamma(1 - \gamma) \left(1 - z_0 + \frac{1}{3} z_0 z_{23}\right) + \gamma^2 \left(1 - \frac{2}{3} z_0 z_{02}\right)$<br>$(1 - \gamma)^2 \frac{1}{3} z_0 z_{13} z_{01} + \gamma(1 - \gamma) z_0 \left(1 - \frac{1}{3} z_{23}\right) + \gamma^2 \frac{1}{3} z_0 z_{02}$<br>$(1 - \gamma)^2 \frac{1}{3} z_0 z_{13} z_{01} + \gamma(1 - \gamma) z_0 \left(1 - \frac{1}{3} z_{23}\right) + \gamma^2 \frac{1}{3} z_0 z_{02}$ |
| (1, 2, 1, 0) | (0, 1, 2, 1) | $1 - \frac{2}{3} z_{23} z_2$<br>$\frac{1}{3} z_{23} z_2$<br>$\frac{1}{3} z_{23} z_2$ |
| (1, 2, 0, 1) | (1, 0, 2, 1) | $(1 - \gamma) \left(1 - \frac{2}{3} z_{23} z_2\right) + \gamma \left(1 - \frac{2}{3} z_2\right)$<br>$(1 - \gamma) \frac{1}{3} z_{23} z_2 + \gamma \frac{1}{3} z_2$<br>$(1 - \gamma) \frac{1}{3} z_{23} z_2 + \gamma \frac{1}{3} z_2$ |
| (1, 1, 1, 1) | | $(1 - \gamma) \left(1 - \frac{2}{3} z_{02}\right) + \gamma \frac{1}{3} z_{01}$<br>$(1 - \gamma) \frac{1}{3} z_{02} + \gamma \left(1 - \frac{2}{3} z_{01}\right)$<br>$(1 - \gamma) \frac{1}{3} z_{02} + \gamma \frac{1}{3} z_{01}$ |

## D Theorems and proofs for generic identifiability of 4-node cycles in semi-directed level-1 phylogenetic networks

**Theorem 3.** Let N_down_ be a semi-directed level-1 6-taxon phylogenetic network with one hybridization event producing a hybridization cycle with 4 nodes. Without loss of generality, let the taxa be partitioned among clades as n_0_ = 1*, n*_1_ = 2*, n*_2_ = 1*, n*_3_ = 2 *(Figure 3). Let the hybrid node be ancestral to the clade n*_0_*. Let N_left_ be a semi-directed level-1 6-taxon phylogenetic network with one hybridization event producing a hybridization cycle with 4 nodes such that the unrooted version of N_left_ agrees with the unrooted version of N_down_. Let the hybrid node in the hybridization cycle in N_left_ be ancestral to the clade n*_1_*. Then, N_down_ and N_left_ are generically identifiable if t*_1_ *<* ∞*, t*_13_ *>* 0*, t*_3_ *<* ∞*, and γ* ∈ (0, 1).

*Proof.* The structure of the proof is the same as for Theorem 2, but we repeat it for completeness. Let *CF* (*N_down_,* ***z***, ***γ***) be the system of CF polynomial equations defined by *N_down_* and let *CF* (*N_left_,* ***z***^′^, ***γ***^′^) be the system of CF polynomial equations defined by *N_left_*. Both systems of equations can be found in the Appendix.

Let P = {*p*(***z****, **γ***) − *q*(***z***^′^*, **γ***^′^) : *p*(***z****, **γ***) ∈ *CF* (*N_down_, **z**, **γ***)*, q*(***z***^′^*, **γ***^′^) ∈ *CF* (*N_left_, **z***^′^*, **γ***^′^)} be the set of polynomial equations resulting from matching *CF* (*N_down_, **z**, **γ***) to *CF* (*N_left_, **z***^′^*, **γ***^′^).

Let C[(***z****, **γ**, **z***^′^*, **γ***^′^) : ***z*** ∈ [0, ∞)*^n^*^−3^*, **γ*** ∈ [0, 1]*^h^, **z***^′^ ∈ [0, ∞)*^n^*^−3^*, **γ***^′^ ∈ [0, 1]*^h^*] be the set of all polynomials on the (***z****, **γ**, **z***^′^*, **γ***^′^) variables.

The resulting ideal is given by

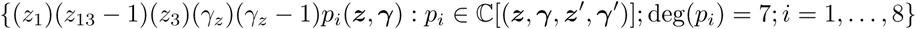

As the polynomials 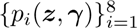 have Lebesgue measure zero, *N_down_* and *N_left_* are not generically identifiable in the subset of parameter space corresponding to {*z*_1_ = 0} ∪ {*z*_13_ = 1} ∪ {*z*_3_ = 0} ∪ {*γ* = 0} ∪ {*γ* = 1}.

**Theorem 4.** Let N_down_ be a semi-directed level-1 6-taxon phylogenetic network with one hybridization event producing a hybridization cycle with 4 nodes. Without loss of generality, let the taxa be partitioned among clades as n_0_ = 1*, n*_1_ = 2*, n*_2_ = 1*, n*_3_ = 2 *(Figure 3). Let the hybrid node be ancestral to the clade n*_0_*. Let N_up_ be a semi-directed level-1 6-taxon phylogenetic network with one hybridization event producing a hybridization cycle with 4 nodes such that the unrooted version of N_up_ agrees with the unrooted version of N_down_. Let the hybrid node in the hybridization cycle in N_up_ be ancestral to the clade n*_3_*. Then, N_down_ and N_up_ are generically identifiable if t*_1_ *<* ∞*, t*_13_ *>* 0*, t*_3_ *<* ∞*, and γ* ∈ (0, 1).

*Proof.* The structure of the proof is the same as for Theorem 3, but we repeat it for completeness. Let *CF* (*N_down_,* ***z***, ***γ***) be the system of CF polynomial equations defined by *N_down_* and let *CF* (*N_up_,* ***z***^′^, ***γ***^′^) be the system of CF polynomial equations defined by *N_up_*. Both systems of equations can be found in the Appendix.

Let P = {*p*(***z****, **γ***) − *q*(***z***^′^*, **γ***^′^) : *p*(***z****, **γ***) ∈ *CF* (*N_down_, **z**, **γ***)*, q*(***z***^′^*, **γ***^′^) ∈ *CF* (*N_up_, **z***^′^*, **γ***^′^)} be the set of polynomial equations resulting from matching *CF* (*N_down_, **z**, **γ***) to *CF* (*N_up_, **z***^′^*, **γ***^′^).

Let C[(***z****, **γ**, **z***^′^*, **γ***^′^) : ***z*** ∈ [0, ∞)*^n^*^−3^*, **γ*** ∈ [0, 1]*^h^, **z***^′^ ∈ [0, ∞)*^n^*^−3^*, **γ***^′^ ∈ [0, 1]*^h^*] be the set of all polynomials on the (***z****, **γ**, **z***^′^*, **γ***^′^) variables.

The resulting ideal is given by

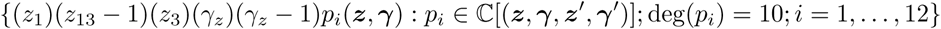

As the polynomials 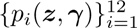 have Lebesgue measure zero, *N_down_* and *N_up_* are not generically identifiable in the subset of parameter space corresponding to {*z*_1_ = 0} ∪ {*z*_13_ = 1} ∪ {*z*_3_ = 0} ∪ {*γ* = 0} ∪ {*γ* = 1}.

## E Subset of CF equations used in Macaulay2

We present below the table with the major CF for all 4-taxon subsets, except (1,1,1,1) for which we present all equations. All equations are multiplied by 3 for conciseness. All Macaulay2 scripts can be found in https://github.com/gtiley/diamoNd-ideNtifiability/tree/maiN/scripts/macaulay2.

**Table 13:**
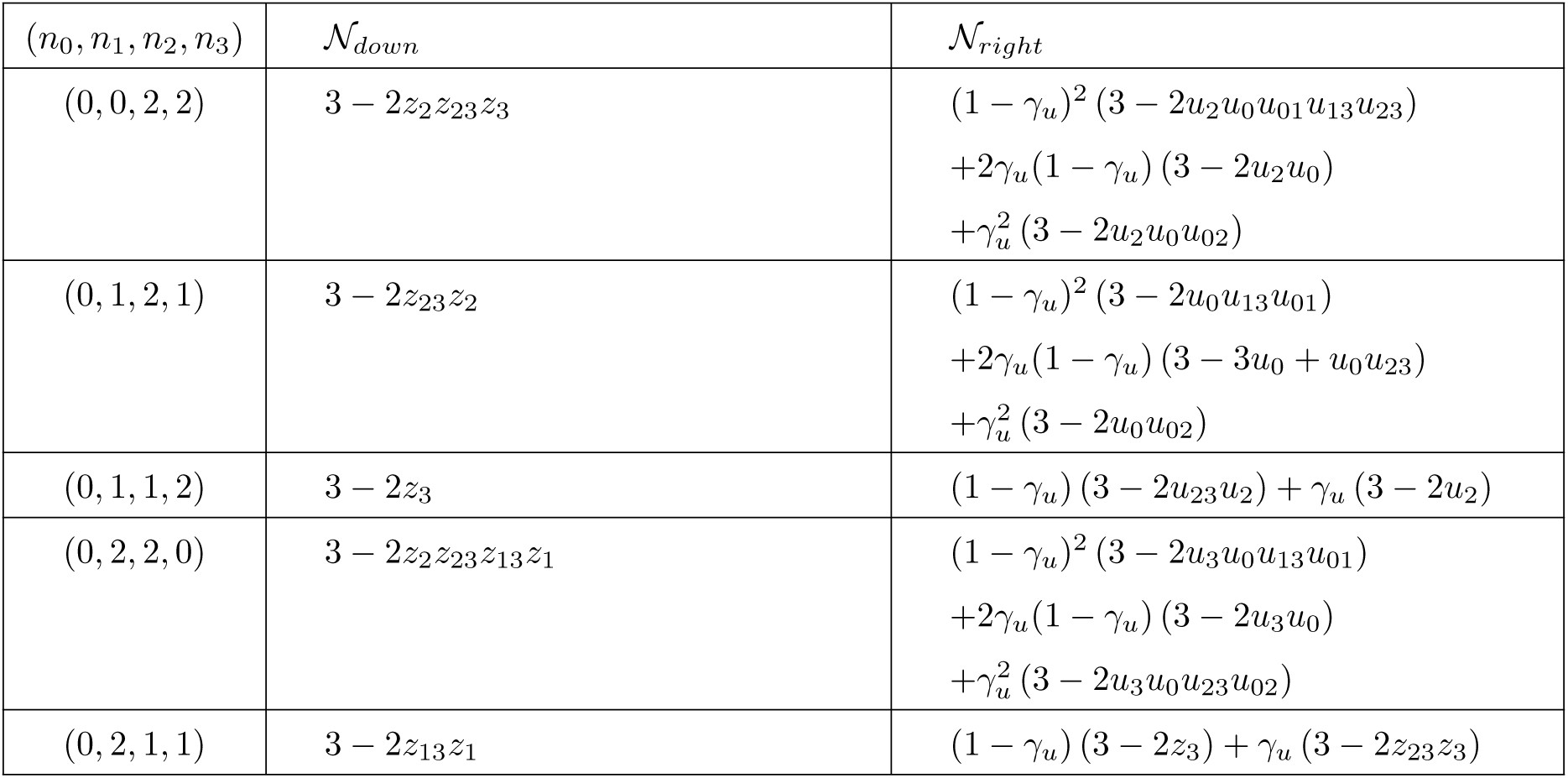
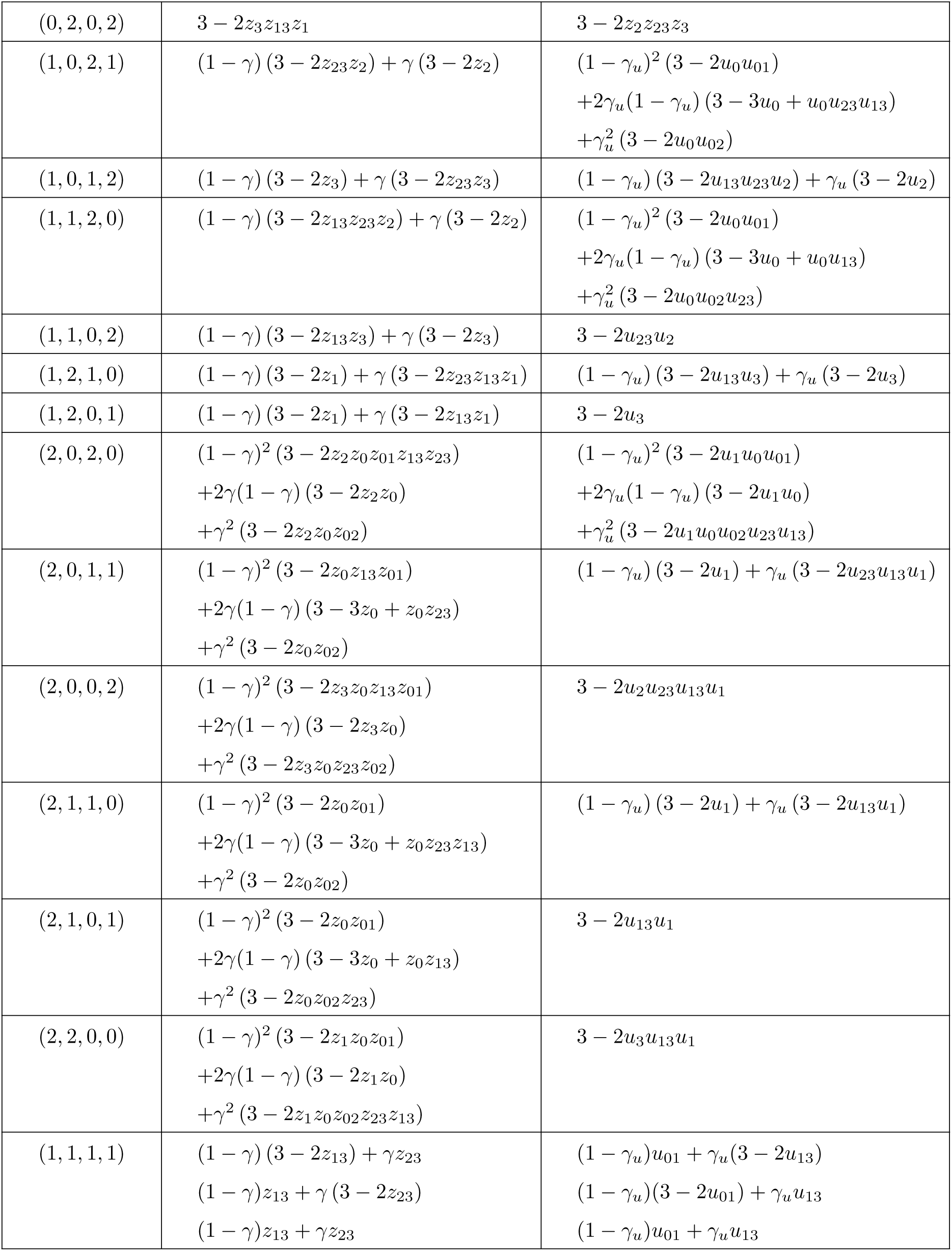
Subset of CF equations used in Macaulay2 (Part 1)

**Figure 11:**
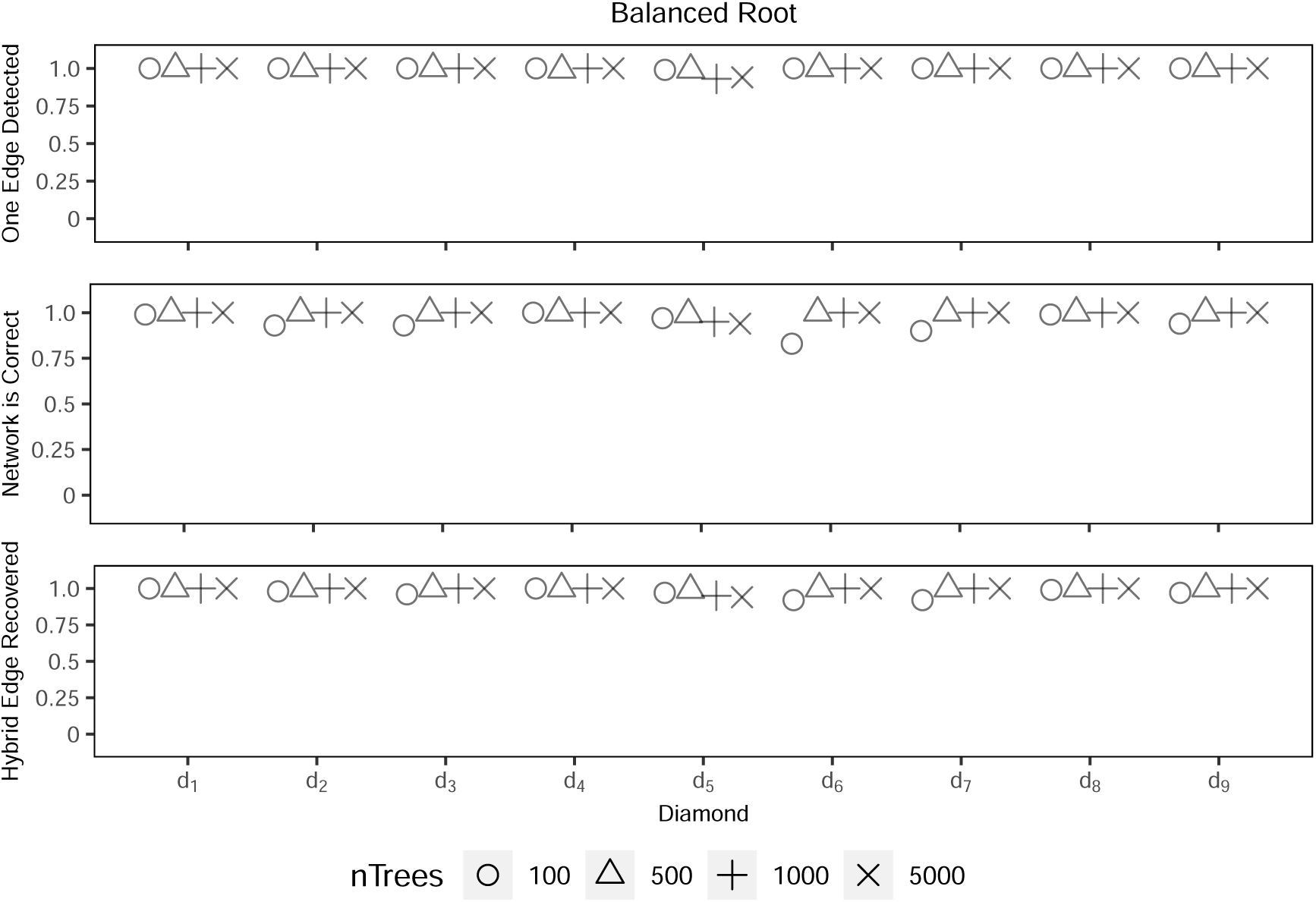
SNaQ results for root 1. *One Edge Detected* is the proportion out of 100 replicates that 1 reticulate edge was correctly inferred by the pseudolikehood scores. *Network is Correct* is the proportion of 100 replicates where the estimated topology when allowing only one reticulation event is identical to the true topology. *Hybrid Edge Recovered* is the proportion of 100 replicates where the correct reticulation edge was inferred, regardless if other parts of the estimated network were incorrect, when allowing only one reticulation event. Diamond d1 through d9 correspond to Fig. 1e.

**Table 14:**
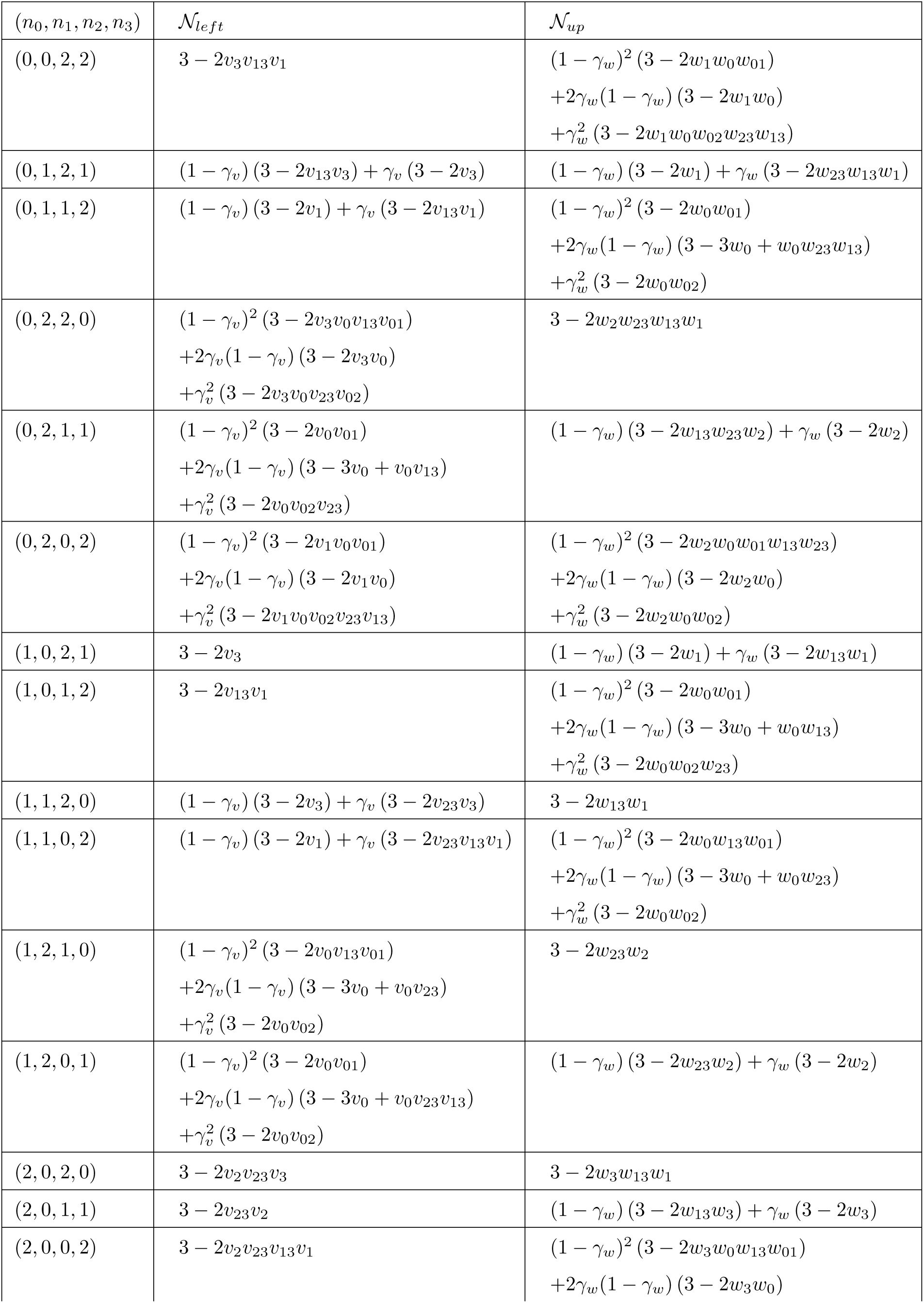
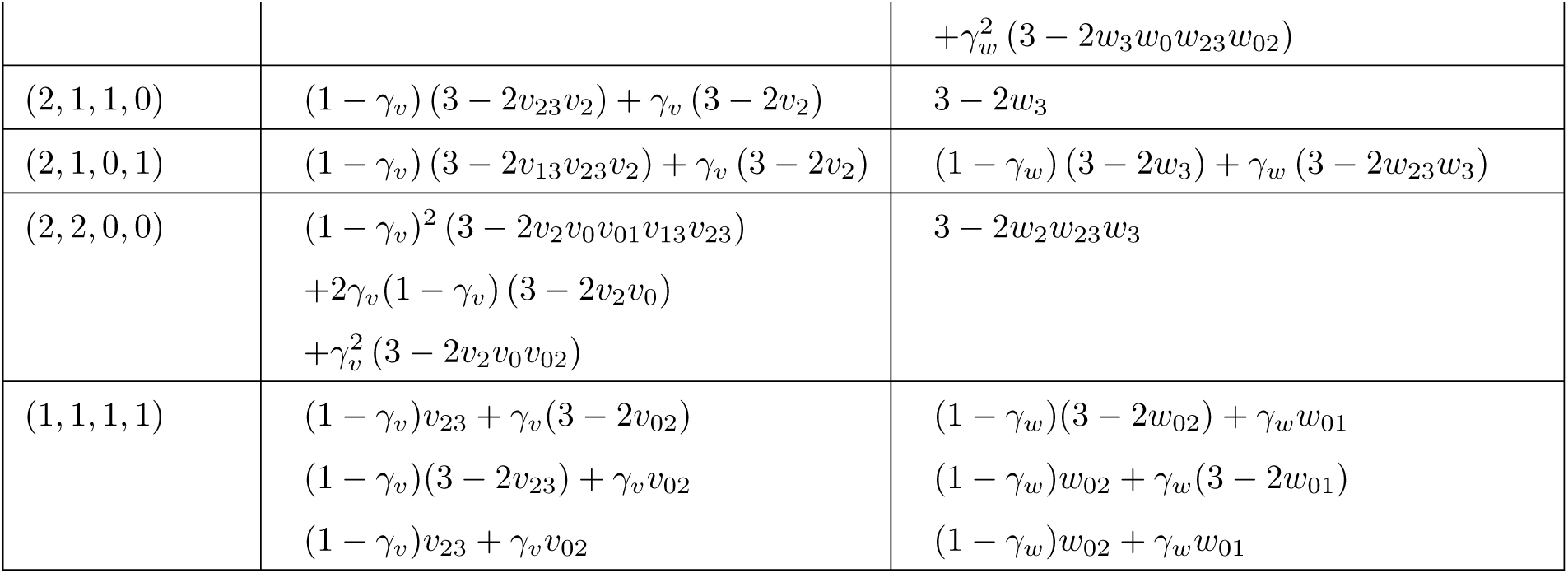
Subset of CF equations used in Macaulay2 (Part 2)

## F Simulation figures and tables

**Figure 12:**
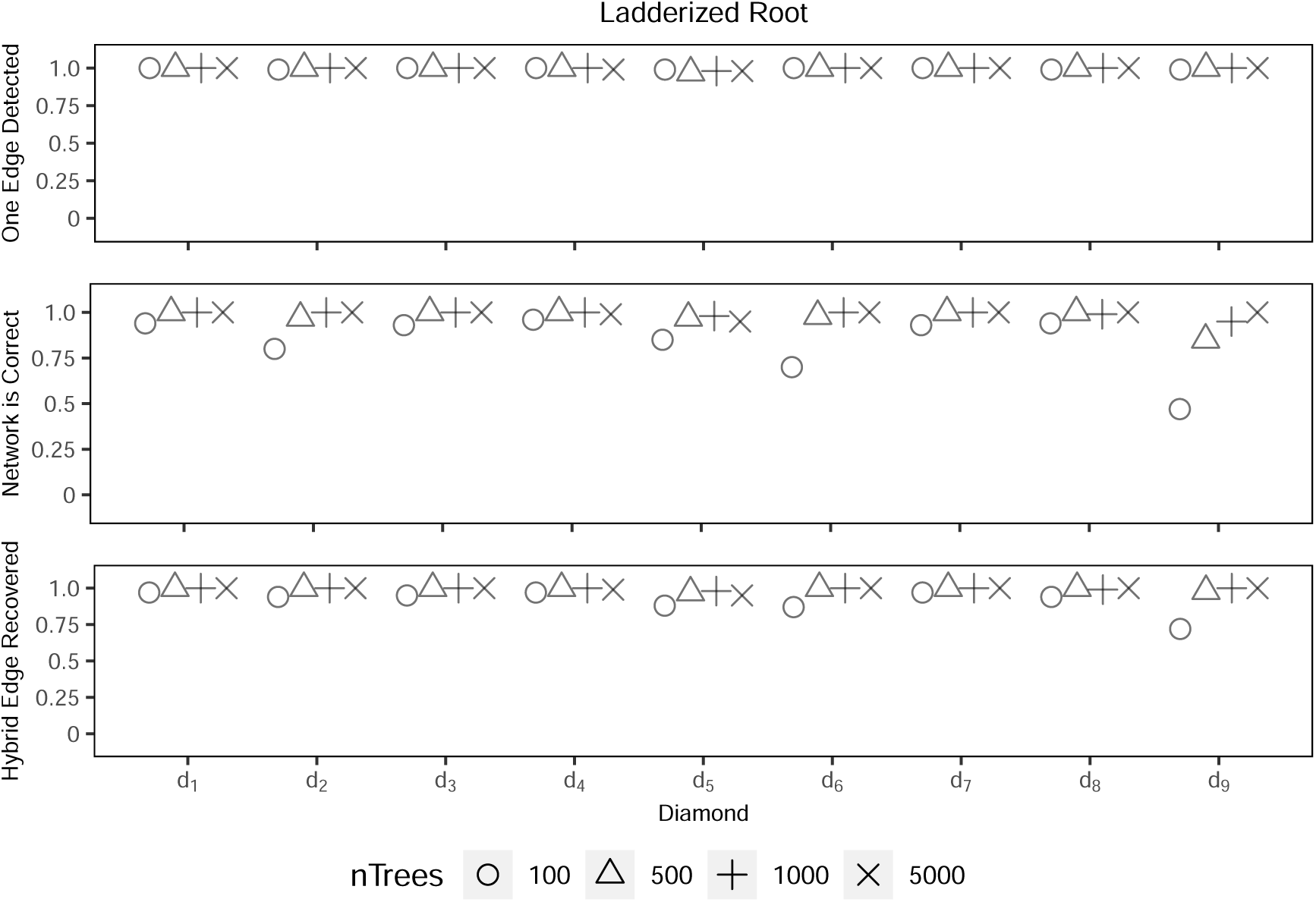
SNaQ results for root 2. *One Edge Detected* is the proportion out of 100 replicates that 1 reticulate edge was correctly inferred by the pseudolikehood scores. *Network is Correct* is the proportion of 100 replicates where the estimated topology when allowing only one reticulation event is identical to the true topology. *Hybrid Edge Recovered* is the proportion of 100 replicates where the correct reticulation edge was inferred, regardless if other parts of the estimated network were incorrect, when allowing only one reticulation event. Diamond d1 through d9 correspond to Fig. 1e.

**Figure 13:**
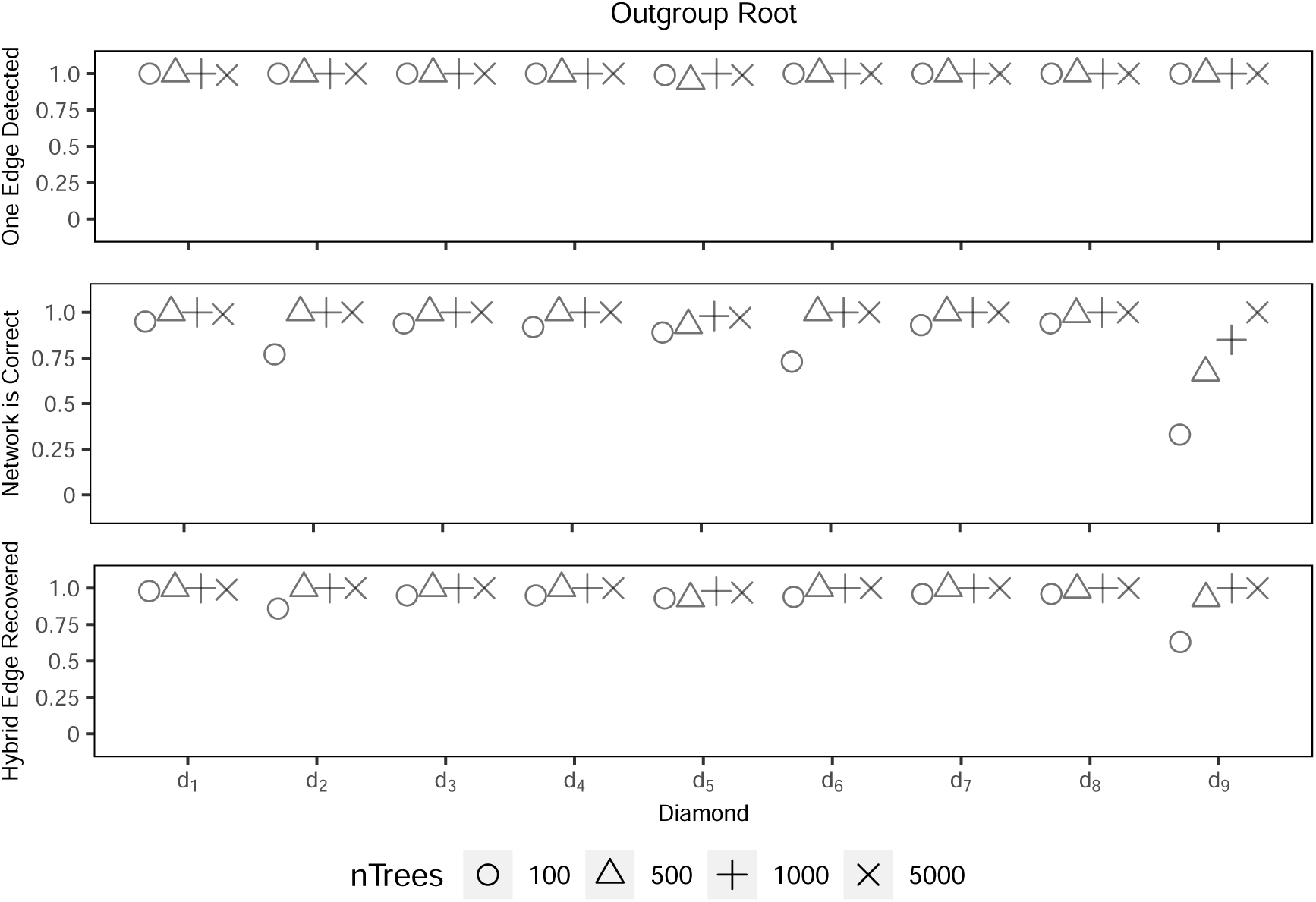
SNaQ results for root 3. *One Edge Detected* is the proportion out of 100 replicates that 1 reticulate edge was correctly inferred by the pseudolikehood scores. *Network is Correct* is the proportion of 100 replicates where the estimated topology when allowing only one reticulation event is identical to the true topology. *Hybrid Edge Recovered* is the proportion of 100 replicates where the correct reticulation edge was inferred, regardless if other parts of the estimated network were incorrect, when allowing only one reticulation event. Diamond d1 through d9 correspond to Fig. 1e.

**Figure 14:**
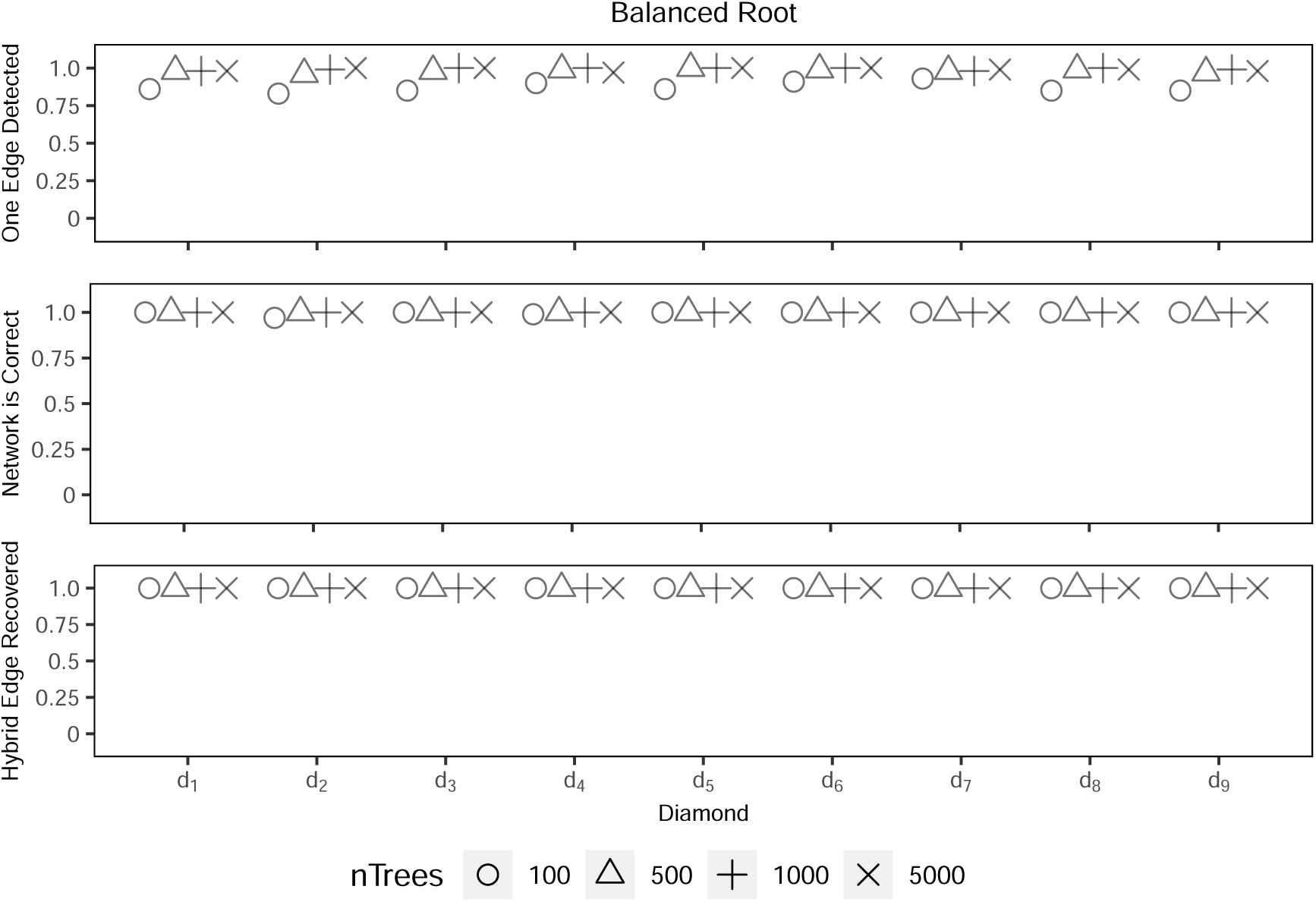
PhyloNet results for root 1. *One Edge Detected* is the proportion out of 100 replicates that 1 reticulate edge was correctly inferred by the pseudolikehood scores. *Network is Correct* is the proportion of 100 replicates where the estimated topology when allowing only one reticulation event is identical to the true topology. *Hybrid Edge Recovered* is the proportion of 100 replicates where the correct reticulation edge was inferred, regardless if other parts of the estimated network were incorrect, when allowing only one reticulation event. Diamond d1 through d9 correspond to Fig. 1e.

**Figure 15:**
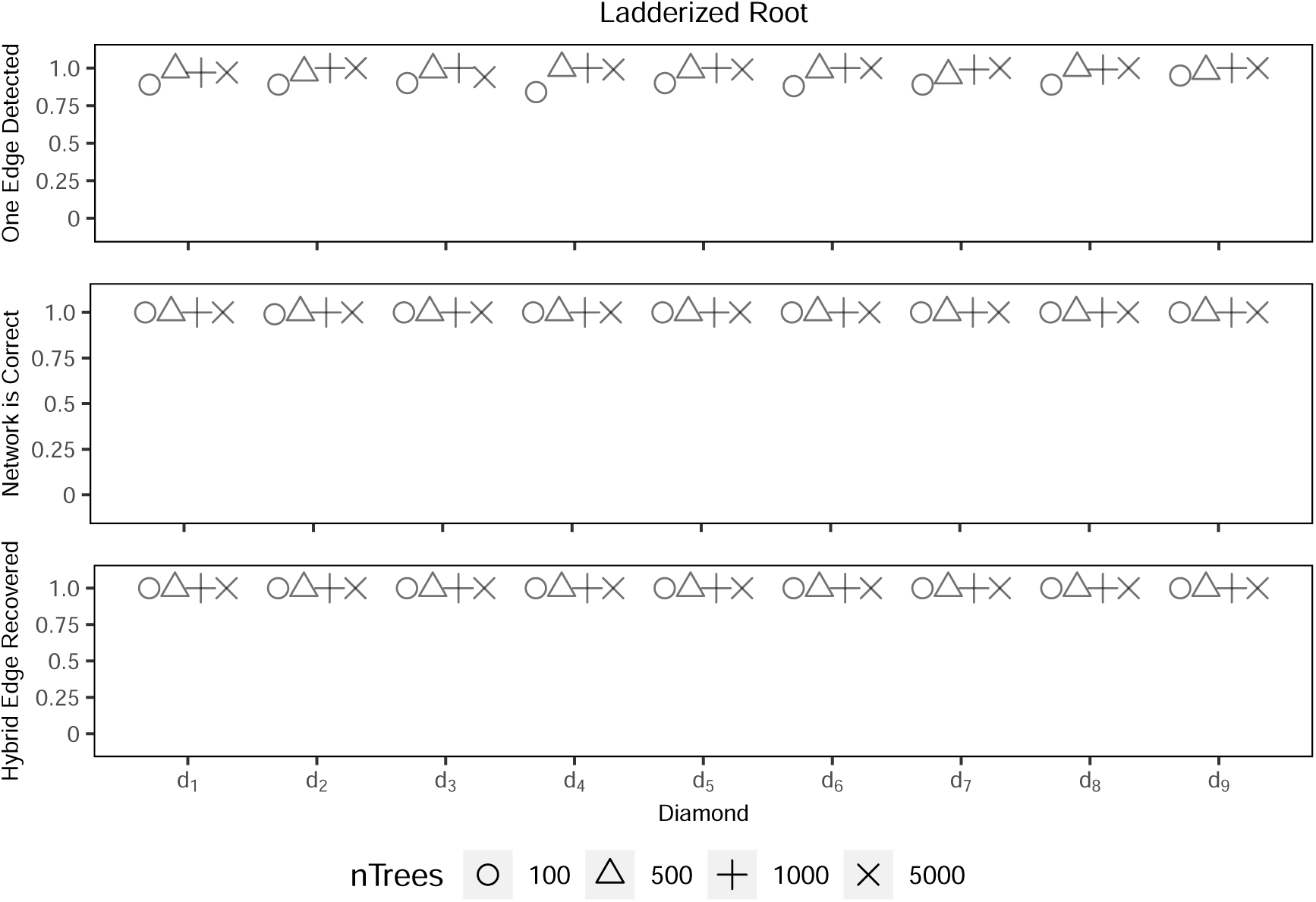
PhyloNet results for root 2. *One Edge Detected* is the proportion out of 100 replicates that 1 reticulate edge was correctly inferred by the pseudolikehood scores. *Network is Correct* is the proportion of 100 replicates where the estimated topology when allowing only one reticulation event is identical to the true topology. *Hybrid Edge Recovered* is the proportion of 100 replicates where the correct reticulation edge was inferred, regardless if other parts of the estimated network were incorrect, when allowing only one reticulation event. Diamond d1 through d9 correspond to Fig. 1e.

**Figure 16:**
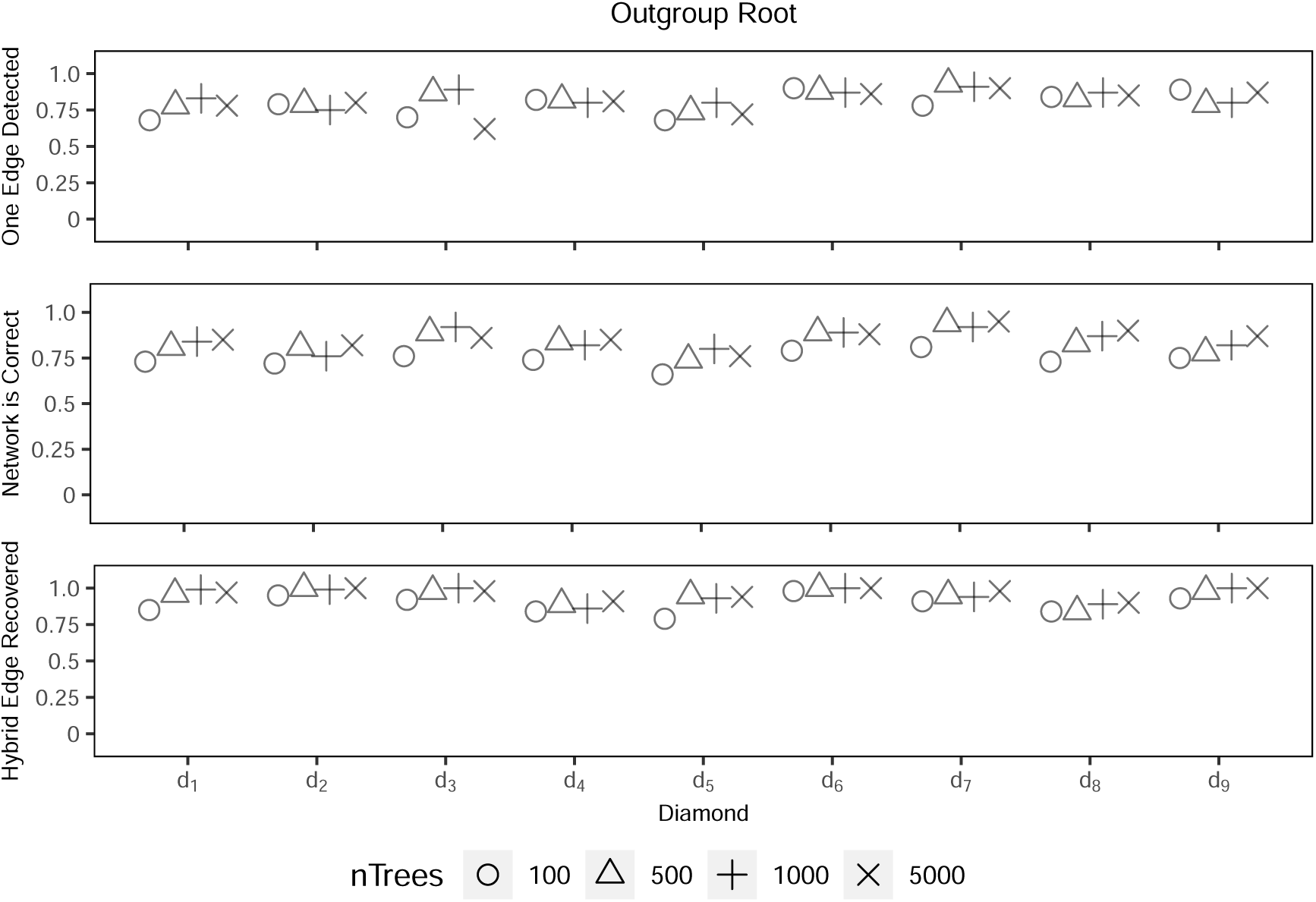
PhyloNet results for root 3. *One Edge Detected* is the proportion out of 100 replicates that 1 reticulate edge was correctly inferred by the pseudolikehood scores. *Network is Correct* is the proportion of 100 replicates where the estimated topology when allowing only one reticulation event is identical to the true topology. *Hybrid Edge Recovered* is the proportion of 100 replicates where the correct reticulation edge was inferred, regardless if other parts of the estimated network were incorrect, when allowing only one reticulation event. Diamond d1 through d9 correspond to Fig. 1e.

**Figure 17:**
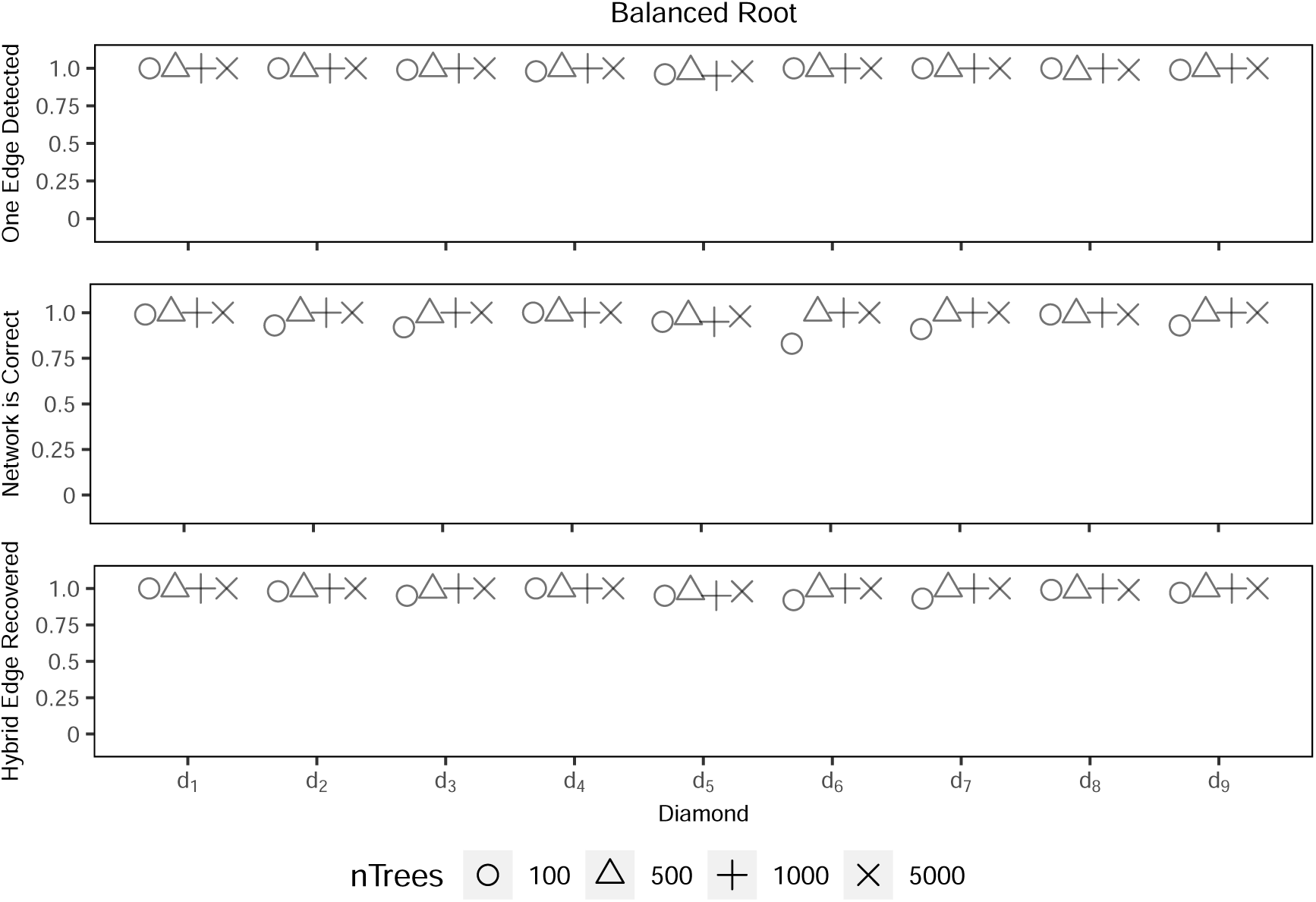
SNaQ results for root 1 with random outgroups for 10% of gene trees.

**Figure 18:**
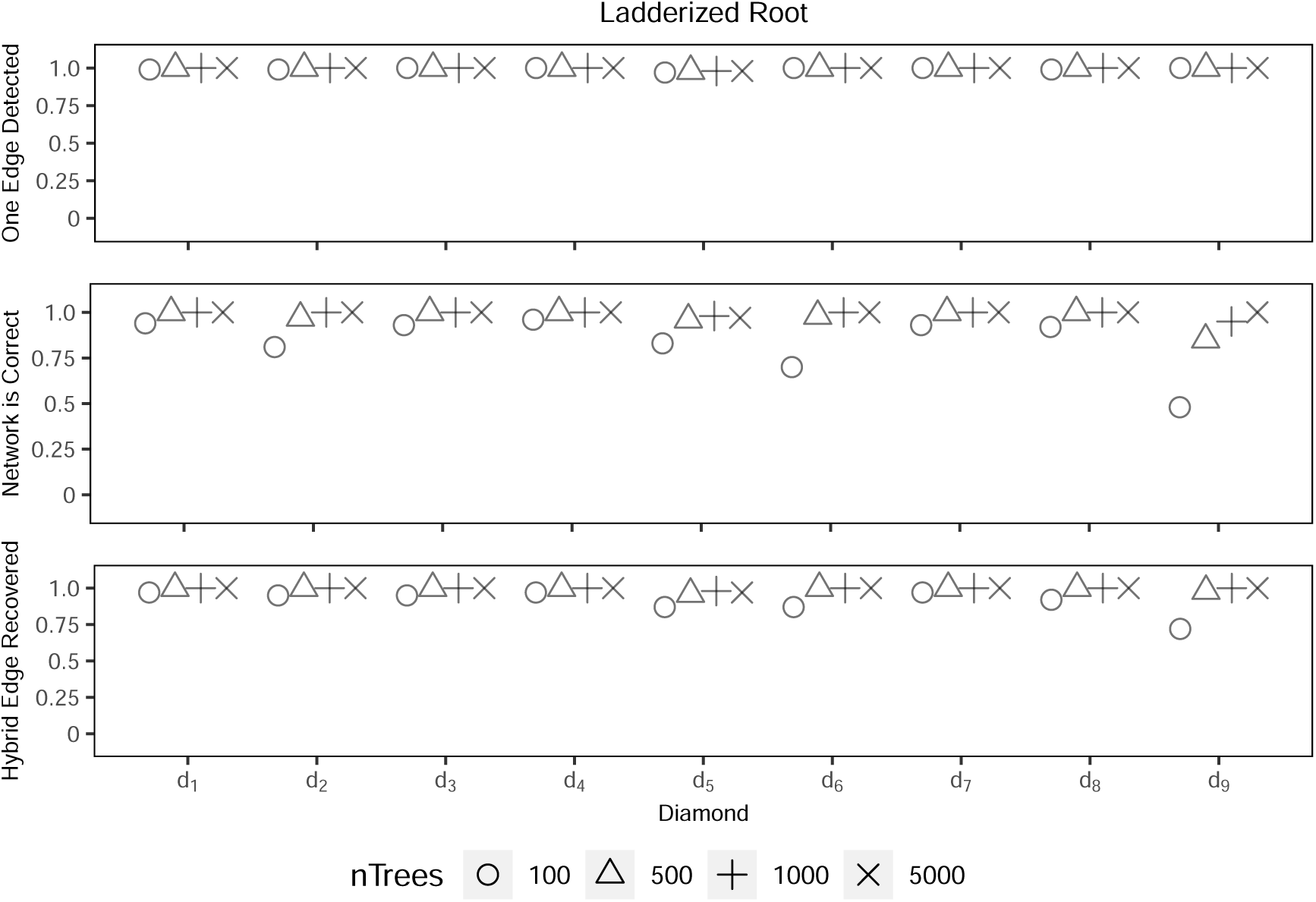
SNaQ results for root 2 with random outgroups for 10% of gene trees.

**Figure 19:**
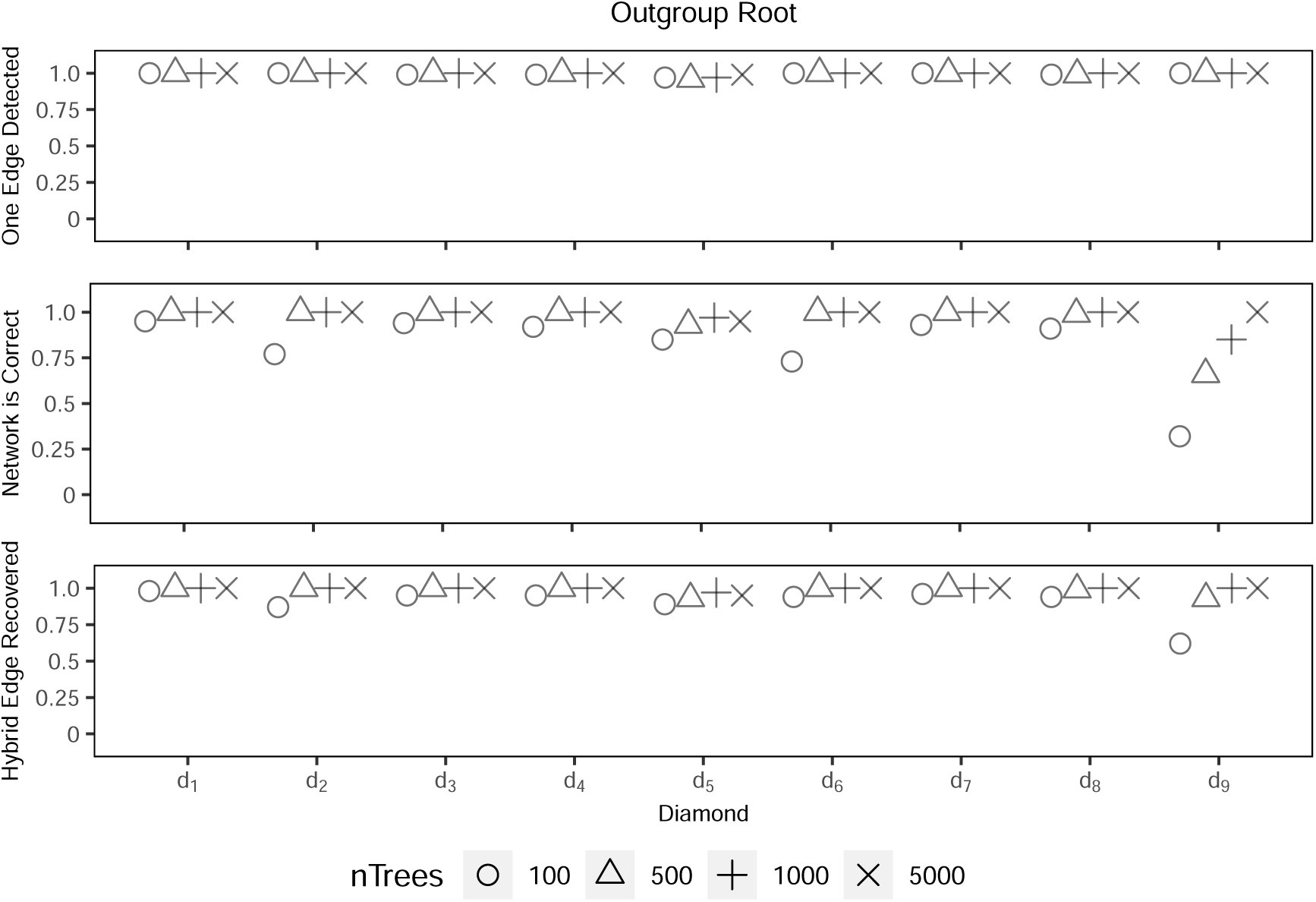
SNaQ results for root 3 with random outgroups for 10% of gene trees.

**Figure 20:**
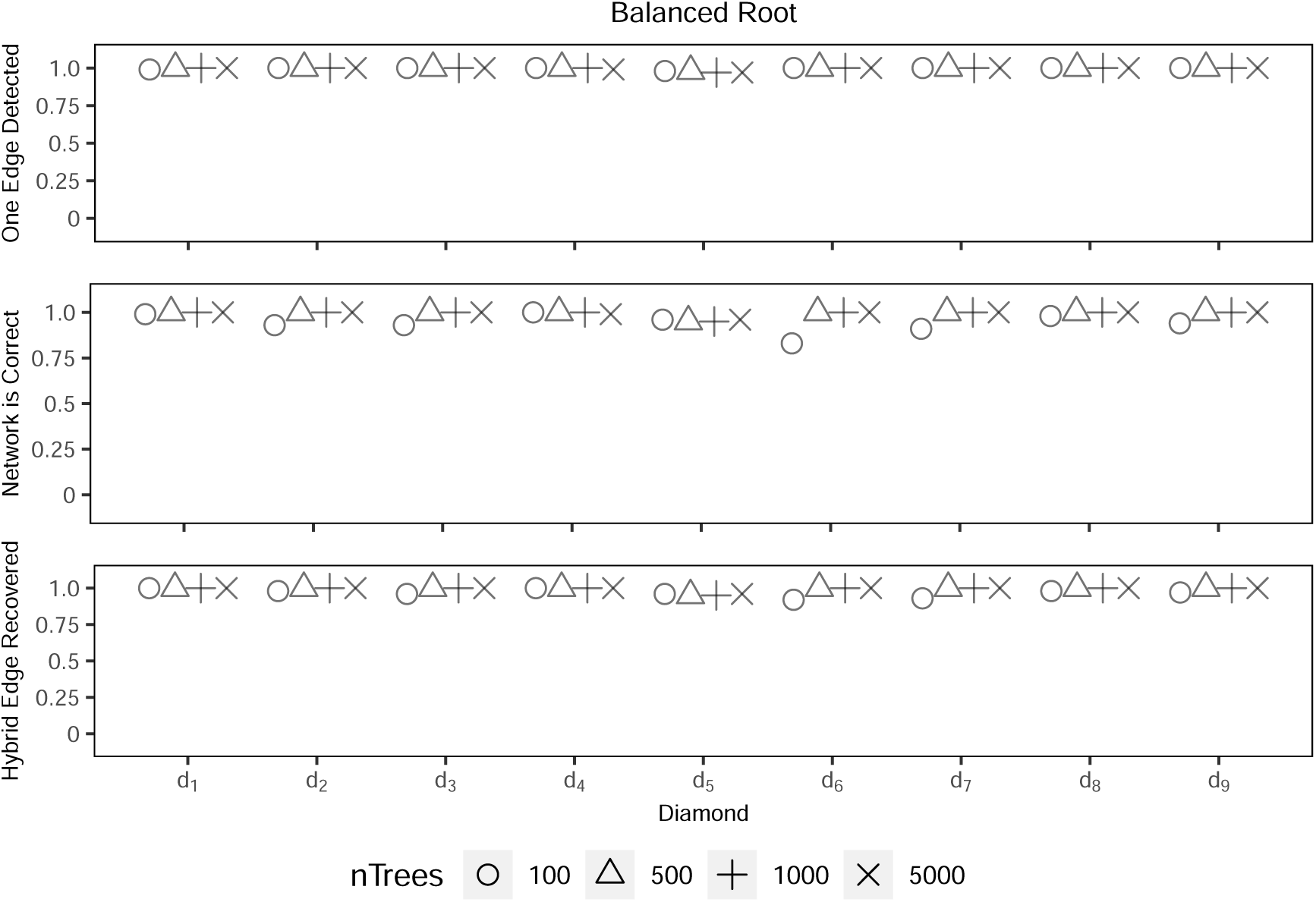
SNaQ results for root 1 with random outgroups for 30% of gene trees.

**Figure 21:**
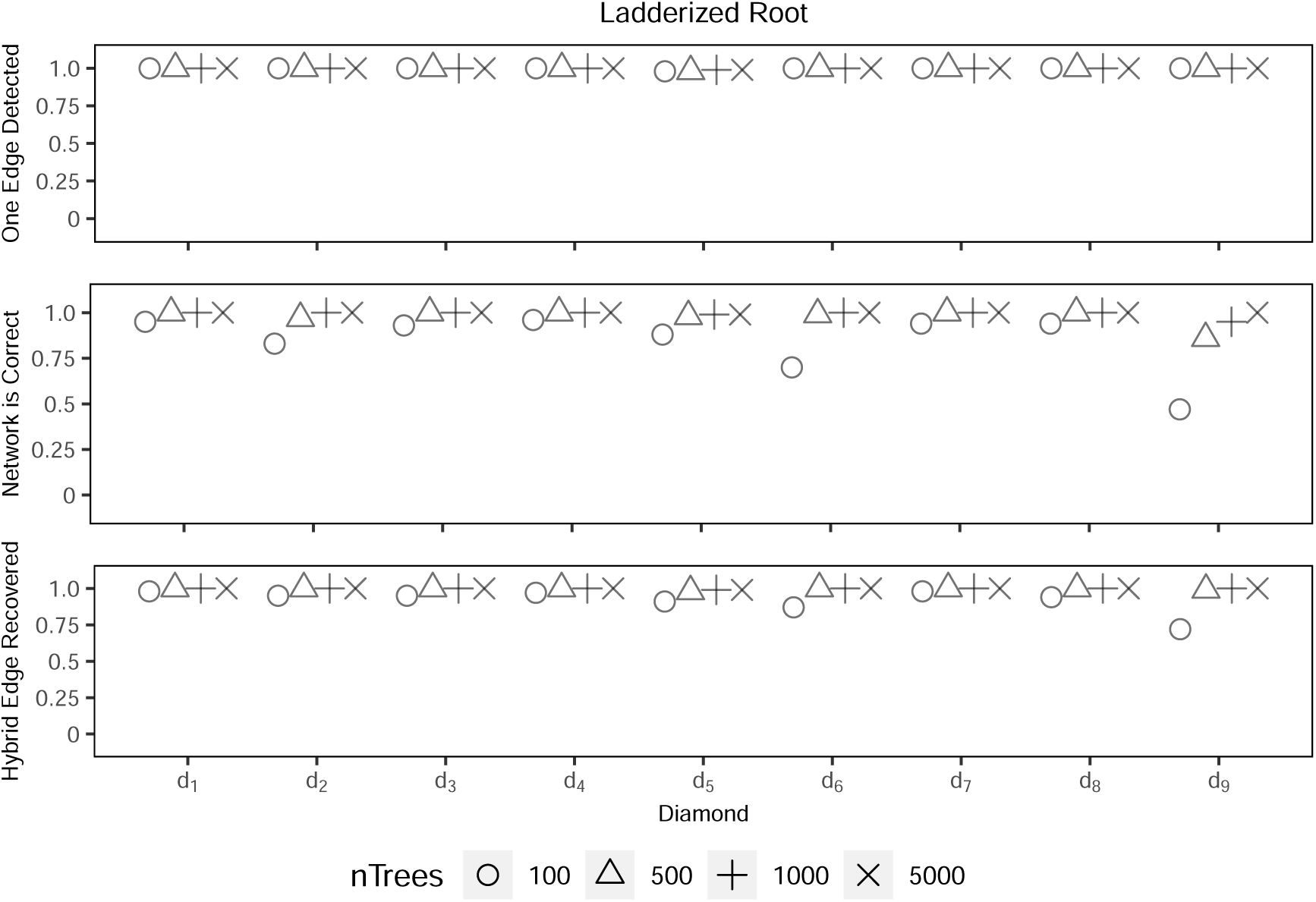
SNaQ results for root 2 with random outgroups for 30% of gene trees.

**Figure 22:**
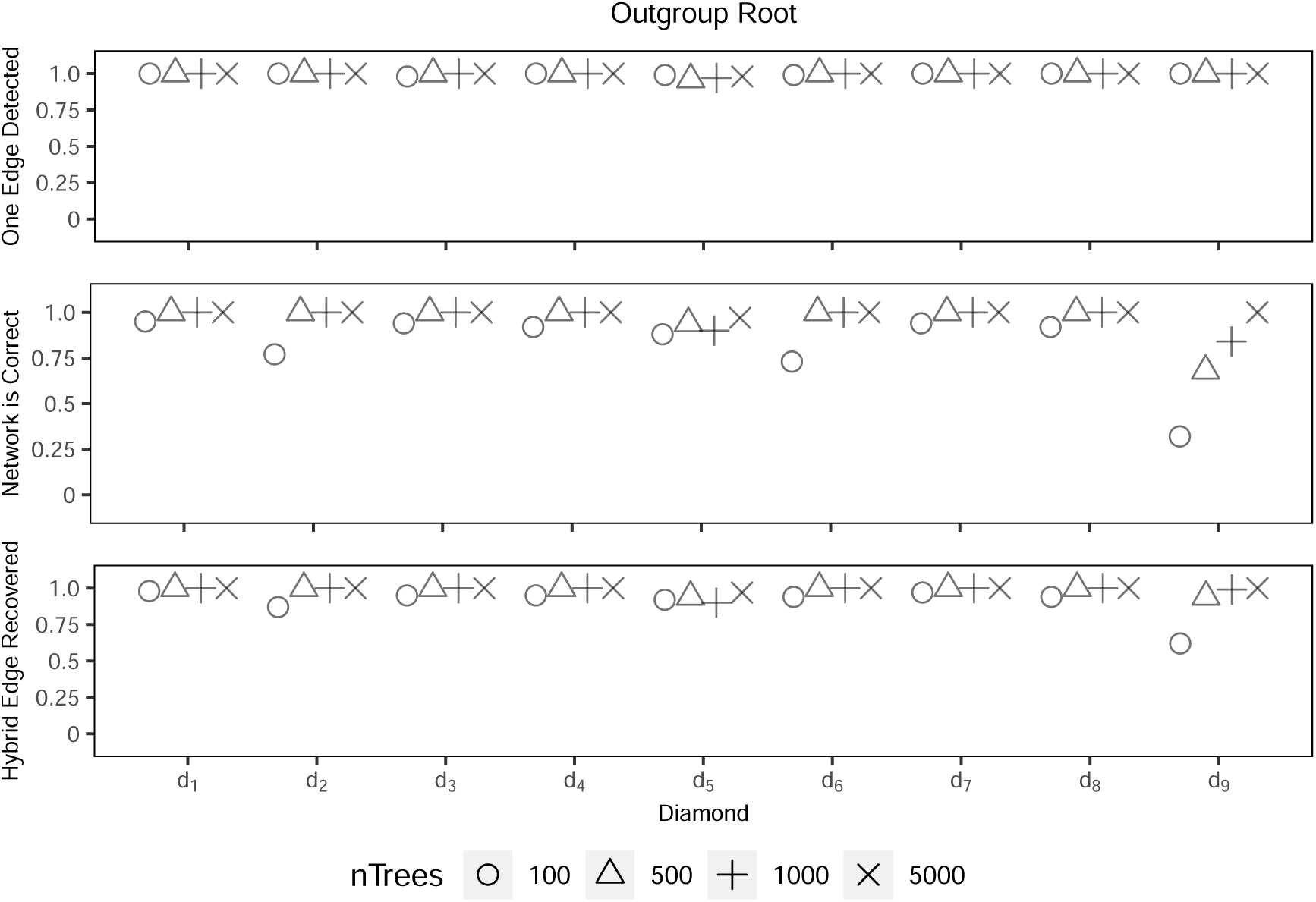
SNaQ results for root 3 with random outgroups for 30% of gene trees.

**Figure 23:**
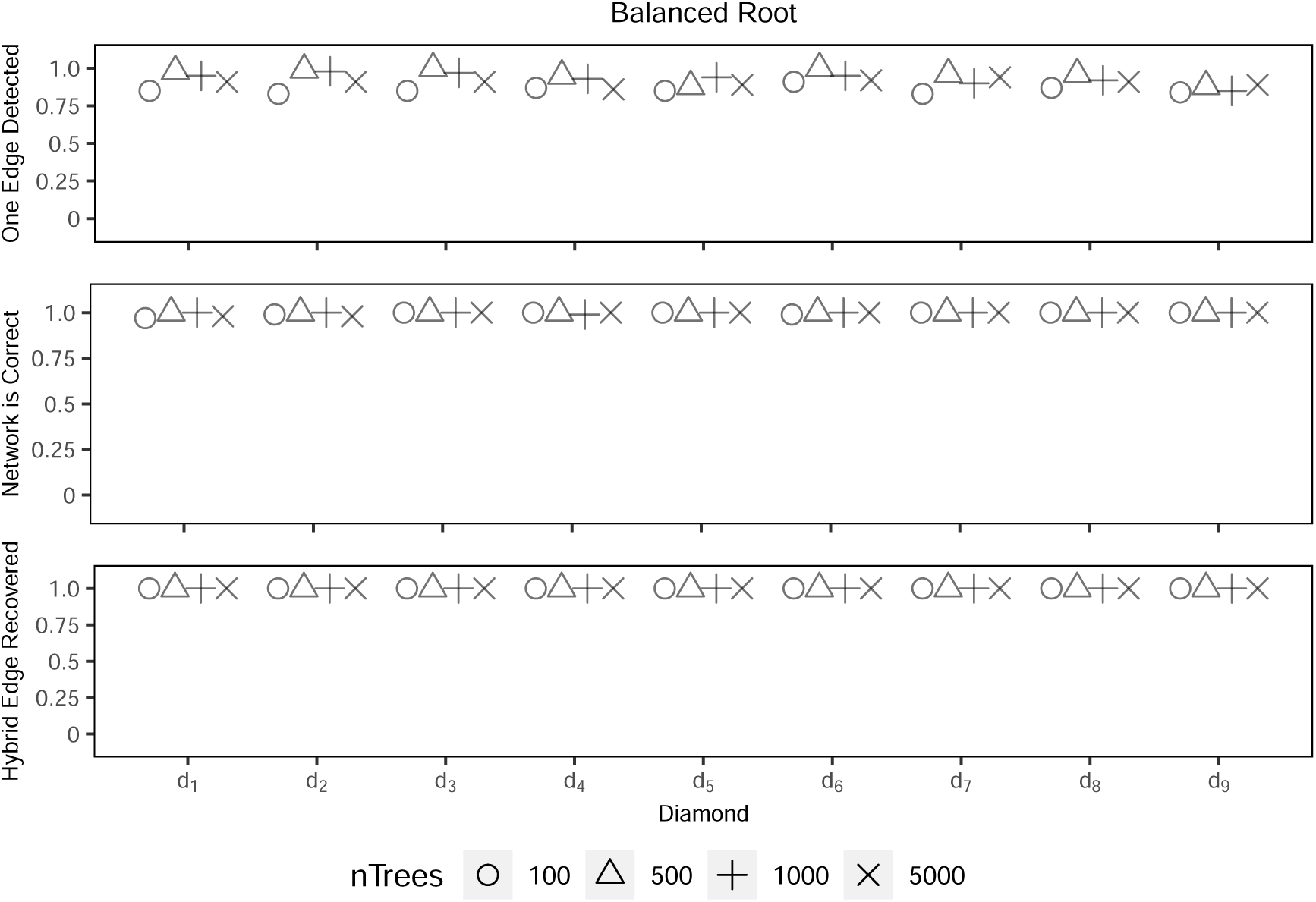
PhyloNet results for root 1 with random outgroups for 10% of gene trees.

**Figure 24:**
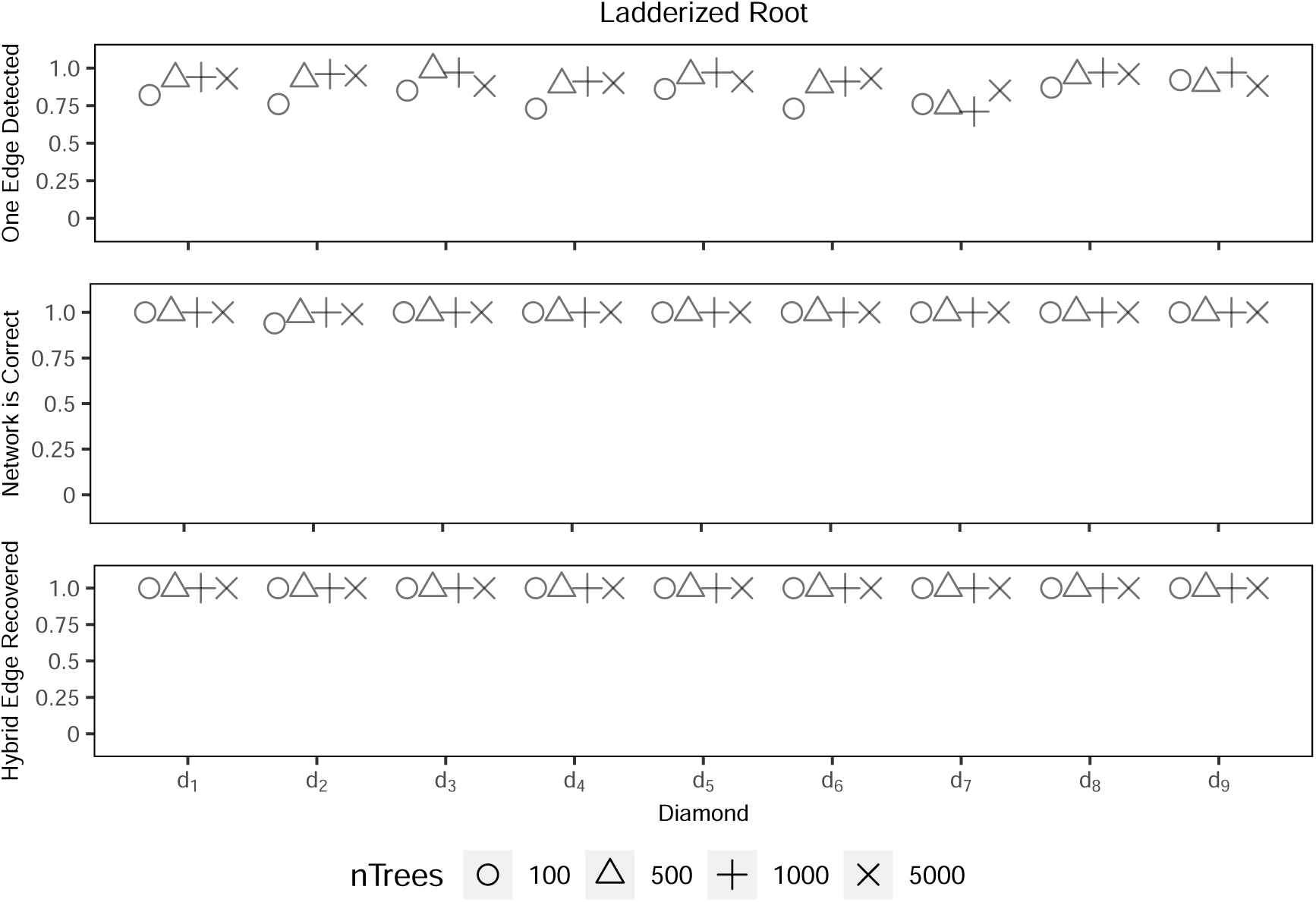
PhyloNet results for root 2 with random outgroups for 10% of gene trees.

**Figure 25:**
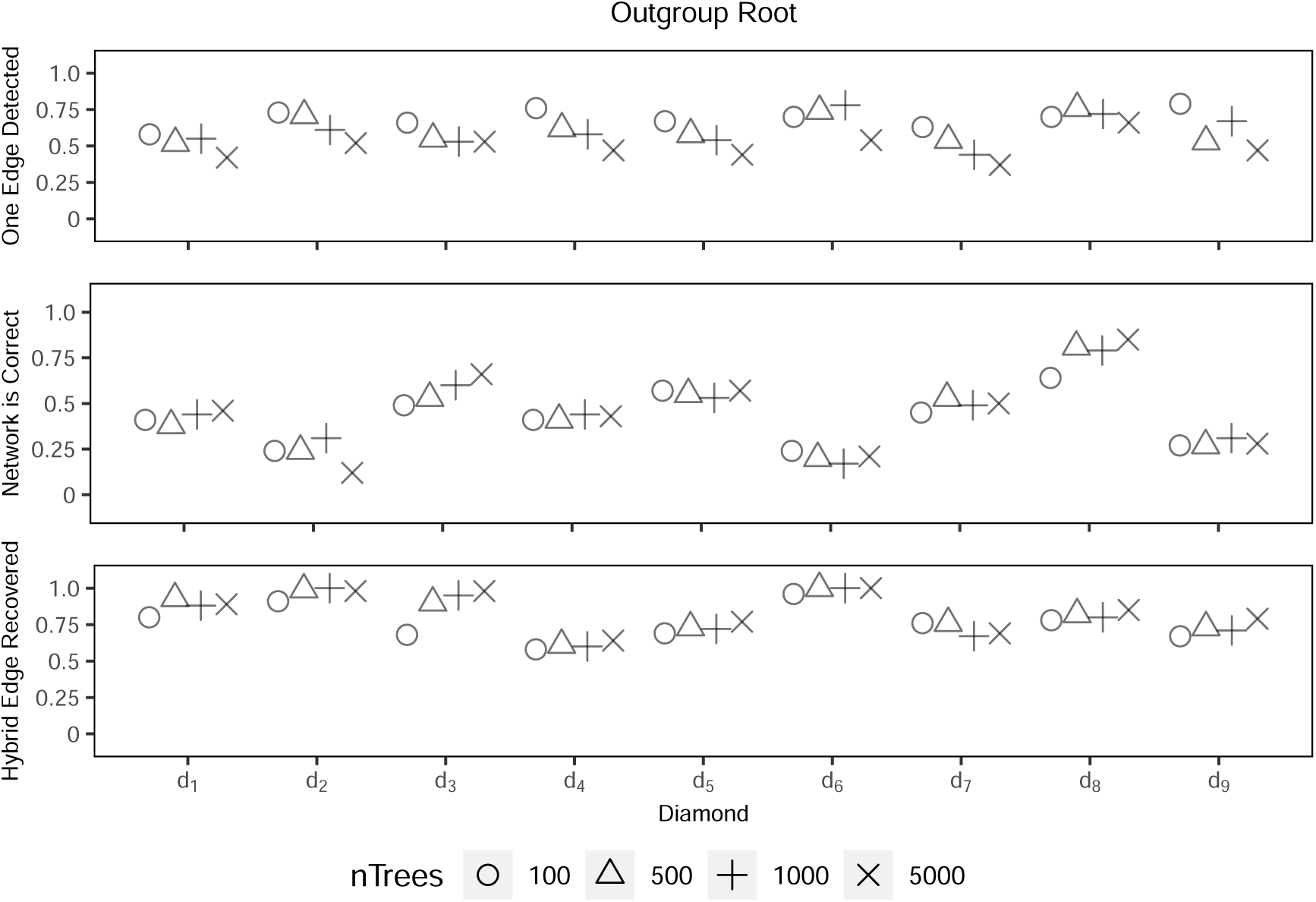
PhyloNet results for root 3 with random outgroups for 10% of gene trees.

**Figure 26:**
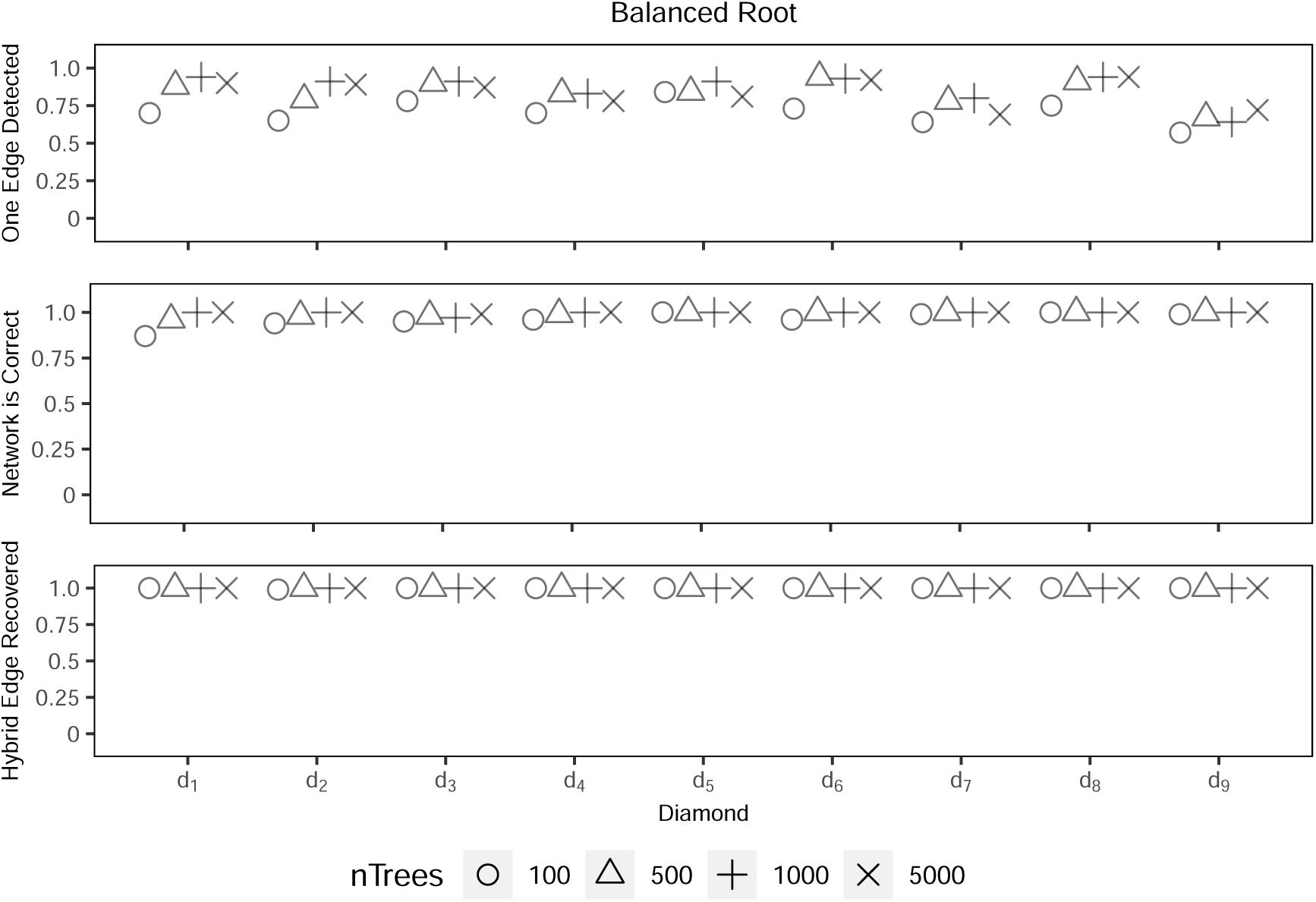
PhyloNet results for root 1 with random outgroups for 30% of gene trees.

**Figure 27:**
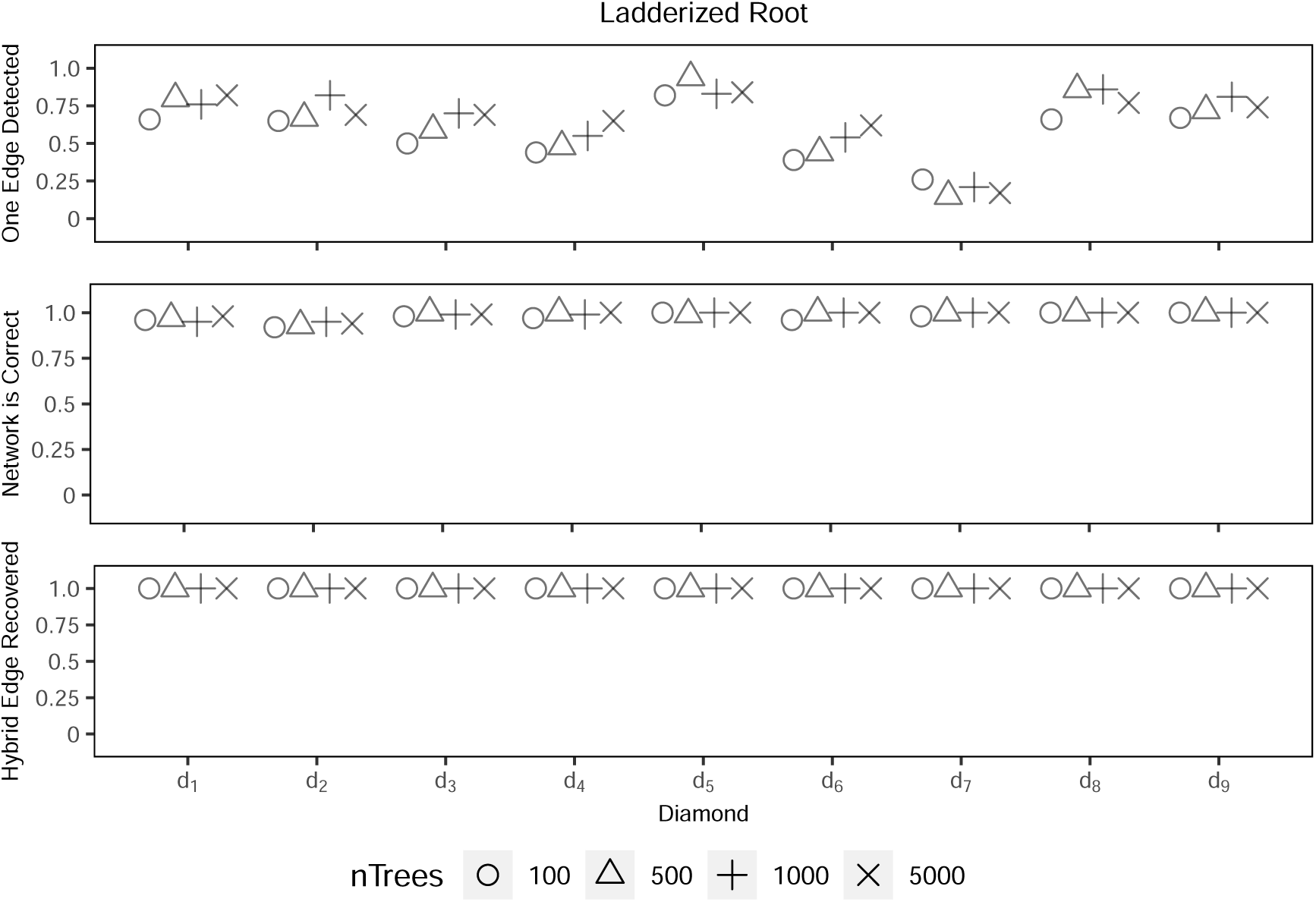
PhyloNet results for root 2 with random outgroups for 30% of gene trees.

**Figure 28:**
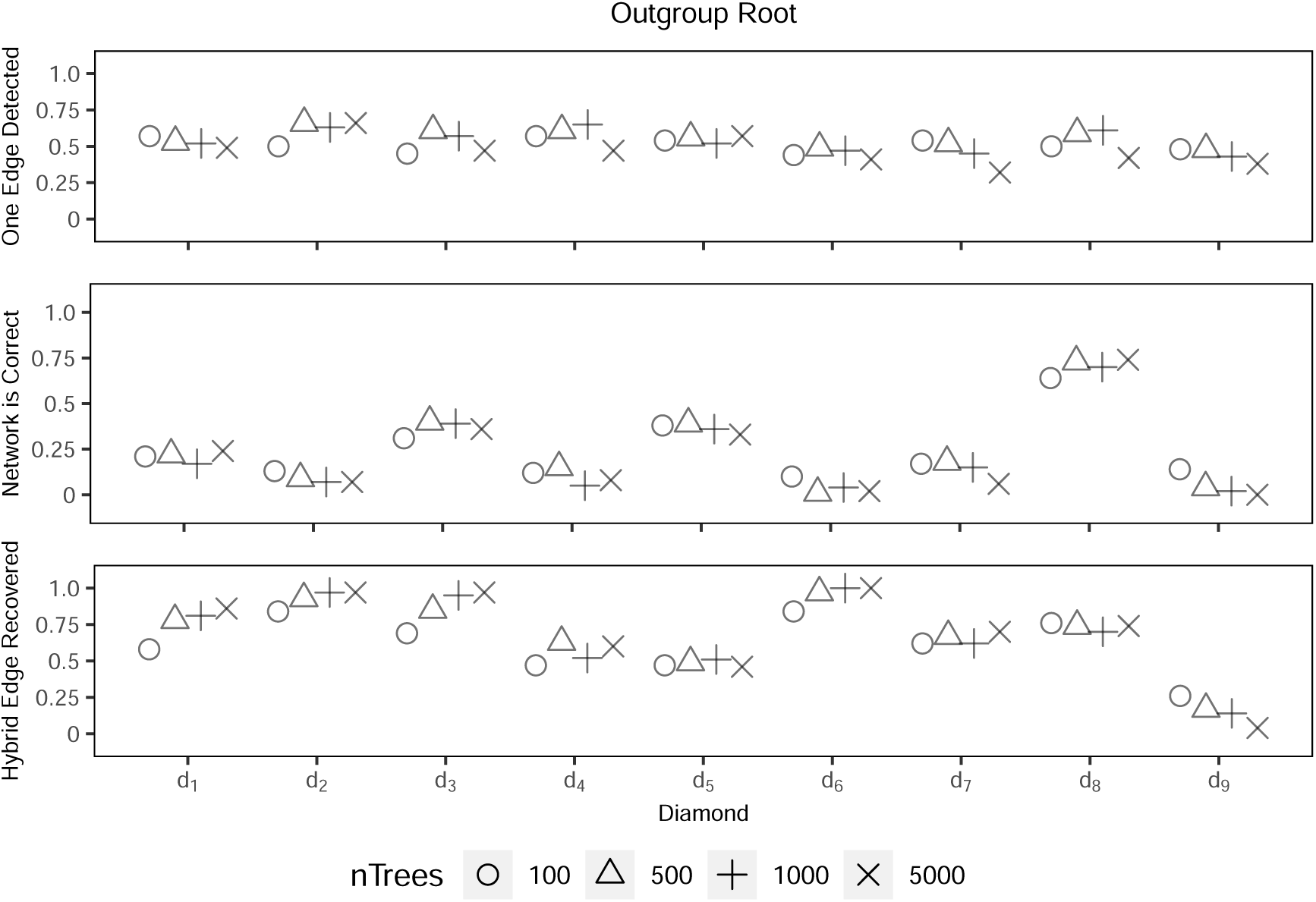
PhyloNet results for root 3 with random outgroups for 30% of gene trees.

**Figure 29:**
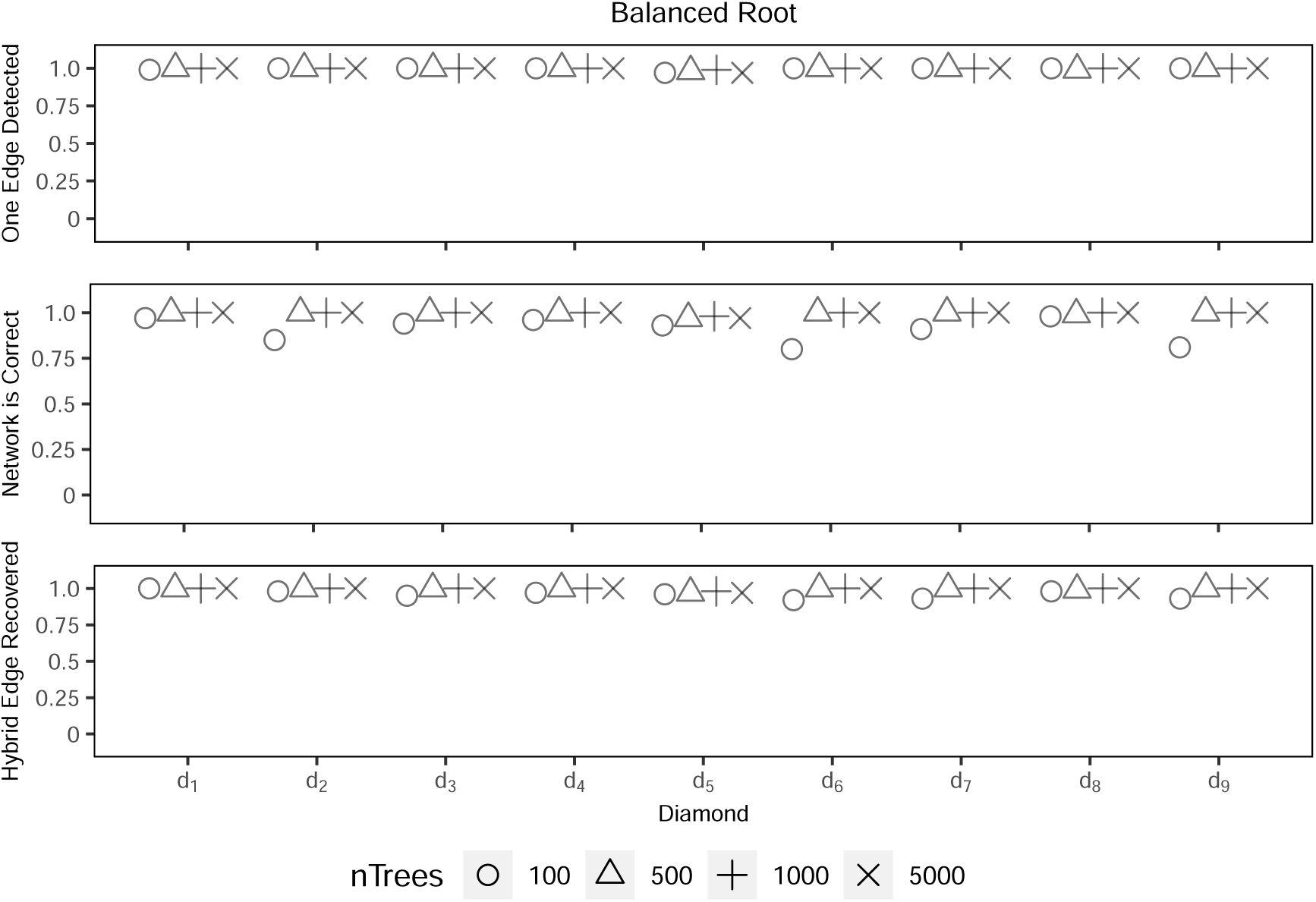
SNaQ results for root 1 with random outgroups and a random NNI move for 10% of gene trees.

**Figure 30:**
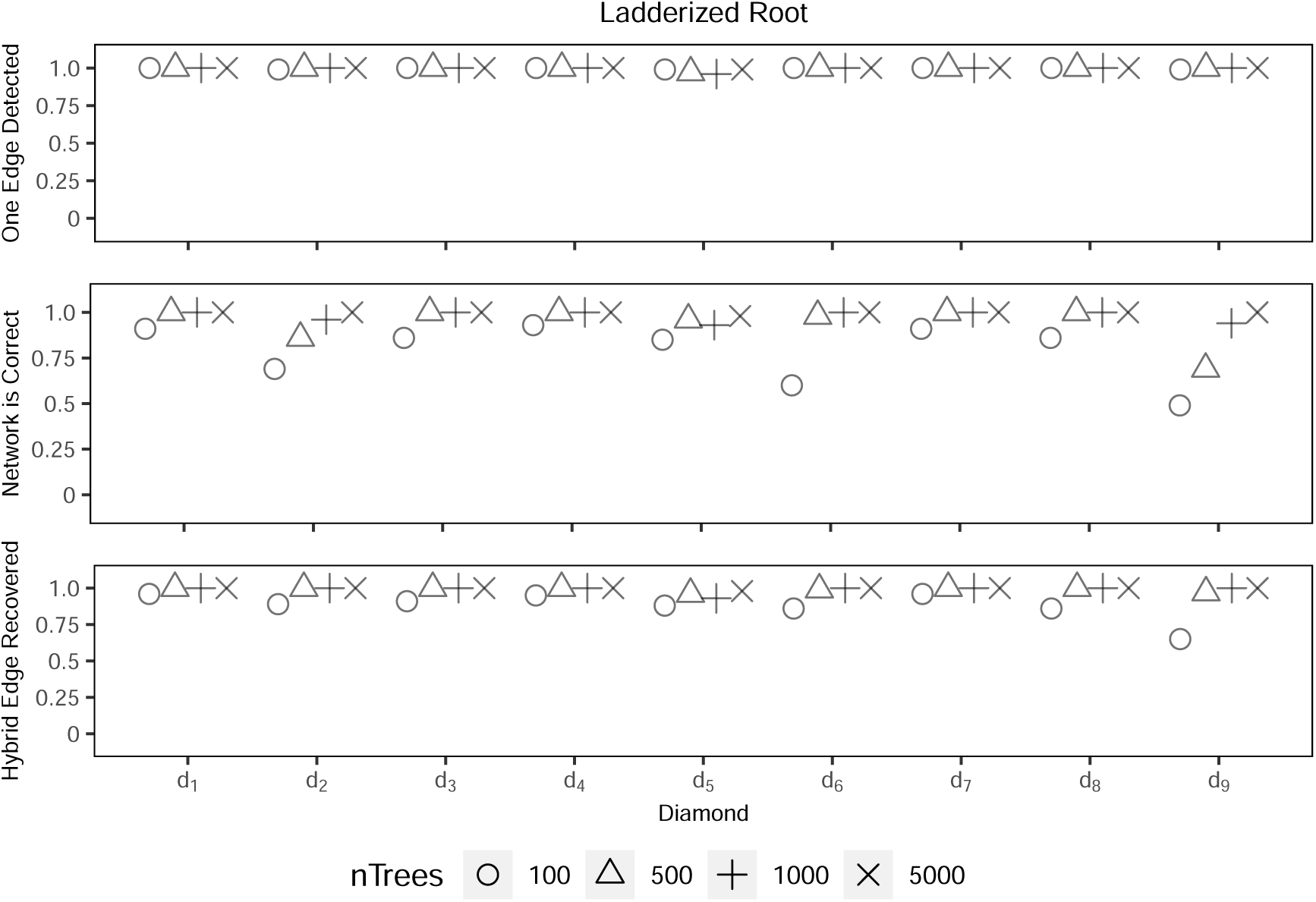
SNaQ results for root 2 with random outgroups and a random NNI move for 10% of gene trees.

**Figure 31:**
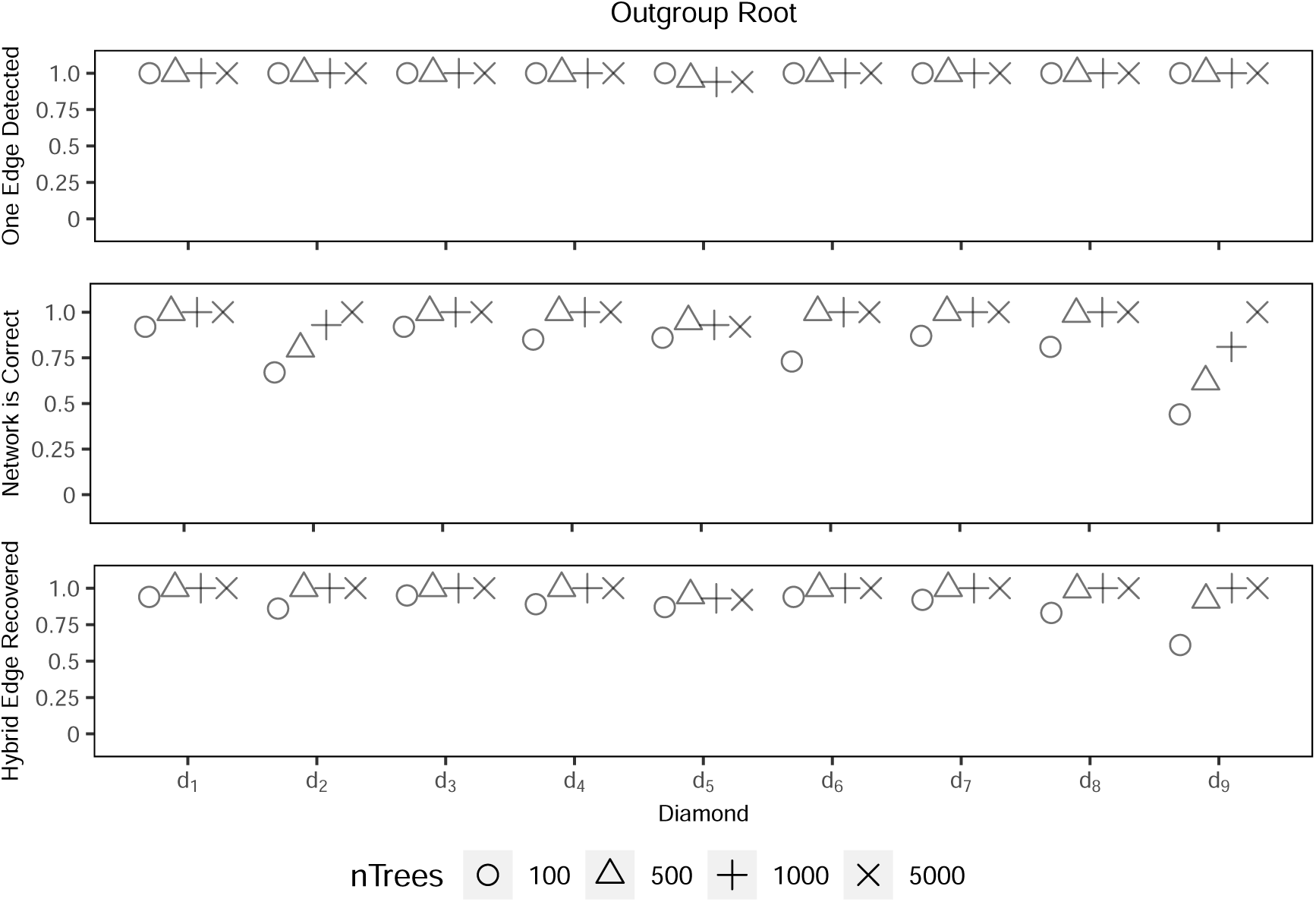
SNaQ results for root 3 with random outgroups and a random NNI move for 10% of gene trees.

**Figure 32:**
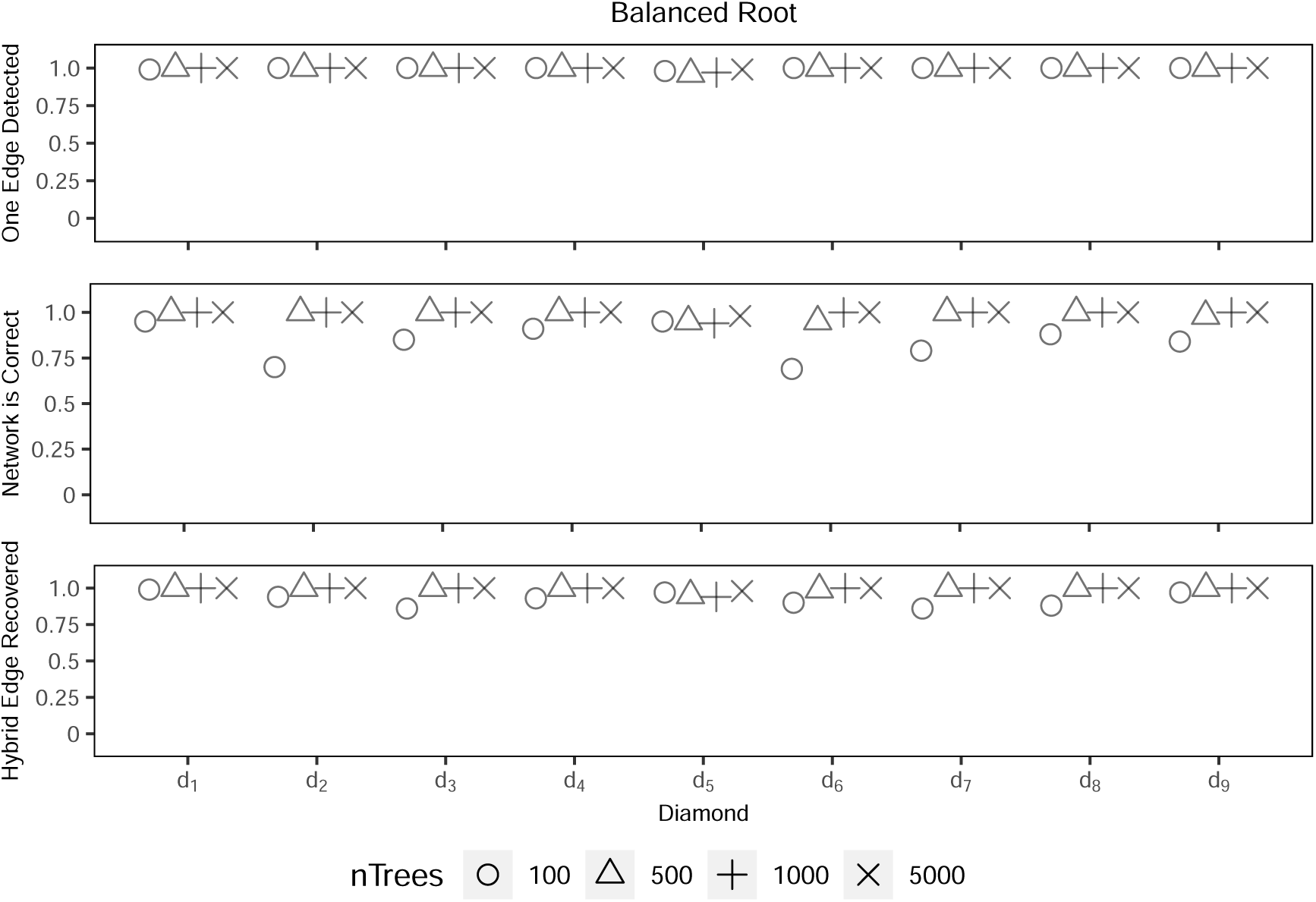
SNaQ results for root 1 with random outgroups and a random NNI move for 30% of gene trees.

**Figure 33:**
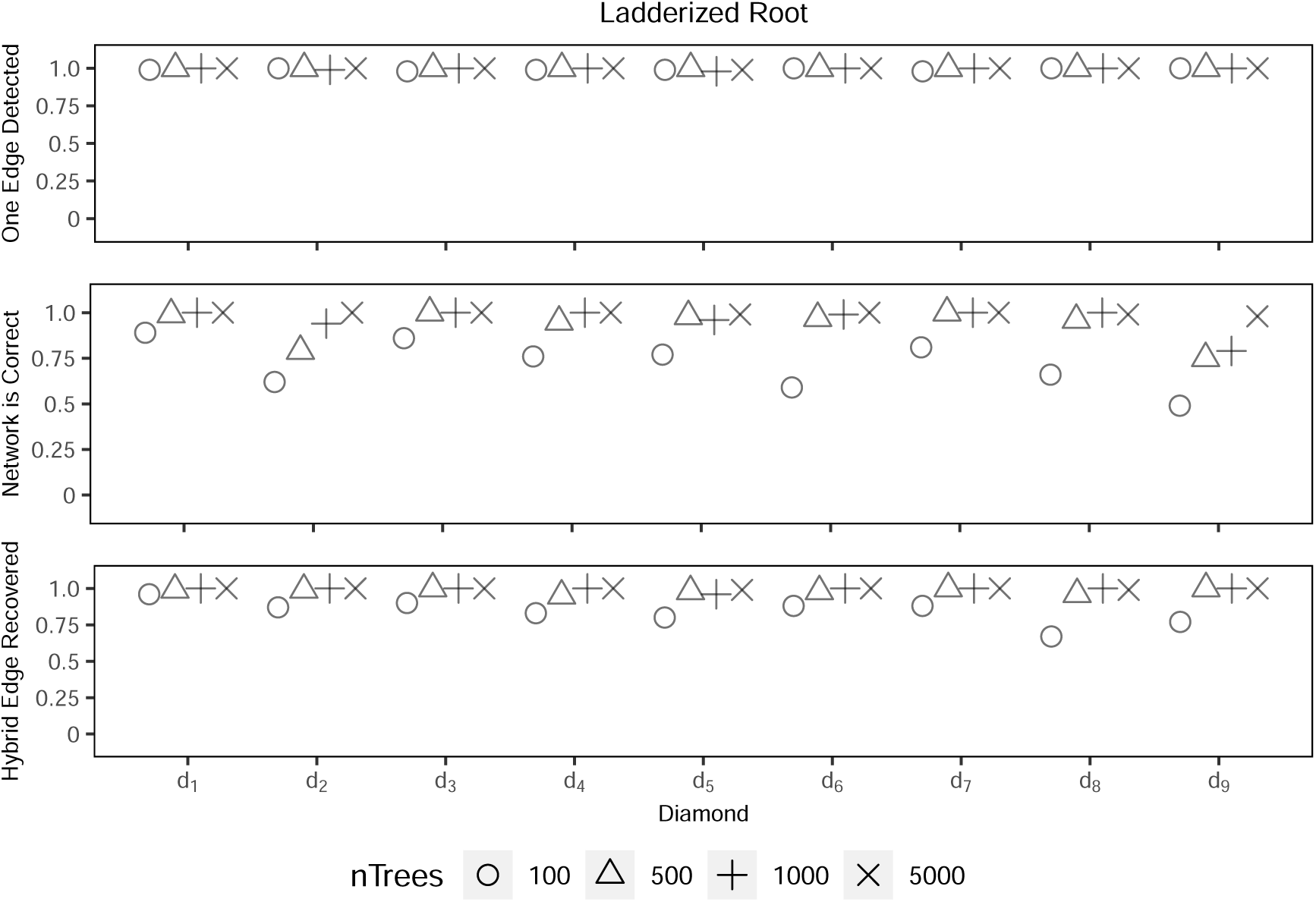
SNaQ results for root 2 with random outgroups and a random NNI move for 30% of gene trees.

**Figure 34:**
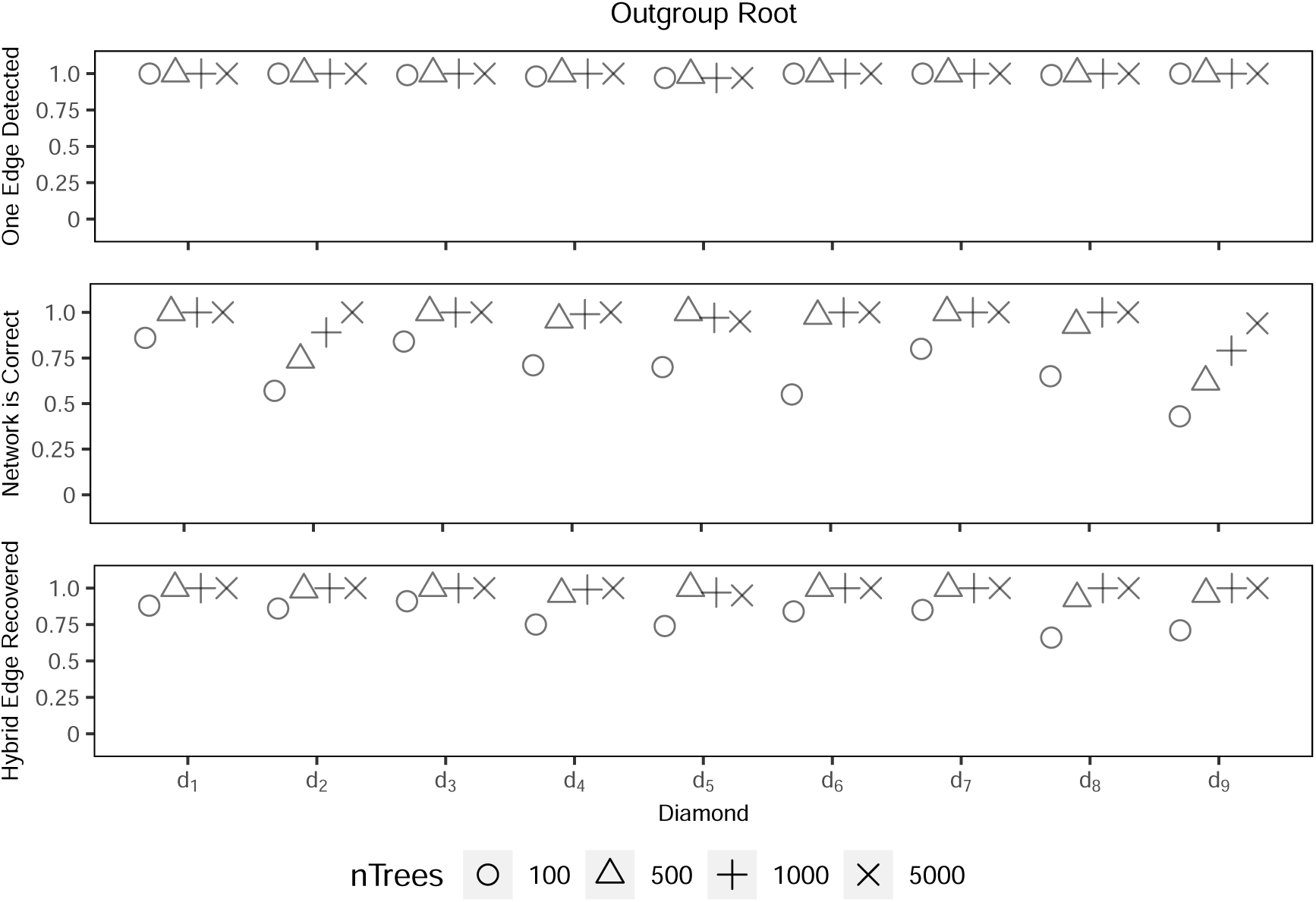
SNaQ results for root 3 with random outgroups and a random NNI move for 30% of gene trees.

**Figure 35:**
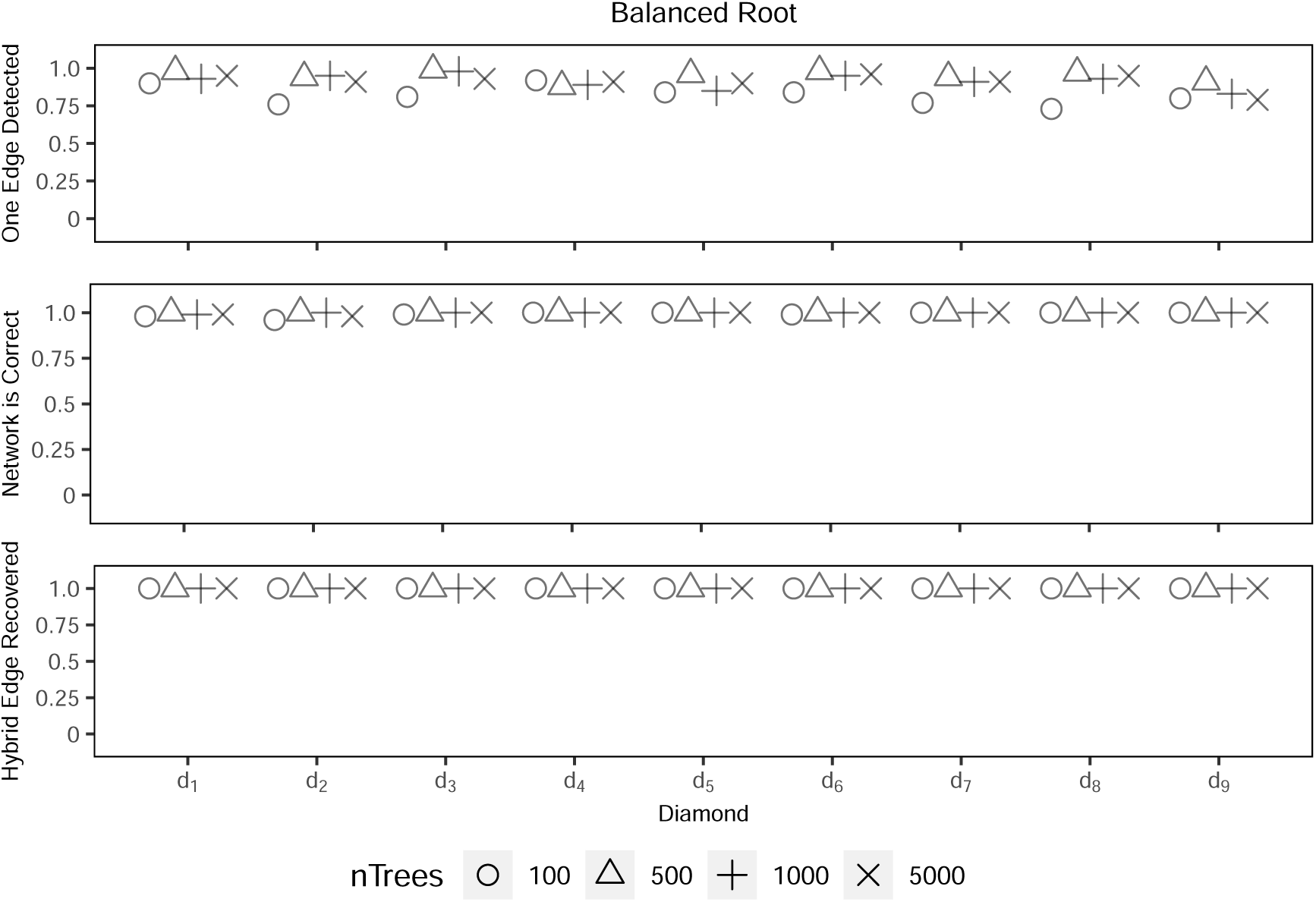
PhyloNet results for root 1 with random outgroups and a random NNI move for 10% of gene trees.

**Figure 36:**
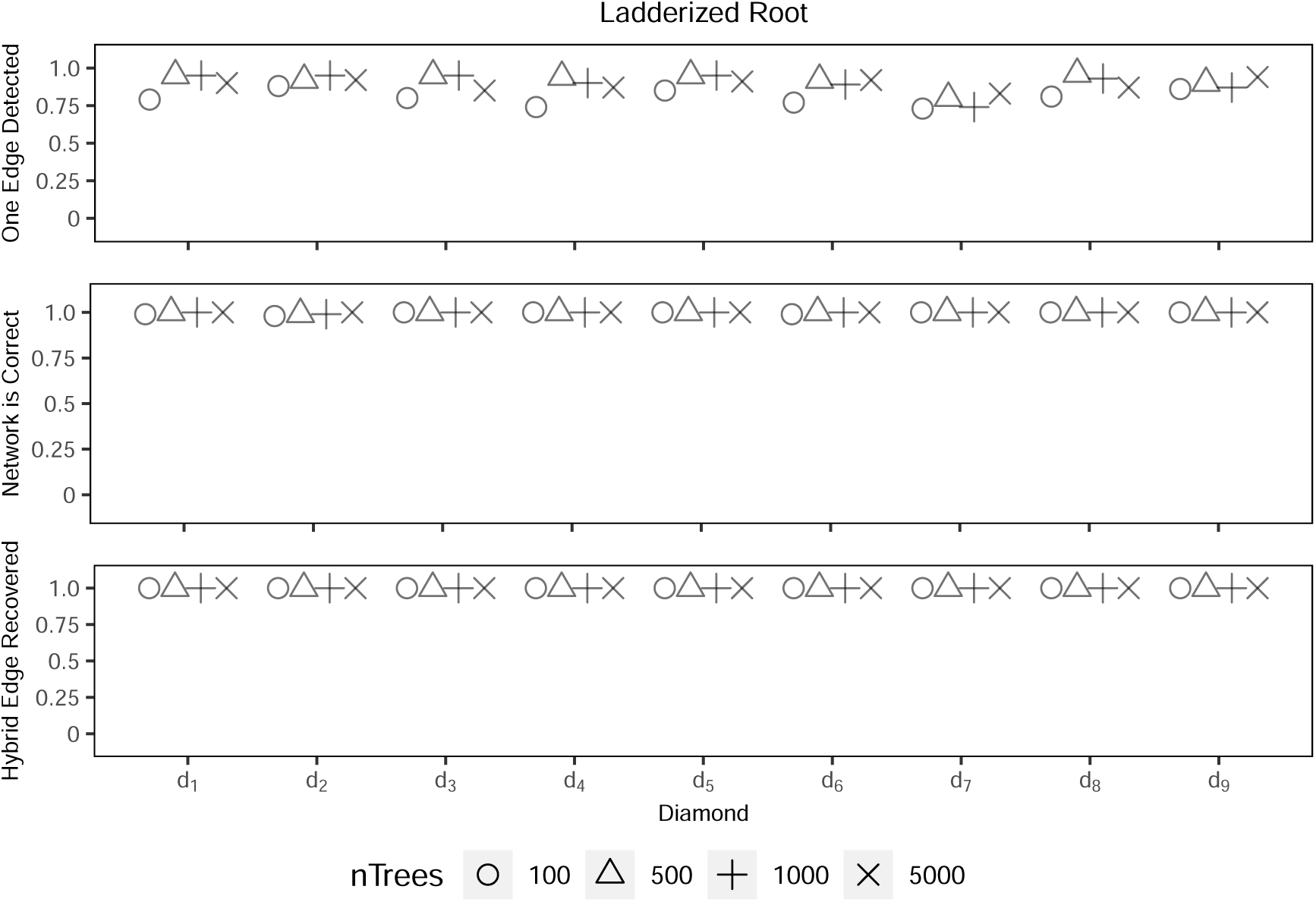
PhyloNet results for root 2 with random outgroups and a random NNI move for 10% of gene trees.

**Figure 37:**
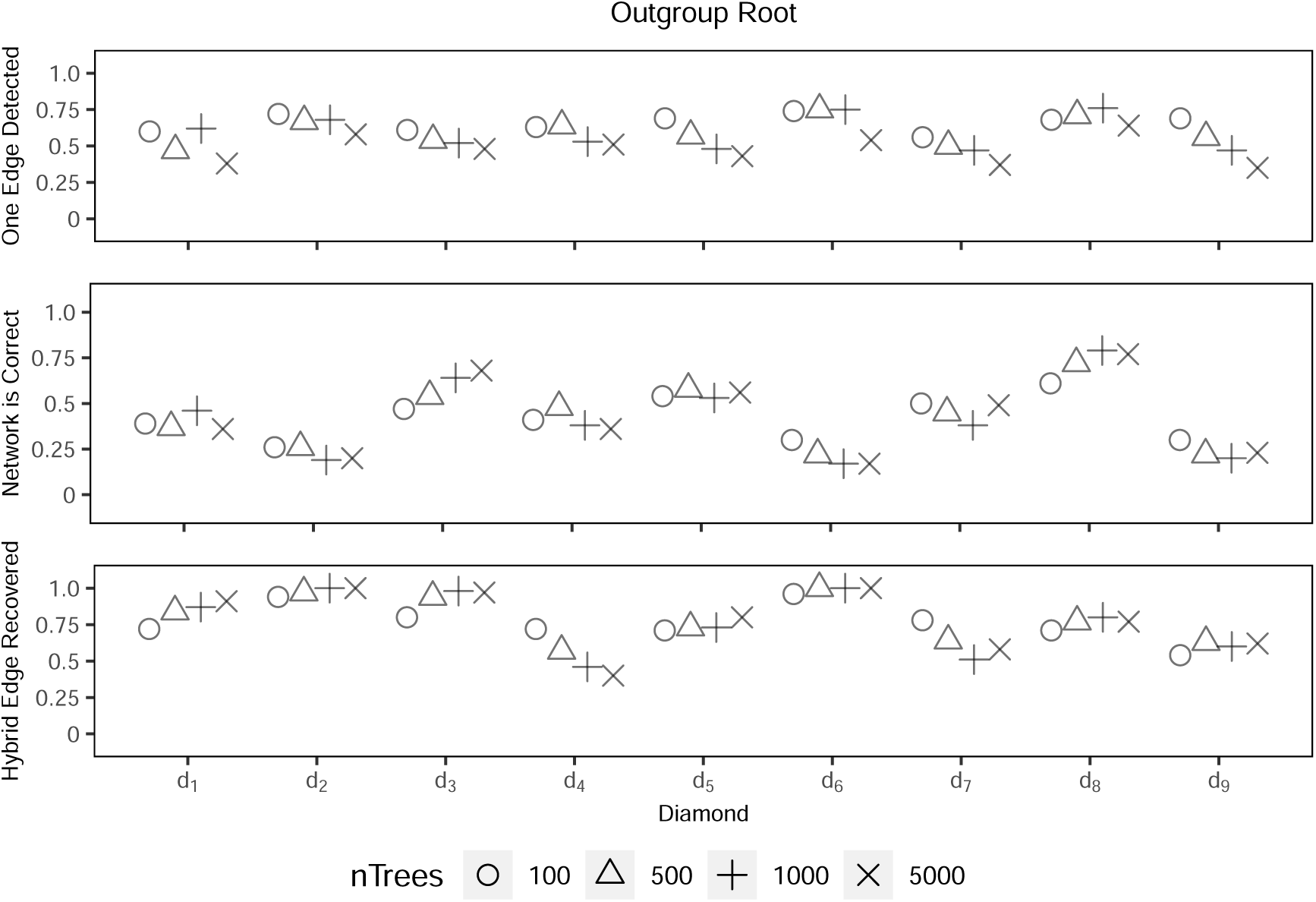
PhyloNet results for root 3 with random outgroups and a random NNI move for 10% of gene trees.

**Figure 38:**
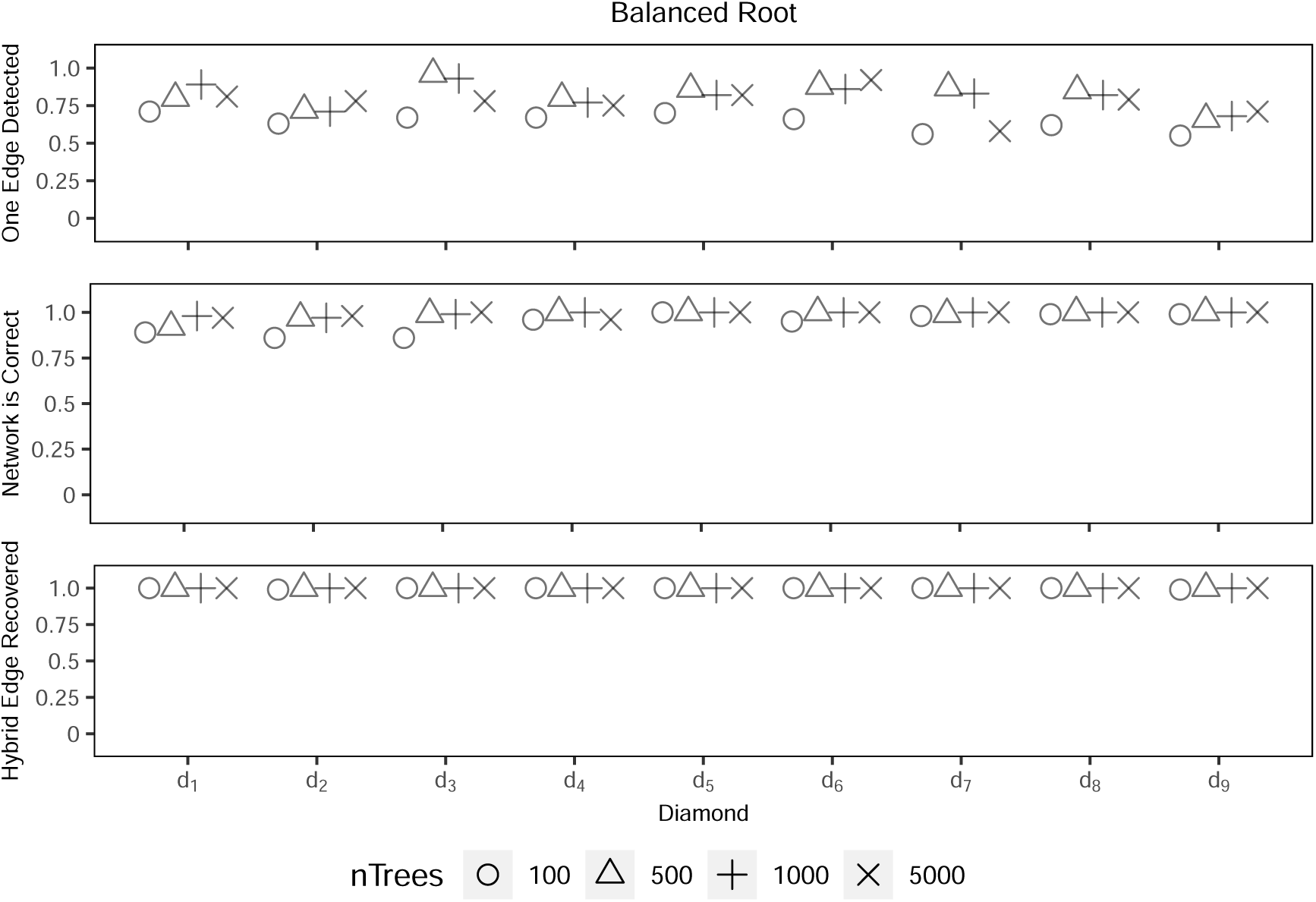
PhyloNet results for root 1 with random outgroups and a random NNI move for 30% of gene trees.

**Figure 39:**
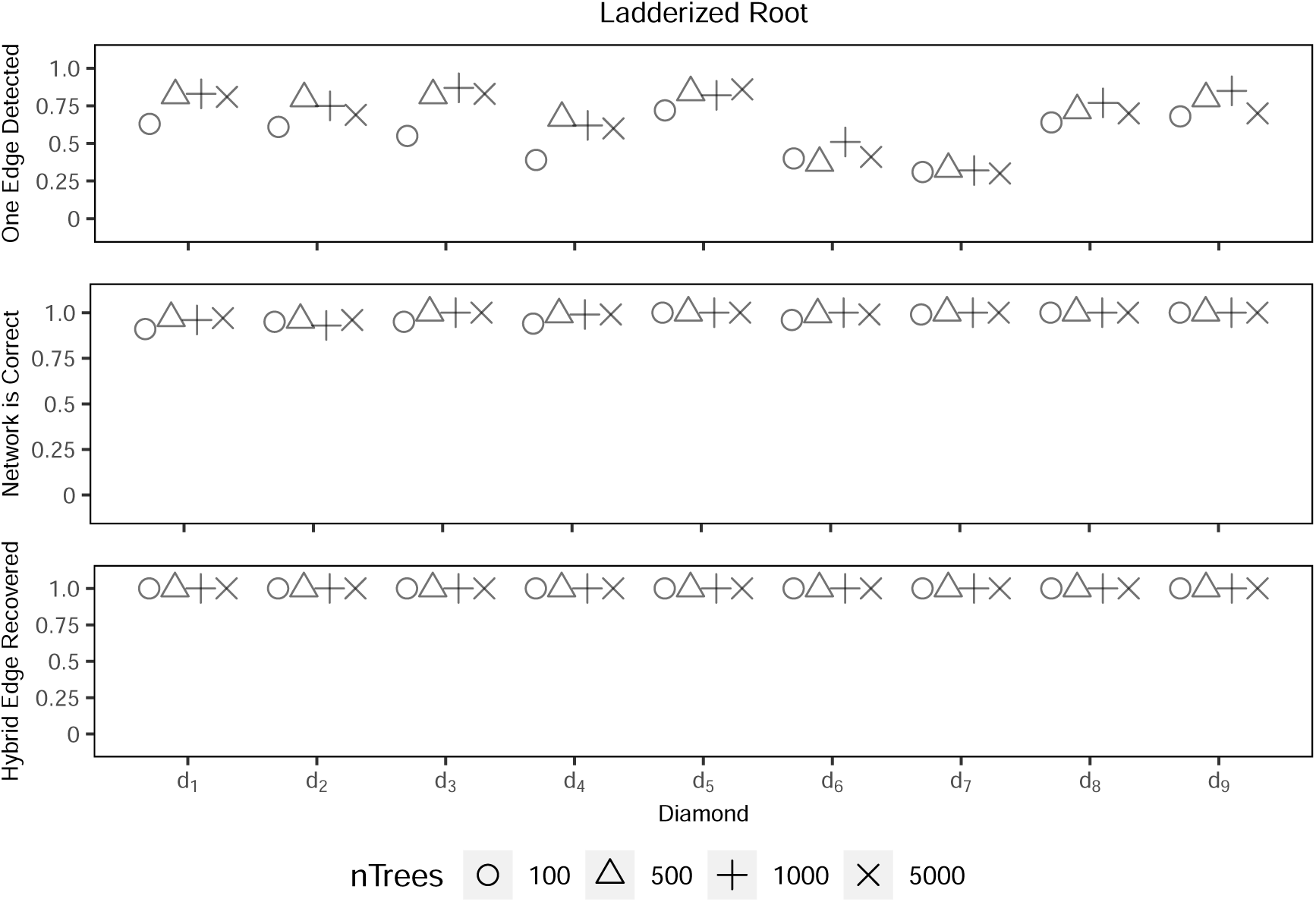
PhyloNet results for root 2 with random outgroups and a random NNI move for 30% of gene trees.

**Figure 40:**
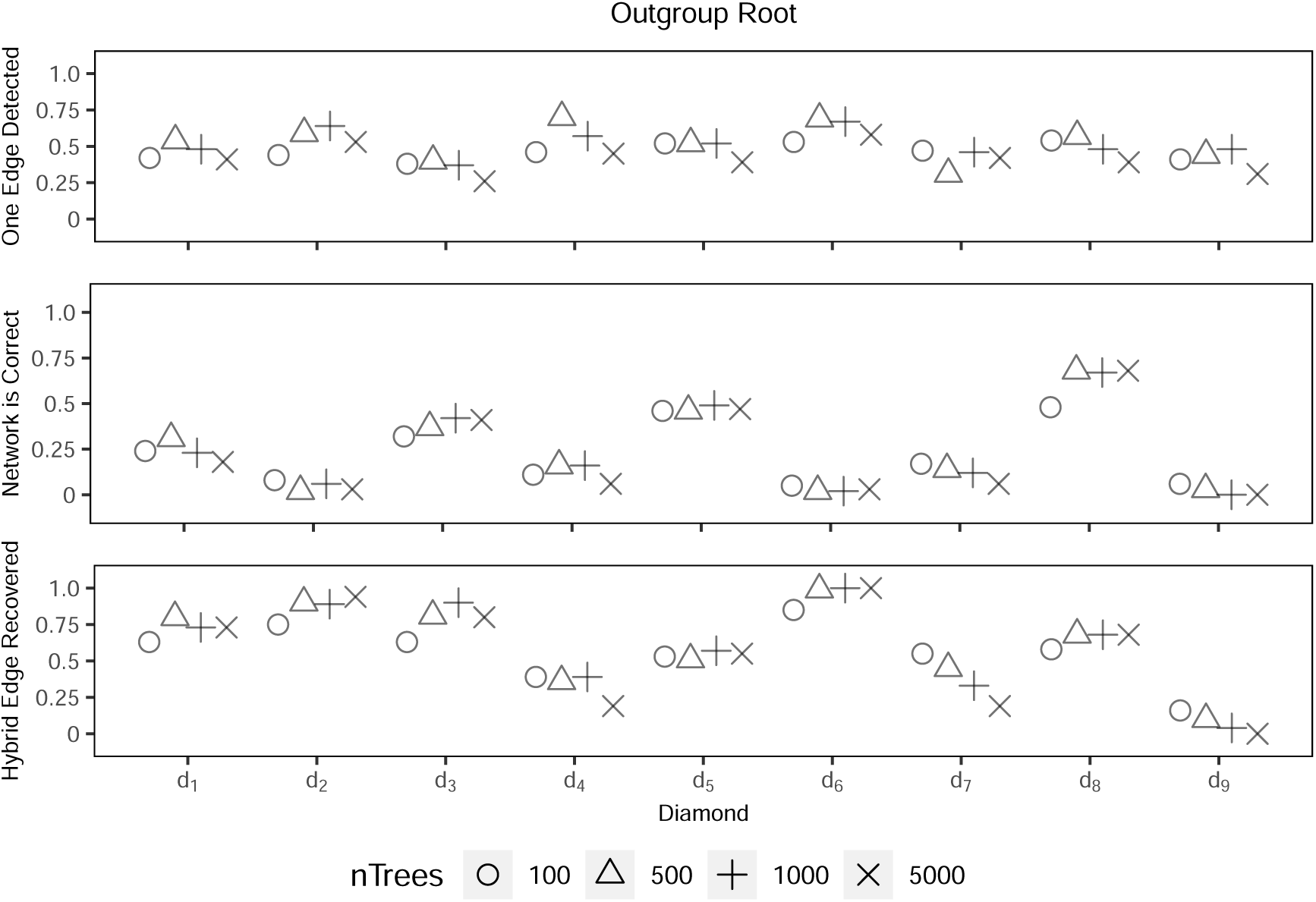
PhyloNet results for root 3 with random outgroups and a random NNI move for 30% of gene trees.

**Table 15:**
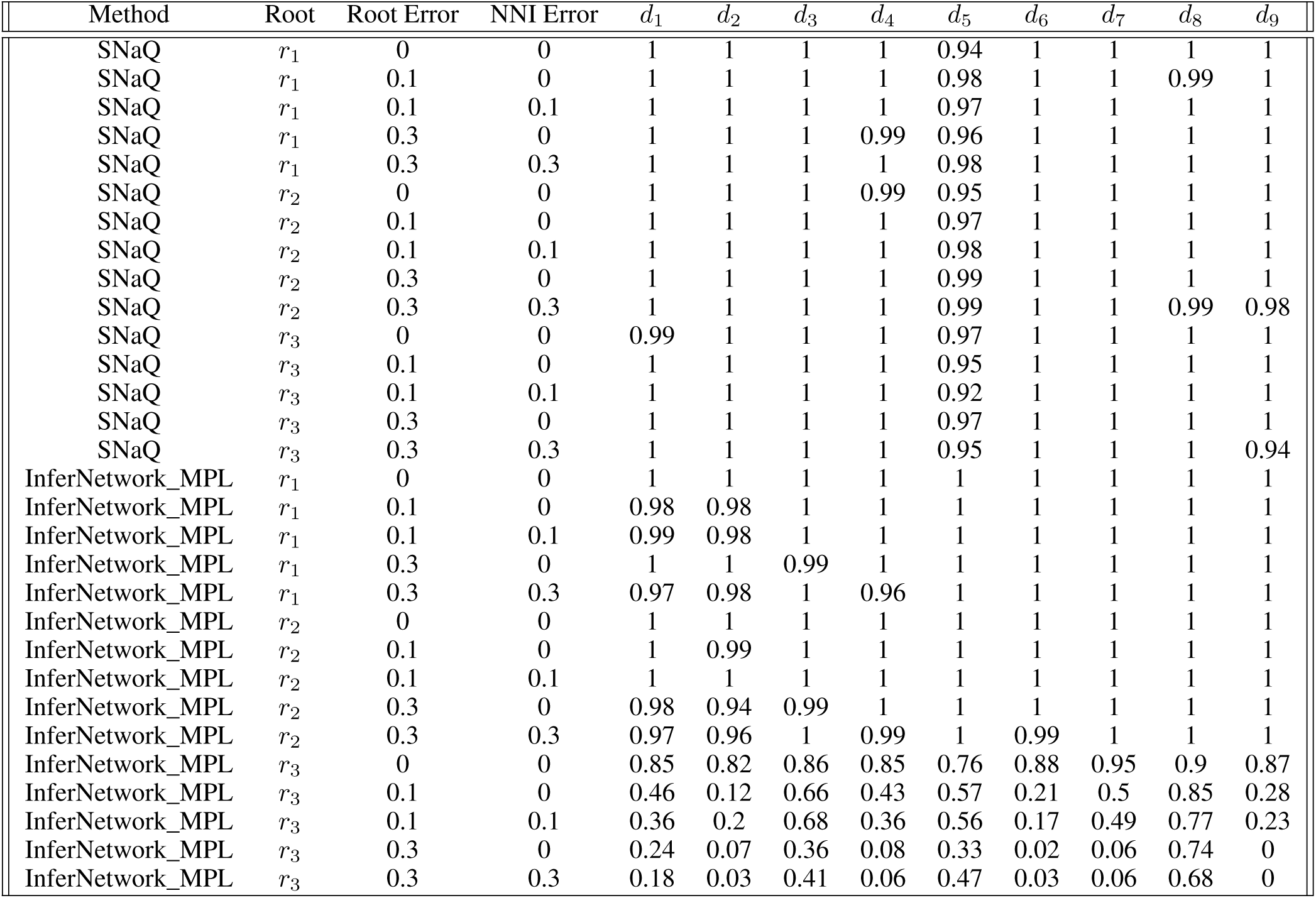
Proportion of correct networks recovered when using 5,000 gene trees. The Root Error and NNI Error are the proportion of gene trees affected by a random re-rooting or NNI move, respectively.

## References

[1] Hirotugu Akaike. A new look at the statistical model identification. IEEE transactions on automatic control, 19(6):716–723, 1974.

[2] Elizabeth S Allman, Hector Baños, Marina Garrote-Lopez, and John A Rhodes. Identifiability of level-1 species networks from gene tree quartets. ArXiv, 2024.

[3] Elizabeth S Allman, Hector Baños, and John A Rhodes. Nanuq: a method for inferring species networks from gene trees under the coalescent model. Algorithms for Molecular Biology, 14:1–25, 2019.

[4] Elizabeth S. Allman, James H. Degnan, and John A. Rhodes. Identifying the rooted species tree from the distribution of unrooted gene trees under the coalescent. Journal of Mathematical Biology, 62(6):833–862, 2011.

[5] Elizabeth S Allman, Sonia Petrovic, John A Rhodes, and Seth Sullivant. Identifiability of two-tree mixtures for group-based models. IEEE/ACM transactions on computational biology and bioinformatics, 8(3):710–722, 2010.

[6] Elizabeth S Allman and John A Rhodes. The identifiability of tree topology for phylogenetic models, including covarion and mixture models. Journal of Computational Biology, 13(5):1101–1113, 2006.

[7] Hector Baños. Identifying species network features from gene tree quartets under the coalescent model. Bulletin of mathematical biology, 81:494–534, 2019.

[8] Jean-Patrick Baudry, Cathy Maugis, and Bertrand Michel. Slope heuristics: overview and implementation. Statistics and Computing, 22(2):455–470, 2012.

[9] O. Chernomor D. Schrempf M.D. Woodhams A. von Haeseler R. Lanfear B.Q. Minh, H.A. Schmidt. Iq-tree 2: New models and efficient methods for phylogenetic inference in the genomic era. Molecular Biology and Evolution, 37(5):1530–1534, 2020.

[10] Ruoyi Cai and Cécile Ané. Assessing the fit of the multi-species network coalescent to multi-locus data. Bioinformatics, 37(5):634–641, 2021.

[11] Joseph T Chang. Full reconstruction of markov models on evolutionary trees: identifiability and consistency. Mathematical biosciences, 137(1):51–73, 1996.

[12] James H Degnan. Modeling hybridization under the network multispecies coalescent. Systematic biology, 67(5):786–799, 2018.

[13] Eric Y Durand, Nick Patterson, David Reich, and Montgomery Slatkin. Testing for ancient admixture between closely related populations. Molecular biology and evolution, 28(8):2239–2252, 2011.

[14] Tomáš Flouri, Xiyun Jiao, Bruce Rannala, and Ziheng Yang. A bayesian implementation of the multispecies coalescent model with introgression for phylogenomic analysis. Molecular biology and evolution, 37(4):1211–1223, 2020.

[15] DR Grayson and ME Stillman. Macaulay2, a software system for research in algebraic geometry. Available at http://www.math.uiuc.edu/Macaulay2/.

[16] Richard E Green, Johannes Krause, Adrian W Briggs, Tomislav Maricic, Udo Stenzel, Martin Kircher, Nick Patterson, Heng Li, Weiwei Zhai, Markus Hsi-Yang Fritz, et al. A draft sequence of the neandertal genome. science, 328(5979):710–722, 2010.

[17] Elizabeth Gross and Colby Long. Distinguishing phylogenetic networks. SIAM Journal on Applied Algebra and Geometry, 2(1):72–93, 2018.

[18] Elizabeth Gross, Leo van Iersel, Remie Janssen, Mark Jones, Colby Long, and Yukihiro Murakami. Distinguishing level-1 phylogenetic networks on the basis of data generated by markov processes. July 2020.

[19] Stefan Grünewald, Kristoffer Forslund, Andreas Dress, and Vincent Moulton. Qnet: an agglomerative method for the construction of phylogenetic networks from weighted quartets. Molecular Biology and Evolution, 24(2):532–538, 2007.

[20] Stefan Grunewald, Andreas Spillner, Sarah Bastkowski, Anja Bogershausen, and Vincent Moulton. Superq: computing supernetworks from quartets. IEEE/ACM transactions on computational biology and bioinformatics, 10(1):151–160, 2013.

[21] Masami Hasegawa, Hirohisa Kishino, and Taka-aki Yano. Dating of the human-ape splitting by a molecular clock of mitochondrial dna. Journal of molecular evolution, 22(2):160–174, 1985.

[22] BR Holland, D Penny, and MD Hendy. Outgroup misplacement and phylogenetic inaccuracy under a molecular clock—a simulation study. Systematic biology, 52(2):229–238, 2003.

[23] Katharina T. Huber, Vincent Moulton, Charles Semple, and Taoyang Wu. Quarnet inference rules for level-1 networks. Bulletin of Mathematical Biology, 80(8):2137–2153, 2018.

[24] Richard R Hudson. Testing the constant-rate neutral allele model with protein sequence data. Evolution, pages 203–217, 1983.

[25] John P Huelsenbeck, Jonathan P Bollback, and Amy M Levine. Inferring the root of a phylogenetic tree. Systematic biology, 51(1):32–43, 2002.

[26] Daniel H Huson and David Bryant. Application of phylogenetic networks in evolutionary studies. Molecular biology and evolution, 23(2):254–267, 2006.

[27] Daniel H Huson, Regula Rupp, and Celine Scornavacca. Phylogenetic networks: concepts, algorithms and applications. Cambridge University Press, 2010.

[28] Xiyun Jiao, Tomáš Flouri, and Ziheng Yang. Multispecies coalescent and its applications to infer species phylogenies and cross-species gene flow. National Science Review, 8(12):nwab127, 2021.

[29] Sungsik Kong, Joan Carles Pons, Laura Kubatko, and Kristina Wicke. Classes of explicit phylogenetic networks and their biological and mathematical significance. Journal of Mathematical Biology, 84(6):47, 2022.

[30] James H Leebens-Mack, Michael S Barker, Eric J Carpenter, Michael K Deyholos, Matthew A Gitzendanner, Sean W Graham, Zheng Grosse, Ivo and Li, Michael Melkonian, Siavash Mirarab, et al. One thousand plant transcriptomes and the phylogenomics of green plants. Nature, 574(7780):679–685, 2019.

[31] Zheng Li, George P Tiley, Sally R Galuska, Chris R Reardon, Thomas I Kidder, Rebecca J Rundell, and Michael S Barker. Multiple large-scale gene and genome duplications during the evolution of hexapods. Proceedings of the National Academy of Sciences, 115(18):4713–4718, 2018.

[32] Colby Long and Laura Kubatko. The effect of gene flow on coalescent-based species-tree inference. Systematic biology, 67(5):770–785, 2018.

[33] Sarah Lutteropp, Céline Scornavacca, Alexey M Kozlov, Benoit Morel, and Alexandros Stamatakis. Netrax: accurate and fast maximum likelihood phylogenetic network inference. Bioinformatics, 38(15):3725–3733, 2022.

[34] Wayne P Maddison. Gene trees in species trees. Systematic biology, 46(3):523–536, 1997.

[35] Alexey Markin, Tavis K Anderson, Venkata Sai Krishna Teja Vadali, and Oliver Eulenstein. Robinson-foulds reticulation networks. In Proceedings of the 10th ACM international conference on bioinformatics, computational biology and health informatics, pages 77–86, 2019.

[36] James Oldman, Taoyang Wu, Leo Van Iersel, and Vincent Moulton. Trilonet: piecing together small networks to reconstruct reticulate evolutionary histories. Molecular biology and evolution, 33(8):2151–2162, 2016.

[37] Fabio Pardi and Celine Scornavacca. Reconstructible phylogenetic networks: do not distinguish the indistinguishable. PLoS computational biology, 11(4):e1004135, 2015.

[38] James B Pease and Matthew W Hahn. Detection and polarization of introgression in a five-taxon phylogeny. Systematic biology, 64(4):651–662, 2015.

[39] Joseph Pickrell and Jonathan Pritchard. Inference of population splits and mixtures from genome-wide allele frequency data. Nature Precedings, pages 1–1, 2012.

[40] Charles-Elie Rabier, Vincent Berry, Marnus Stoltz, Joao D Santos, Wensheng Wang, Jean-Christophe Glaszmann, Fabio Pardi, and Celine Scornavacca. On the inference of complex phylogenetic networks by markov chain monte-carlo. PLoS Computational Biology, 17(9):e1008380, 2021.

[41] Bruce Rannala and Ziheng Yang. Bayes estimation of species divergence times and ancestral population sizes using dna sequences from multiple loci. Genetics, 164(4):1645–1656, 2003.

[42] John A Rhodes and Seth Sullivant. Identifiability of large phylogenetic mixture models. Bulletin of mathematical biology, 74:212–231, 2012.

[43] Francesc Rosselló and Gabriel Valiente. All that glisters is not galled. Mathematical biosciences, 221(1):54–59, 2009.

[44] Claudia Solís-Lemus and Cécile Ané. Inferring phylogenetic networks with maximum pseudolikelihood under incomplete lineage sorting. PLoS Genet., 12(3):e1005896, March 2016.

[45] Claudia Solís-Lemus, Paul Bastide, and Cécile Ané. Phylonetworks: a package for phylogenetic networks. Molecular biology and evolution, 34(12):3292–3298, 2017.

[46] Claudia Solis-Lemus, Arrigo Coen, and Cecile Ane. On the identifiability of phylogenetic networks under a pseudolikelihood model. October 2020.

[47] Ming Tan, Haixia Long, Bo Liao, Zhi Cao, Dawei Yuan, Geng Tian, Jujuan Zhuang, and Jialiang Yang. Qs-net: Reconstructing phylogenetic networks based on quartet and sextet. Frontiers in Genetics, 10:607, 2019.

[48] Dingqiao Wen, Yun Yu, Jiafan Zhu, and Luay Nakhleh. Inferring phylogenetic networks using phylonet. Systematic biology, 67(4):735–740, 2018.

[49] Matthieu Willems, Nadia Tahiri, and Vladimir Makarenkov. A new efficient algorithm for inferring explicit hybridization networks following the neighbor-joining principle. Journal of bioinformatics and computational biology, 12(05):1450024, 2014.

[50] Jingcheng Xu and Cécile Ané. Identifiability of local and global features of phylogenetic networks from average distances. Journal of Mathematical Biology, 86(1):12, 2023.

[51] Jialiang Yang, Stefan Grünewald, and Xiu-Feng Wan. Quartet-net: a quartet-based method to reconstruct phylogenetic networks. Molecular biology and evolution, 30(5):1206–1217, 2013.

[52] Ziheng Yang. Maximum likelihood phylogenetic estimation from dna sequences with variable rates over sites: approximate methods. Journal of Molecular evolution, 39(3):306–314, 1994.

[53] Yun Yu, James H. Degnan, and Luay Nakhleh. The probability of a gene tree topology within a phylogenetic network with applications to hybridization detection. PLoS genetics, 8(4):e1002660, jan 2012.

[54] Yun Yu, Jianrong Dong, Kevin J Liu, and Luay Nakhleh. Maximum Likelihood Inference of Reticulate Evolutionary Histories. PNAS, 111(46):16448–16453, 2014.

[55] Yun Yu and Luay Nakhleh. A maximum pseudo-likelihood approach for phylogenetic networks. BMC genomics, 16(10):1–10, 2015.

[56] Chi Zhang, Huw A Ogilvie, Alexei J Drummond, and Tanja Stadler. Bayesian inference of species networks from multilocus sequence data. Molecular biology and evolution, 35(2):504–517, 2018.

